# Caspase-11 drives macrophage hyperinflammation in models of Polg-related mitochondrial disease

**DOI:** 10.1101/2024.05.11.593693

**Authors:** Jordyn J. VanPortfliet, Yuanjiu Lei, Muthumeena Ramanathan, Camila Guerra Martinez, Jessica Wong, Tim J. Stodola, Brian R. Hoffmann, Kathryn Pflug, Raquel Sitcheran, Stephen C. Kneeland, Stephen A. Murray, Peter. J. McGuire, Carolyn L. Cannon, A. Phillip West

**Affiliations:** The Jackson Laboratory, Bar Harbor, Maine 04609, USA; Department of Microbial Pathogenesis and Immunology, College of Medicine, Texas A&M University, Bryan, Texas 77807, USA; Department of Pathology, Yale School of Medicine, New Haven, CT 06520, USA; Department of Cell Biology and Genetics, College of Medicine, Texas A&M University, Bryan, Texas 77807, USA; Metabolism, Infection and Immunity Section, National Human Genome Research Institute, National Institutes of Health, Bethesda, Maryland, 20892, USA

## Abstract

Mitochondrial diseases (MtD) represent a significant public health challenge due to their heterogenous clinical presentation, often severe and progressive symptoms, and lack of effective therapies. Environmental exposures, such bacterial and viral infection, can further compromise mitochondrial function and exacerbate the progression of MtD. Infections in MtD patients more frequently progress to sepsis, pneumonia, and other detrimental inflammatory endpoints. However, the underlying immune alterations that enhance immunopathology in MtD remain unclear, constituting a key gap in knowledge that complicates treatment and increases mortality in this vulnerable population. Here we employ in vitro and in vivo approaches to clarify the molecular and cellular basis for innate immune hyperactivity in models of polymerase gamma (Polg)-related MtD. We reveal that type I interferon (IFN-I)-mediated upregulation of caspase-11 and guanylate-binding proteins (GBPs) increase macrophage sensing of the opportunistic microbe *Pseudomonas aeruginosa* (PA) in *Polg* mutant mice. Furthermore, we show that excessive cytokine secretion and activation of pyroptotic cell death pathways contribute to lung inflammation and morbidity after infection with PA. Our work sheds new light on innate immune dysregulation in MtD and reveals potential targets for limiting infection- and inflammation-related complications in Polg-related MtD.

## Introduction

Mitochondrial diseases (MtD) are the most common inborn error of metabolism and are caused by deleterious variants in genes that function in oxidative phosphorylation (OXPHOS) and mitochondrial DNA (mtDNA) maintenance^1^. MtD can present at any age, vary in severity, and display clinical heterogeneity with differing degrees of organ system involvement^2,3^. Although the genetic basis of MtD is becoming clearer, the underlying mechanisms that cause pathology, as well as the contribution of environmental factors to disease progression, are less clear. Notably, viral and bacterial infections may unmask MtD or hasten the stepwise progression of disease by triggering metabolic decompensation, a dangerous deterioration in mitochondrial function^4^. Clinical evidence suggests that patients with MtD are highly susceptible to infection by opportunistic respiratory pathogens. Moreover, pneumonia and microbial sepsis are the primary causes of early death in pediatric patients, and some adults, with MtD^5,6^. Thus, there is ample evidence highlighting the damaging toll of infection in MtD^7^. However, the underlying immune alterations that drive immunopathology and prevent proper control of infection represent a significant knowledge gap that complicates treatment and enhances morbidity and mortality in this vulnerable patient population.

In addition to their bioenergetic functions, mitochondria are key metabolic and signaling hubs in the mammalian immune system^8^. Mitochondria are also important sources of damage-associated molecular patterns (DAMPs), such as mtDNA, which engage innate immune signaling when released from the organelle and are linked to inflammation in an increasing number of human diseases^9,10^. There is growing appreciation for a bidirectional relationship between MtD and inflammation, where mitochondrial dysfunction in MtD activates the innate immune system leading to inflammatory responses that further compromise mitochondrial activity^11^. In agreement with this hypothesis, elevated inflammatory cytokines have been observed in patients with Polymerase gamma (Polg)-related Alpers-Huttenlocher syndrome^12^, Barth syndrome^13^, and Friedreich ataxia^14,15^. Animal models have also revealed innate immune system dysregulation as a frequent feature of MtD. For example, the *Ndufs4^−/−^*mouse model of neurodegeneration and Leigh syndrome exhibits increased circulating pro-inflammatory cytokines along with macrophage and microglial activation^16–19^.

We recently uncovered that the *Polg^D257A^* mutator model of mtDNA instability exhibits a hyperinflammatory innate immune status due to chronic engagement of the cyclic GMP-AMP synthase (cGAS)—Stimulator of Interferon Genes (STING)—type one interferon (IFN-I) axis^20^. Notably, we reported that persistent IFN-I signaling in these mice causes widespread repression of the antioxidant and anti-inflammatory transcription factor nuclear factor erythroid 2-related factor 2 (Nrf2)^21,22^ in myeloid immune cells and organs of the *Polg^D257A^* mouse. Consequently, *Polg^D257A^*mice are more sensitive to septic shock caused by intraperitoneal injection of a high dose of lipopolysaccharide (LPS)^20^. Although *Polg^D257A^* mutator mice do not phenocopy any particular human MtD, these animals present pathology overlapping the MtD spectrum, including hearing loss, cardiomyopathy, progressive anemia, and lymphopenia^23–25^. Moreover, increased susceptibility to septic shock and elevated inflammatory and IFN-I signatures are common features of both *Polg* mutator mice and patients with diverse MtD^11^.

Overall, our prior work provided foundational evidence linking mitochondrial dysfunction to hyperinflammatory innate immune signaling in the *Polg^D257A^* model of MtD. However, there remains a critical need to expand these findings into relevant infection models, given the high rate of infections that frequently lead to innate immune hyperactivity and sepsis in MtD. *Pseudomonas aeruginosa* (PA) is a common opportunistic pathogen of concern in MtD, as infection can lead to sepsis and pneumonia^6^. PA is a robust activator of innate immunity in the respiratory tract, and lung colonization by PA drives persistent recruitment of monocytes and neutrophils leading to pneumonia, lung injury, and/or systemic bacterial spread^26,27^. PA is especially problematic in MtD due to the prevalence of antibiotic resistant strains, abundance in hospitals, and high rate of recurrence^26,28,29^. Thus, it is important to define how PA and other opportunistic bacteria engages the innate immune system in MtD to define immunomodulatory strategies that boost anti-bacterial immunity, while limiting hyperinflammation that may hasten metabolic decompensation and disease progression.

In this study, we utilized in vitro and in vivo PA infection models to characterize the underlying innate immune mechanisms governing hyperinflammation in *Polg^D257A^* mice. We also report a new mouse model that incorporates a pathogenic *Polg* variant observed in humans. We reveal that both *Polg* mutant models exhibit excessive macrophage inflammasome activation, cytokine secretion, and pyroptotic cell death, which contribute to inflammation, acute lung injury, and morbidity after infection with PA. These findings further advance knowledge of how mitochondrial dysfunction impacts innate immune responses during bacterial infection and Polg-related diseases and reveals innate immune targets for limiting infection- and inflammation-related complications in MtD.

## Results

### *Polg^D257A^* mutant macrophages exhibit increased IFN-I and pro-inflammatory responses after bacterial challenge

Our recent work uncovered that elevated IFN-I signaling augments innate immune activity and inflammation in *Polg^D257A^* mice challenged with Toll-like receptor (TLR) agonist, LPS^20^. We hypothesized that the hyperinflammatory phenotype of *Polg^D257A^* mice might be further exacerbated during bacterial infection. Therefore, we challenged bone marrow-derived macrophages (BMDMs) with *Pseudomonas aeruginosa* strain O1 (PAO1) (Fig. 1a) and performed RNA sequencing (RNAseq) and pathway analysis (Fig. 1b). Similar to BMDMs treated with LPS^20^, PAO1-challenged *Polg^D257A^* BMDMs displayed heightened expression of pro-inflammatory cytokines, IFN-I, interferon-stimulated genes (ISGs), and innate immune signaling factors (Fig. 1c-d; Supplementary Fig. 1a). Gene Set Enrichment Analysis confirmed PAO1-infected *Polg^D257A^*BMDMs exhibit more robust pro-inflammatory cytokine, chemokine, and IFN-I gene expression relative to wild-type (WT) BMDMs (Supplementary Fig. 1b). Additional experiments substantiated RNAseq datasets, revealing heightened expression of ISGs after challenge of *Polg^D257A^* BMDMs with PAO1 between six to twenty four hours (Fig. 1e-f). Proteomic analysis of PAO1-challenged BMDMs revealed strong concurrence with RNAseq data, showing elevated IFN-I and inflammatory responses in *Polg^D257A^*BMDMs (Fig. 1g-h, Supplementary Fig. 1c). Moreover, multi-analyte cytokine analysis by LegendPlex showed a global elevation of proinflammatory cytokines produced by *Polg^D257A^* peritoneal macrophages (PerMacs) six hours after PAO1 challenge (Fig. 1i) and ELISA showed a more rapid and sustained release of these cytokines over a one-day time course (Fig. 1j). In sum, our data highlight that *Polg^D257A^* macrophages are hyperactivated by both LPS and PAO1 challenge.

**Fig. 1:**
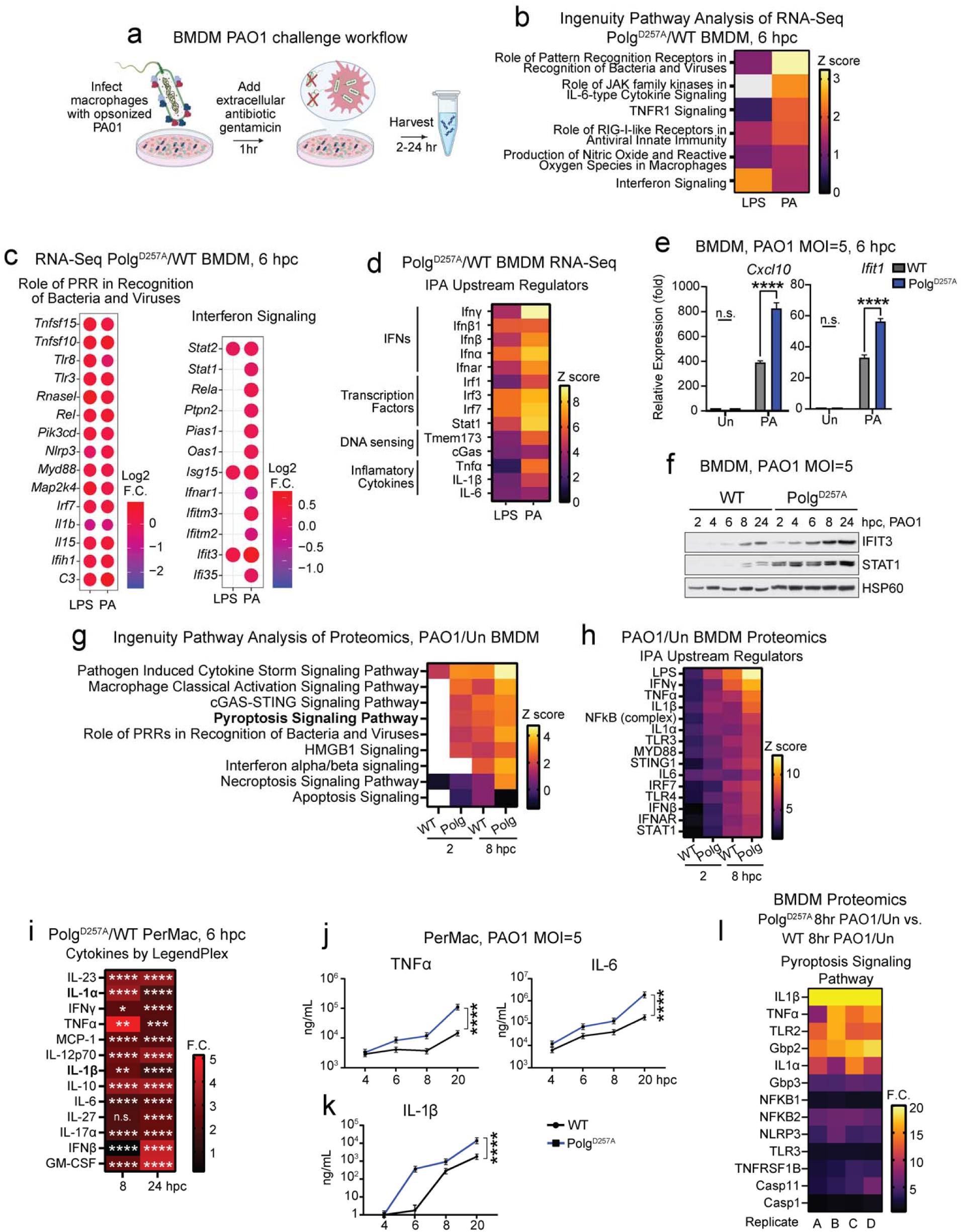
*Polg^D257A^*mutator macrophages exhibit increased IFN-I and pro-inflammatory responses after challenge with LPS or *Pseudomonas aeruginosa*. **(a)** Workflow for preparing RNA from *Polg* mutant BMDMs after *Pseudomonas aeruginosa* (PA) strain O1 (PAO1) challenge. **(b)** Ingenuity Pathway Analysis (IPA) of RNA-seq from bone marrow-derived macrophages (BMDMs) showing Z-scores of pathways in *Polg^D257A^* homozygous mutant BMDMs compared to wild-type (WT) after LPS or PAO1 challenge. hpc=hours post challenge. N=3 biological replicates/genotype/condition. **(c)** Log^2^ fold changes of representative genes in IPA pathways from (**b**) in *Polg^D257A^*BMDMs compared to WT after LPS or PAO1. **(d)** Protein abundance of interferon-stimulated genes (ISGs) in BMDMs exposed to PAO1 at a multiplicity of infection (MOI) of 5 at the indicated hours post challenge (hpc). **(e)** qRT-PCR analysis of ISGs in WT and *Polg^D257A^* BMDMs after PAO1. N=3 biological replicates, 2 technical replicates/genotype/condition. **(f)** Western blot analysis of interferon stimulated genes (ISGs) over time course infection. **(g)** Ingenuity pathway analysis showing z-scores of upregulated pathways by proteomics from *Polg^D257A^* (PAO1/Un)/WT (PAO1/Un) BMDM. N=4 biological replicates **(h)** IPA Upstream Regulator analysis showing Z-scores in *Polg^D257A^* (PAO1/Un)/WT (PAO1/Un). **(i)** Legendplex pro-inflammatory cytokines analysis of PerMac supernatants 24 hours post challenge (hpc). N=3 biological replicates **(j)** Proinflammatory cytokines in supernatant of peritoneal macrophages (PerMacs) measured by ELISA. MOI= multiplicity of infection, N=2 biological replicates per time point. **(k)** IL-1β in supernatant of peritoneal macrophages (PerMacs) measured by ELISA. MOI= multiplicity of infection, N=2 biological replicates per time point. **(l)** Ingenuity pathway analysis of proteomics from *Polg^D257A^* (PAO1, 8hpc/Un)/WT (PAO1, 8 hpc/Un) BMDM proteins in the Pyroptotic Signaling Pathway. N=4 biological replicates Statistics: (e, i & j) Two-Way ANOVA. *P<0.05, **P<0.01, ***P<0.001, ****P<0.0001 Error bars represent SEM.

### *Polg^D257A^* mutant macrophages exhibit potentiated pyroptotic cell death upon bacterial challenge

As we observed increased IL-1β secretion from *Polg^D257A^* PerMacs (Fig. 1i, k), we next investigated inflammatory cell death pathways during PAO1 infection. RNAseq and pathway analysis revealed significant enrichment of genes involved in inflammasome signaling, immunogenic cell death, and death receptor signaling in *Polg^D257A^* BMDMs after PAO1 challenge (Supplementary Fig. 1d-e). Pyroptotic cell death proteins were also highly upregulated in proteomic analysis (Fig. 1l). To further confirm a role for pyroptosis in hyperinflammation and increased cell death in *Polg^D257A^* macrophages, we incubated BMDMs with Ac-YVAD-cmk (AcYVAD), a selective caspase-1 inhibitor. AcYVAD nearly abrogated IL-1β secretion from *Polg^D257A^*macrophages, while also reducing the levels of inflammasome-independent inflammatory cytokines TNFα and IL-6 upon PAO1 challenge (Fig. 2a). In contrast, QVD-OPH, a more potent inhibitor of apoptotic caspases, had minor effects on PAO1-induced cytokine secretion in *Polg^D257A^* BMDMs (Fig. 2a). Moreover, we did not observe striking differences in necroptotic signaling proteins during PAO1 challenge, and necrostatin-1 failed to blunt elevated cytokine secretion from *Polg^D257A^* BMDMs (Supplementary Fig. 2a-b).

**Fig. 2:**
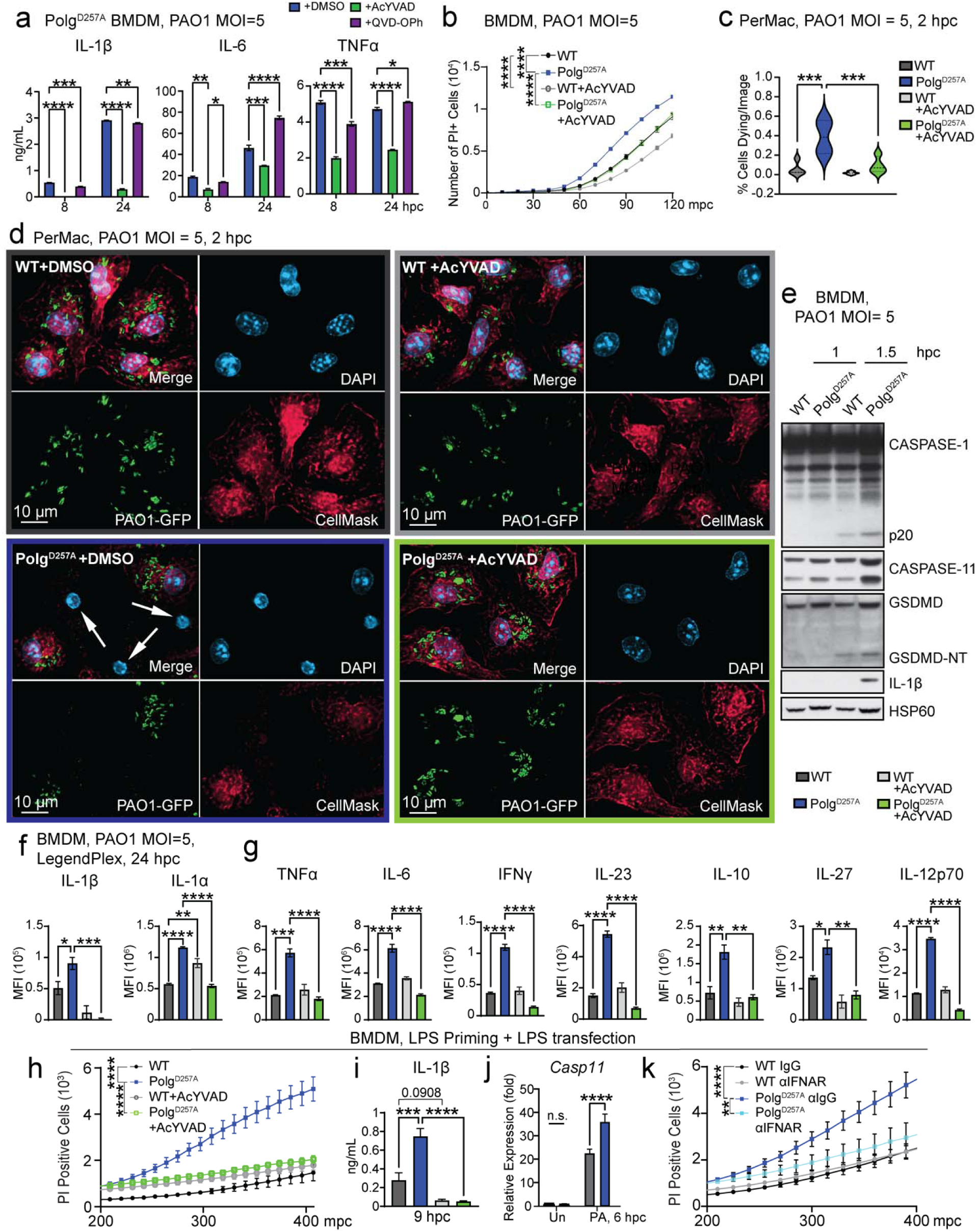
Caspase-11 and caspase-1 drive pyroptosis and inflammation in *Polg^D257A^* macrophages challenged with PAO1. **(a)** ELISA analysis of inflammatory cytokines in BMDM supernatant after PAO1 or PAO1 plus caspase inhibitors. N= 2 biological replicates/genotype/condition/timepoint. **(b)** Cytation5 image analysis of PI positive BMDMs after PAO1 or PAO1 plus caspase-1 inhibitor. N= 6 biological replicates/genotype/condition. **(c)** Percent PerMacs dying after PAO1 and PAO1 plus caspase-1 inhibitor from IF **(d)**. N=4 IF images/genotype/condition. **(d)** Representative IF of PerMacs after PAO1 or PAO1 plus caspase inhibitor. Cells were stained with DAPI to mark the nucleus, CellMask to mark the total cell volume, and anti-GFP to mark GFP tagged PAO1. White arrows highlight dying cells characterized by condensed nuclei and loss of CellMask staining. **(e)** Western blot analysis of lysates from BMDMs challenged with PAO1. **(f)** LegendPlex IL-1 cytokine analysis of BMDM supernatant 24 hpc. All undefined comparisons between WT and WT+AcYVAD are not significant. N=3 biological replicates/genotype/condition. **(g)** LegendPlex pro-inflammatory cytokine analysis of BMDM supernatant 24 hpc. All undefined comparisons between WT and WT+AcYVAD are not significant. N=3 biological replicates/genotype/condition. **(h)** Cytation5 image quantification of PI positive BMDMs after LPS priming plus LPS transfection and LPS priming plus LPS transfection plus caspase-1 inhibitor. N= 3-5 biological replicates/genotype/condition. **(i)** ELISA analysis of IL-1β in BMDM supernatant after LPS priming plus LPS transfection in the presence or absence of caspase-1 inhibitor. N=4 biological replicates **(j)** qRT-PCR analysis of Caspase-11 RNA expression in BMDMs after PAO1. N=3 biological replicates, 3 technical replicates/genotype/condition. **(k)** Cytation5 image quantification of PI positive cells in BMDMs after LPS priming plus LPS transfection and incubated with αIFNAR blocking antibody or αIgG control. N= 4-6 biological replicates/genotype/condition. Statistics: (c, f, g & i) One-Way ANOVA (a, b, h, j & k) Two-Way ANOVA. *P<0.05, **P<0.01, ***P<0.001, ****P<0.0001 Error bars represent SEM.

Consistent with cell death gene expression patterns, *Polg^D257A^*BMDMs more rapidly became propidium iodide (PI) positive during PAO1 challenge (Fig. 2b). *Polg^D257A^* PerMacs stained weakly for the nuclear and plasma membrane dye CellMask HCS Deep Red and exhibited rapid condensation of nuclei after PAO1 challenge (Fig. 2c-d). Reduced CellMask staining was also observed in ASC positive WT BMDMs after LPS priming and nigericin treatment (Supplementary Fig. 2c-d), indicating that low CellMask staining marks cells undergoing pyroptosis. *Polg^D257^*^A^ BMDMs also released more lactate dehydrogenase (LDH) after PAO1 challenge (Supplementary Fig. 2e). We also noted elevated release of PAO1 into the culture supernatant from *Polg^D257A^*BMDMs, concomitant with lower numbers of intracellular bacteria (Supplementary Fig. 3a). Reduced intracellular bacteria was not the result of impaired phagocytosis, as *Polg^D257A^* BMDMs displayed no impairment in uptake of LPS coated fluorescent microspheres (Supplementary Fig. 3b-c). AcYVAD treatment was sufficient to blunt pyroptotic phenotypes of *Polg^D257A^* macrophages (Fig. 2b-d) after incubation with PAO1, further supporting a caspase-1-dependent cell death pathway.

In addition to activating NLRP3, PA can also activate the NLRC4 canonical inflammasome and the non-canonical caspase-11 inflammasome^27,30–32^. Culture of *Polg^D257A^* BMDMs with non-pathogenic *E. coli* (DH5α), which lack flagellin and therefore can only activate NLRP3 or caspase-11, also promoted pyroptotic cell death (Supplementary Fig. 3d-e) and increased IL-1β secretion (Supplementary Fig. 3f), while boosting TNFα and IL-6 secretion (Supplementary Fig. 3g). Collectively, these data indicate that elevated cell death and cytokine secretion in *Polg^D257A^* macrophages during bacterial challenge are not due to PAO1 virulence factors but are likely caused by potentiated activation of NLRP3 and/or caspase-11 inflammasomes.

### Elevated IFN-I enhances expression and activation of caspase-11 and caspase-1 in *Polg^D257A^* macrophages

In WT macrophages, PAO1 mainly engages the NLRC4 inflammasome, while activation of NLRP3 and caspase-11 inflammasomes occur at later stages of infection^30,33–35^. Given elevated pyroptosis in PAO1-infected *Polg^D257A^* BMDMs, we next performed isolated challenge experiments to evaluate hyperactivation of particular inflammasome pathways in *Polg^D257A^* macrophages. *Polg^D257A^* BMDMs treated with LPS alone did not exhibit increased PI uptake, suggesting that priming is not sufficient to potentiate cell death (Supplementary Fig. 4a). Surprisingly, LPS priming followed by flagellin transfection to activate the NLRC4 inflammasome revealed that *Polg^D257A^* macrophages exhibit reduced PI uptake and IL-1β secretion (Supplementary Fig. 4b-c). LPS priming followed by the addition of ATP also revealed modestly impaired NLRP3 inflammasome activation in *Polg^D257A^*BMDMs, as measured by PI uptake, IL-1β secretion, and caspase-1 cleavage (Supplementary Fig. 4d-f). LPS priming followed by nigericin treatment revealed similar impairments in NLRP3 inflammasome activity (Supplementary Fig. 3g-i). OXPHOS has recently been shown to be vital for NLRP3 activation in response to nigericin and other stimuli^36^. *Polg^D257A^* BMDMs displayed dysfunctional mitochondrial morphology by TEM as well as reduced basal and maximal respiration at rest and after LPS challenge (Supplementary Fig. 4j-m), suggesting that reduced canonical NLRP3 activation is likely due to OXPHOS impairment in *Polg^D257A^* BMDMs.

Although *Polg^D257A^* BMDMs exhibited reduced NLRP3 and NLRC4 responses when triggered with pure inflammasome activators, mutant BMDMs displayed elevated caspase-1 and Gasdermin D (GSDMD) cleavage, as well as increased caspase-11 and IL-1β expression, after incubation with PAO1 as compared with WT controls (Fig. 2e). Moreover, AcYVAD treatment inhibited the hypersecretion of IL-1β and other pro-inflammatory cytokines in *Polg^D257A^* BMDMs challenged with PAO1 for 24 hours (Fig. 2f-g). Notably, IL-1α secretion was markedly higher in *Polg^D257A^* macrophages after PAO1 challenge (Fig. 1i and 2f). IL-1α is an alarmin that feeds forward to augment production of other cytokines, and the robust secretion of IL-1α from myeloid cells involves caspase-11^37–39^. Thus, given elevated caspase-11 expression in *Polg^D257A^* BMDMs, we next explored its involvement in cell death and cytokine secretion. After LPS priming followed by LPS transfection, *Polg^D257A^* BMDMs rapidly became PI positive and secreted increased levels of IL-1β (Fig. 2h-i). Both phenotypes could be markedly inhibited by treatment with AcYVAD, indicating that heightened activation of caspase-11 feeds forward to activate caspase-1 more robustly in *Polg^D257A^* BMDMs (Fig. 2h-i). Inhibition of NLRP3 with MCC950 reduced cell death and hyperinflammation in *Polg^D257A^* BMDMs after PAO1 challenge (Supplementary Fig. 5a-d). NLRP3-dependent caspase-1 activation can drive GSDMD to permeabilize mitochondrial membranes^40^. Therefore, we isolated mitochondria from unchallenged and PAO1-challenged BMDMs and examined GSDMD accumulation in mitochondria-enriched fractions. Although there was no difference in the mitochondrial accumulation of GSDMD between genotypes, we observed markedly increased GSDMD cleavage in whole cell lysates of *Polg^D257A^* BMDMs rapidly upon PAO1 challenge (Supplementary Fig. 5e).

As caspase-11 activation requires IFN-I priming^41^ and *Polg^D257A^* macrophages and mice exhibit elevated IFN-I signaling^20^, we reasoned that IFN-I is responsible for caspase-11 expression and activation in *Polg^D257A^* BMDMs. In agreement, we observed potentiated caspase-11 mRNA and protein after PAO1 infection (Fig. 1l, 2e, 2j). Notably, IFN alpha receptor (IFNAR) blockade using the MAR1-5A3 monoclonal antibody markedly reduced PI uptake after caspase-11 activation by transfected LPS (Fig. 2k). Collectively, these data indicate that the IFN-I-dependent potentiation of caspase-11 drives non-canonical caspase-1 activation and hyperinflammation in *Polg^D257A^* macrophages exposed to PAO1.

### Elevated IFN-I signaling increases GBP expression and inflammation in *Polg^D257A^* macrophages

We next hypothesized that elevated IFN-I may contribute to the hyperinflammatory response of *Polg^D257A^* macrophages during PAO1 infection. Priming macrophages with an anti-IFNAR antibody prior to PAO1 challenge reduced cell death and IL-1β release in *Polg^D257A^* BMDMs (Fig. 3a-c). This blockade also reduced production of NF-κΒ-dependent cytokines and ISG expression at the RNA level over a day time course (Fig 3d-e).

**Fig. 3:**
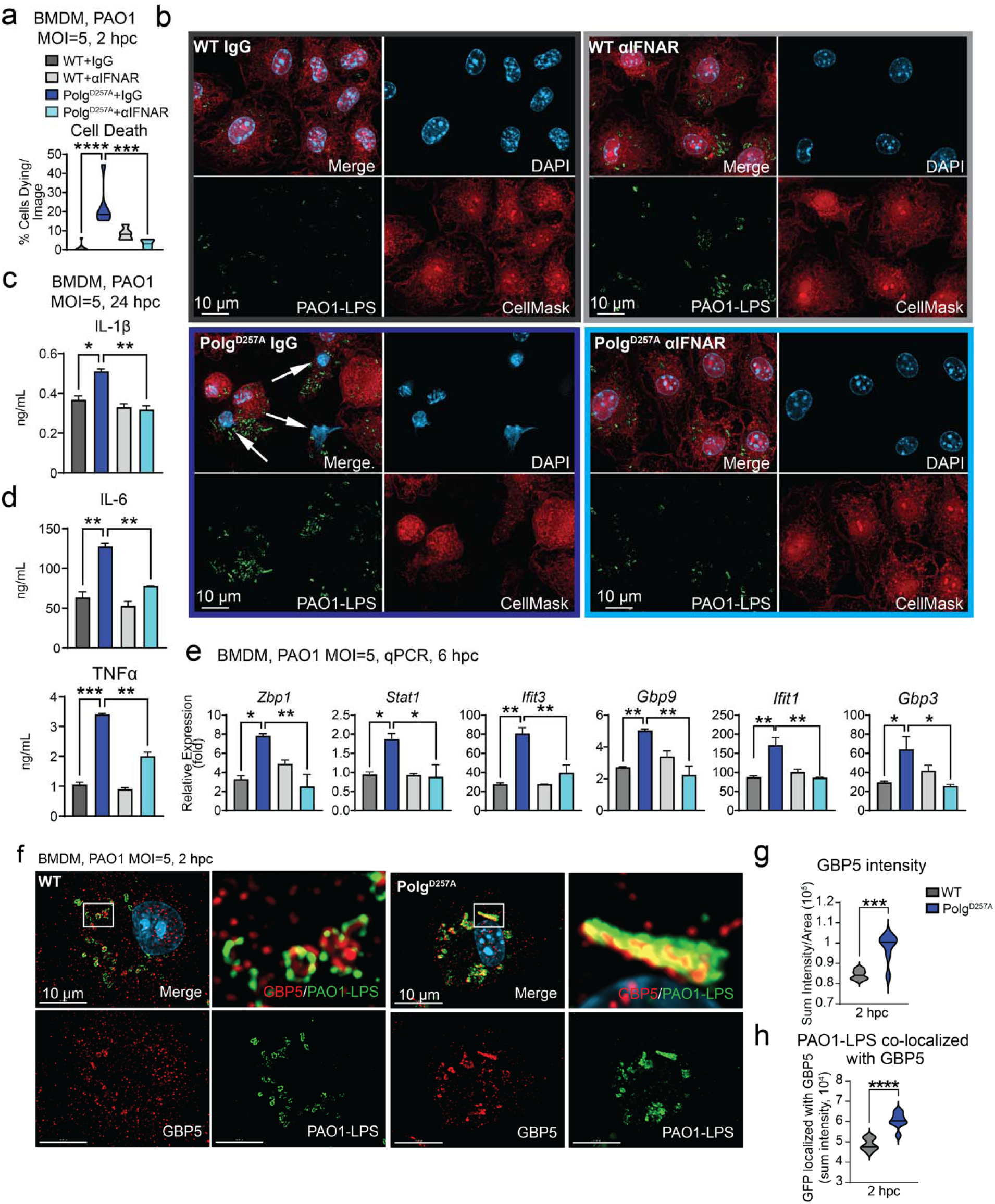
Elevated interferon signaling contributes to elevated GBP expression and hyperinflammation in *Polg^D257A^*macrophages. **(a)** Percent BMDMs dying after PAO1 and PAO1 plus anti-IFNAR blockade from **(b)** representative IF. N=5-6 IF images/genotype/condition. **(c)** ELISA analysis of pyroptotic cytokines in BMDM supernatant after PAO1 or PAO1 plus anti-IFNAR blockade. N= 2 biological replicates/genotype/condition. **(d)** ELISA analysis of inflammatory cytokines in BMDM supernatant after PAO1 or PAO1 plus anti-IFNAR blockade. N= 2 biological replicates/genotype/condition. **(e)** qRT-PCR analysis of ISGs in WT and *Polg^D257A^* BMDMs after PAO1 and PAO1 plus anti-IFNAR blockade. N=3 technical replicates/genotype/condition. **(f)** Representative IF of BMDMs 2 hpc with PAO1 at a multiplicity of infection (MOI) of 5. Cells were fixed, permeabilized, and stained with DAPI to mark the nucleus, anti-LPS antibody to stain PAO1 LPS, and anti-GBP5 antibody. **(g)** Sum intensity of GBP5 per cell area and **(h)** PAO1-LPS localized with GBP5 in BMDMs after PAO1 in **(f)** IF. N=7-8 images/genotype. Statistics: (g & h) Student’s t-test (a, c, d, & e) One-Way ANOVA. *P<0.05, **P<0.01, ***P<0.001, ****P<0.0001 Error bars represent SEM.

Prior work has shown that IFN-I signaling during bacterial infection drives the expression of guanylate binding proteins (GBPs)^42^, which facilitate innate immune responses and non-canonical inflammasome activation by disrupting membranes of pathogen-containing vacuoles, or pathogens themselves, to liberate LPS or other PAMPs^31,43–45^. Consistent with elevated IFN-I signaling, RNAseq analysis revealed more robust GBP expression in *Polg^D257A^* BMDMs after challenge with PAO1 (Supplementary Fig. 5f). Proteomics analysis also showed elevated GBPs, with marked increases in GBP2 and GBP5 in *Polg^D257A^* BMDMs six hours post challenge with PAO1 (Supplementary Fig. 5g). Immunofluorescence microscopy revealed increased GBP5 expression in resting *Polg^D257A^*BMDMs (Supplementary Fig. 5h-i), as well as during PAO1 challenge (Fig. 3f-g). Notably, we observed a more robust colocalization of GBP5 with PAO1 in *Polg^D257A^*macrophages after staining with an antibody specific for PA LPS. Moreover, GBP5 staining was more aggregated on PAO1-LPS positive bacteria than in WT BMDMs (Fig. 3h). These findings suggest that elevated IFN-I in *Polg^D257A^*macrophages augments GBP expression, likely enhancing sensing of LPS via caspase-11 that contributes to potentiated cell death and pro-inflammatory cytokine release.

### Caspase-11 activation, accentuated by GBP expression, controls pyroptosis and subsequent inflammation in *Polg^D257A^*macrophages

Our data suggest that potentiation of caspase-11-mediated pyroptosis drives excessive cell death and hyperinflammation in *Polg^D257A^* macrophages challenged with PAO1. Therefore, we hypothesized that elevated levels GBPs may contribute to the enhanced release of phagolysosomal PAO1 LPS for sensing by caspase-11 in the cytoplasm. We next used siRNA to silence caspase-11 or GBP2 and GBP5 in WT and *Polg^D257A^* BMDMs to repress this pathway. Knockdown efficiency of siRNAs was confirmed by qPCR of RNA at the time of infection (Supplementary Fig. 6a). Notably, caspase-11 silencing reduced elevated cell death, heightened cleavage of both GSDMD caspase-1, and increased secretion of IL-1 cytokines from *Polg^D257A^*BMDMs 2 hours post challenge with PAO1, while having little effect on WT BMDMs (Fig 4a-d). Similar results were obtained by silencing GBP2 and GBP5 (Fig. 4a-d). Furthermore, heightened caspase-11 association with LPS in *Polg^D257A^* BMDMs was reduced upon either caspase-11 or GBP silencing, although to a lesser extent after GBP knockdown (Fig. 4e). Caspase-11 or GBP knockdown lowered hypersecretion of proinflammatory cytokines in *Polg^D257A^*BMDMs at the twenty-four hour timepoint (Fig. 4f). Finally, we utilized an LPS-deficient PAO1 mutant to further investigate caspase-11 activation. The *algC* gene product is important for synthesis of the LPS core, and therefore this mutant displays altered lipid A levels and lower LPS overall^46^, which we confirmed by LPS staining intensity as measured with an anti-PAO1 LPS antibody (Supplementary Fig. 6b). Indeed, *Polg^D257A^* BMDMs challenged with PAO1-*algC* displayed reduced cell death, inflammasome activation, IL-1β release, IFN-I secretion, and pro-inflammatory cytokine production (Supplementary Fig. 6c-g), further supporting that increased detection of PAO1 LPS by caspase-11 is a predominant driver of hyperinflammatory phenotypes in *Polg^D257A^*macrophages.

**Fig. 4:**
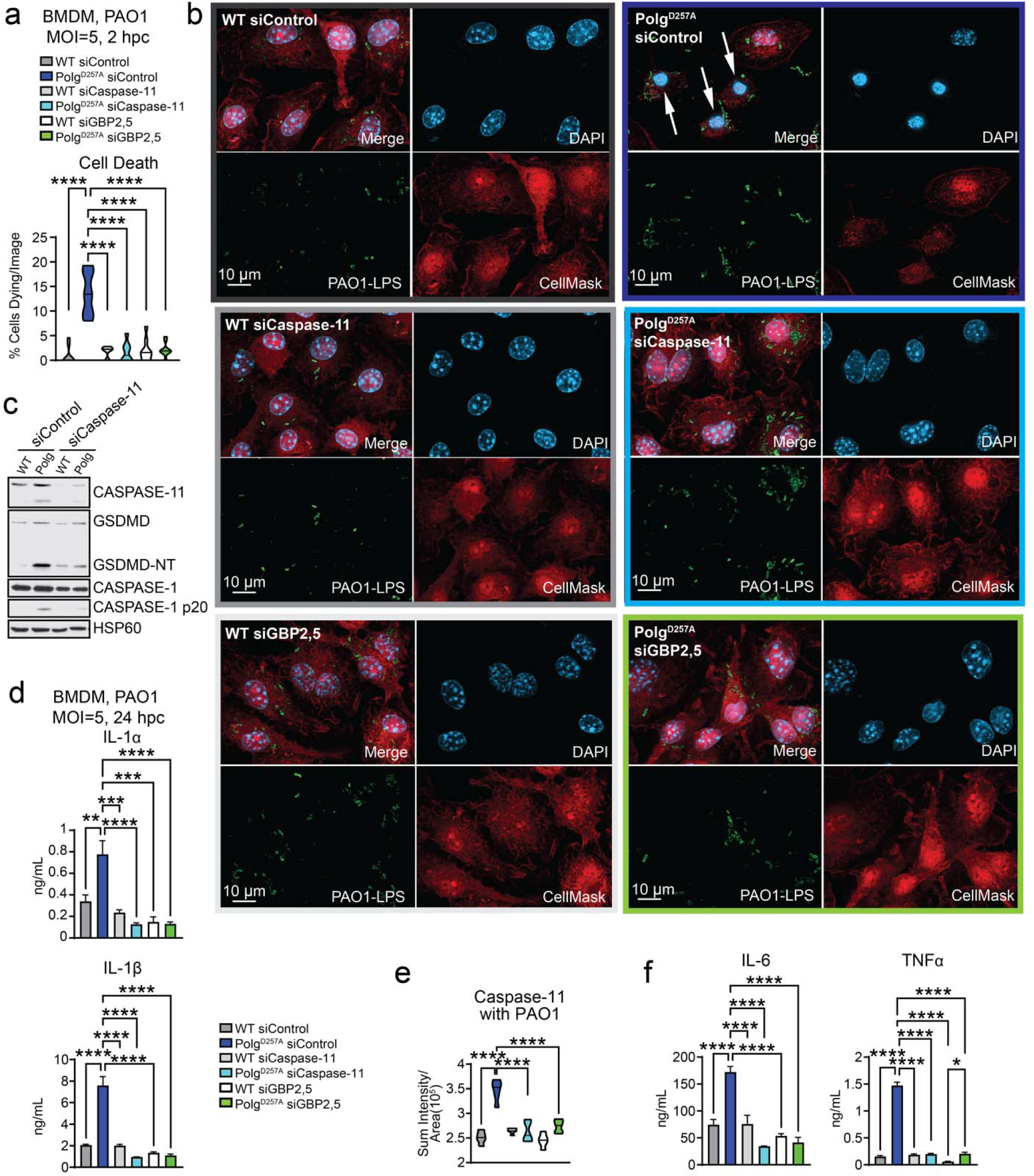
Caspase-11 activation, accentuated by GBPs, induces cell death and subsequent inflammation in *Polg^D257A^* macrophages. **(a)** Percent BMDMs dying after being given an siRNA control or siRNA to knock down caspase-11 or GBPs and then challenged with PAO1 from **(b)** representative IF. N=8-9 IF images/genotype/condition. **(c)** Western blot analysis of lysates from BMDMs challenged with PAO1 after siRNA knock down of caspase-11. **(d)** ELISA analysis of IL-1 cytokines in BMDM supernatant after being given an siRNA control or siRNA to knock down caspase-11 or GBPs and then challenged with PAO1. N= 3-4 biological replicates/genotype/condition. **(e)** Quantification of caspase-11 associated with LPS in BMDMs after knock down of caspase-11 or GBPs by siRNA and then challenge with PAO1. N=5 IF images/genotype/condition. **(f)** ELISA analysis of inflammatory cytokines in BMDM supernatant after being given an siRNA control or siRNA to knock down caspase-11or GBPs and then challenged with PAO1. N= 4 biological replicates/genotype/condition. Statistics: (a, d, e & f) One-Way ANOVA. *P<0.05, **P<0.01, ***P<0.001, ****P<0.0001 Error bars represent SEM. Comparisons between WT siCtrl and WT siCaspase-11 or WT siGBP2,5 that are not shown are n.s.

### Increased intracellular availability of LPS enhances caspase-11 activation and initiation of IL-1 signaling in *Polg^D257A^* macrophages

Transmission electron microscopy (TEM) revealed that while *Polg^D257A^*BMDMs have similar nuclear and cellular area at baseline (Supplementary Fig. 7a), *Polg^D257A^* BMDMs present condensed nuclei and enlarged phagolysosomes 2 hours post challenge with PAO1 (Fig. 5a-b and Supplementary Fig. 7c). Mitochondrial dysfunction can drive lysosomal defects and leakiness^47–49^, and we observed abnormal phagolysosomes in *Polg^D257A^* BMDMs by TEM two hours after PAO1 (Fig. 5a-b). We also observed increased presence of vesicles adjacent to PAO1 in *Polg^D257A^* phagolysosomes, which resembled bacterial outer membrane vesicles (OMVs)^50,51^. At eight hours post PAO1 challenge, *Polg^D257A^* BMDMs contained more intact bacteria with OMVs that were contained in phagolysosomes with poorly defined membranes (Supplementary Fig. 7-e). This was accompanied by increased PAO1 protein expression as assessed by proteomics analysis (Supplementary Fig. 7f). Oxidative stress is a critical driver of PA OMV generation, and we noted increased reactive oxygen species^20^ and nitrite production from *Polg^D257A^* BMDMs after PAO1 challenge (Supplementary Fig. 7g). Therefore, we hypothesized that caspase-11 hyperactivity in *Polg^D257A^* macrophages may be due to increased sensing of LPS on OMVs or lingering bacteria, leading to more robust recruitment and activation of caspase-11. Quantitation of IF imaging showed a greater intensity of caspase-11 and GBP5 colocalizing with PAO1 in *Polg^D257A^* BMDMs (Fig. 5c-d), which was not observed after challenge with LPS-deficient PAO1-*algC* (Supplementary Fig. 7h-i). To determine if these phenotypes were in part due to LPS uptake and/or degradation kinetics, we utilized a fluorescently labeled LPS. Notably, WT and *Polg^D257A^* BMDMs took up comparable amounts of LPS after LPS priming followed by LPS transfection at acute timepoints (Supplementary Fig. 8a-b). However, we observed more fluorescent LPS in the cytosol seven hours after LPS transfection of LPS primed BMDMs (Supplementary Fig. 8c-d), as well as significantly more caspase-11 associated with LPS by IF analysis (Supplementary Fig. 8e-f). WT and *Polg^D257A^* BMDMs contained similar levels of fluorescent LPS signal immediately after passive uptake from the culture media (Supplementary Fig. 8g-h), and *Polg* mutant BMDMs did not release IL-1β at early timepoints (Supplementary Fig. 8k). However, 48 hours after the addition of LPS to the cell culture media, *Polg^D257A^* BMDMs had a marked increase of cell-associated LPS (Supplementary Fig. 8i-j) and released small amounts of IL-1β (Supplementary Fig. 8k). Together, these data suggest that elevated levels of GBPs and caspase-11 in *Polg^D257A^* macrophages, coupled with increased bacterial OMVs and LPS in the cytosol, promote enhanced caspase-11 activation, leading pyroptosis and hyperinflammation.

**Fig. 5:**
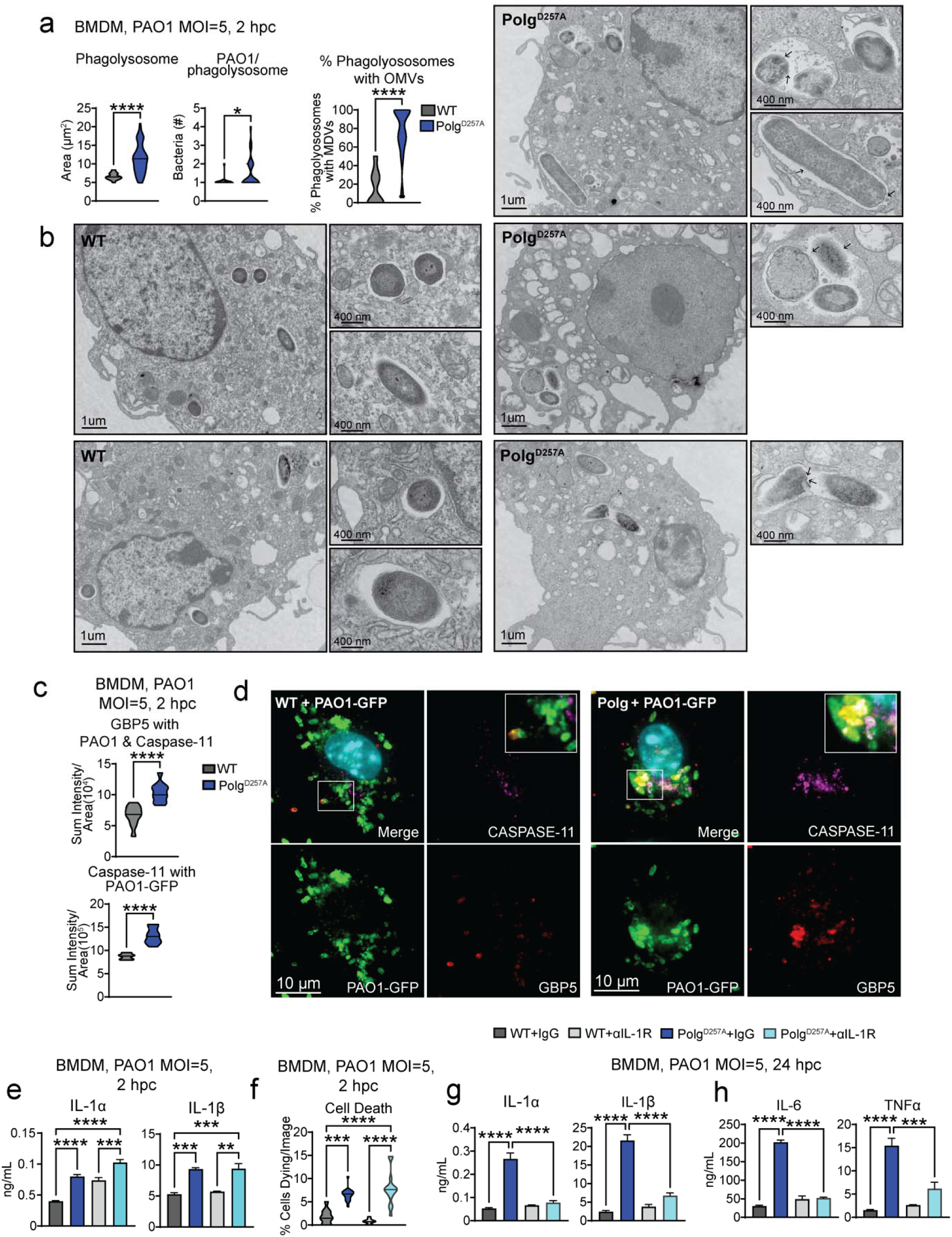
Elevated levels of caspase-11 binding to LPS contributes to potentiated pyroptosis in *Polg^D257A^* macrophages. **(a)** Quantitation of PAO1 and lysosomes in BMDMs from **(b)** TEM images. Black arrows highlight outer membrane derived vesicles. N= 9 images/condition. **(c)** Quantification of capase-11 and GBP5 intensity in the area containing caspase-11, GBP5, and PAO1 from representative IF images **(d)**. N=10-12 IF images/genotype/condition. **(i)** ELISA analysis of IL-1 cytokines in BMDM supernatant after PAO1 or PAO1 plus anti-IL-1R blockade 2 hpi. N= 4 biological replicates/genotype/condition. **(j)** Percent BMDMs dying after PAO1 and PAO1 plus anti-IL-1R blockade from **(Supplementary Fig. 9b)** representative IF. N=11-13 IF images/genotype/condition. **(k)** ELISA analysis of IL-1 cytokines in BMDM supernatant after PAO1 or PAO1 plus anti-IL-1R blockade 24 hpi. N= 4 biological replicates/genotype/condition. **(l)** ELISA analysis of inflammatory cytokines in BMDM supernatant after PAO1 or PAO1 plus anti-IL-1R blockade 24 hpi. N= 4 biological replicates/genotype/condition. Statistics: (a, c, e, & g) Student’s t-test (i, j, k, & l) Two-Way ANOVA. *P<0.05, **P<0.01, ***P<0.001,

In WT macrophages, Caspase-11 and caspase-1 are not directly involved in the generation of nuclear-factor kappa B (NF-κB)-dependent cytokines such as TNFα and IL-6. Therefore, we hypothesized that elevated caspase-11 activity may potentiate autocrine or paracrine signaling via the early release of alarmins that hyperactive PA-induced IFN-I or NF-κB. A kinetic stimulation study revealed increased phosphorylation of factors involved in IFN-I signaling (i.e. TBK1, STAT1) in *Polg^D257A^* BMDMs at early timepoints after PAO1 challenge, as well as elevated phospho-IκBα levels at early timepoints (Supplementary Fig. 9a). As mentioned above, IL-1α release is associated with caspase-11 activation and GSDMD cleavage^52^. IL-1α signaling through the IL-1 receptor (IL-1R) activates NF-κB and induces transcription of pro-IL-1β, inflammasome components including NLRP3, and other proinflammatory cytokines including TNFα and IL-6 ^53–55^. To determine if early caspase-11-dependent IL-1α signaling feeds forward to potentiate cytokine secretion in *Polg^D257A^* BMDMs at later timepoints, we used a neutralizing antibody to block IL-1R signaling. Notably, the IL-1R antibody did not reduce elevated IL-1 cytokine release, cell death, or GSDMD and caspase-11 cleavage in *Polg^D257A^* BMDMs two hours post challenge (Fig. 5e-f, Supplementary Fig. 9b-c). However, IL-1R blockade was sufficient to markedly decrease IL-1 cytokines, IL-6, and TNFα from *Polg^D257A^* BMDMs 24 hours post PAO1 challenge (Fig. 5g-h). Overall, these data indicate that the rapid, caspase-11-dependent release of IL-1α after PAO1 challenge drives feed forward NF-κB and IFN signaling to potentiate inflammatory cytokine production in *Polg^D257A^*macrophages.

### A novel *Polg^R292C^* mouse model incorporating a deleterious patient variant exhibit macrophage mtDNA depletion, elevated IFN-I responses, increased pyroptosis, and hyper-inflammation after challenge with PAO1

To determine if our findings in *Polg^D257A^* macrophages are more broadly applicable to other models of OXPHOS deficiency due to mtDNA perturbations, we leveraged a novel model of Polg-related mitochondrial disease harboring a single amino acid missense mutation (Arg292Cys) in exon 4 of the exonuclease domain of Polg (Supplementary Fig. 10a), which is homologous to the human Arg309Cys (c.925C>T) pathogenic variant^56,57^. We generated these mice using CRISPR/Cas9 editing of ES cells and verified germline transmission of the mutation by sequencing (Supplementary Fig. 10b). The Arg309Cys variant alone or in combination with other *Polg* variants causes reduced mtDNA copy number, mtDNA deletions, and severe phenotypes including mitochondrial myopathy, encephalopathy, lactic acidosis and stroke-like episodes (MELAS), peripheral neuropathy, ataxia, and epilepsy in patients. Although being born below expected mendelian ratios, homozygous *Polg^R292C^*mice are viable and display a reduced mitochondrial DNA copy number and elevated ISGs in multiple tissues (Supplementary Fig. 10c-e).

BMDMs derived from *Polg^R292C^* mice exhibited mtDNA depletion at rest (Supplementary Fig. 11a), as well as lower numbers of nucleoids and fragmented mitochondrial networks after stimulation with LPS (Supplementary Fig. 11b-c). In line with our data from the *Polg^D257A^* mutator mice, *Polg^R292C^*BMDMs displayed a constitutive elevation of ISGs at rest and potentiated ISG expression after LPS stimulation (Supplementary Fig. 10d-e). Using extracellular flux analysis, we also noted markedly reduced basal and maximal respiration after LPS stimulation (Supplementary Fig. 11f-h), indicating that *Polg^R292C^* macrophages exhibit OXPHOS deficiencies resembling those of *Polg^D257A^* BMDMs (Supplementary Fig. 4l-m).

After challenge with PAO1, *Polg^R292C^* BMDMs exhibited increased ISG and GBP expression (Fig. 6a), while also secreting more cytokines over a twenty-four-hour time course (Fig. 6b). Similar to *Polg^D257A^* BMDMs, *Polg^R292C^*exhibited elevated rates of PAO1-induced cell death, IL-1 cytokine secretion, and proinflammatory cytokine production, which were blunted by treatment with caspase-1 inhibitor AcYVAD (Fig. 6c-e, Supplementary Fig. 12) or IFNAR blockade (Fig. 6f-h). Finally, silencing of caspase-11 in *Polg^R292C^* BMDMs (Fig. 6i) alleviated acute cell death (Fig. 6j-k), IL-1 release (Fig. 6l), and secretion of IL-6 and TNFα at late timepoints (Fig. 6m). These data suggest that elevated IFN-I, hyperinflammation, and pyroptosis in response to PAO1 are common features of macrophages with deficient Polg exonuclease activity, including the MtD-relevant Arg292Cys variant.

**Fig. 6:**
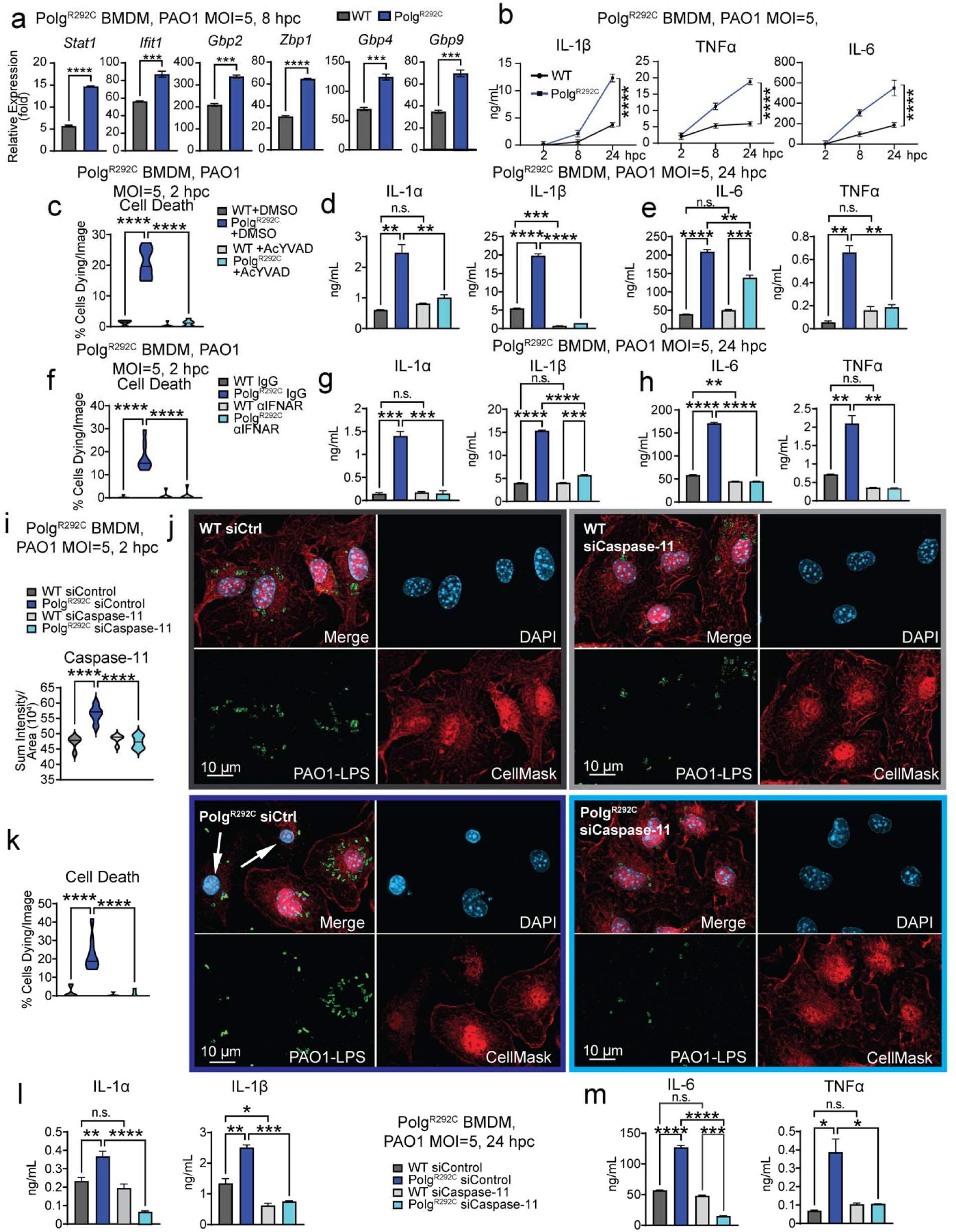
Macrophages incorporating a deleterious patient mutation (*Polg^R292C^*) exhibit mtDNA depletion, elevated IFN-I responses, increased pyroptosis, and hyper-inflammation after challenge with PAO1. **(a)** qRT-PCR analysis of ISGs in BMDMs after PAO1, N=3 technical replicates/genotype. **(b)** ELISA analysis of inflammatory cytokines in BMDM supernatant after PAO1. N= 4 biological replicates/genotype/timepoint. **(c)** Percent BMDMs dying after PAO1 and PAO1 plus caspase-1/11 inhibitor. N=7 IF images/genotype/condition. **(d)** ELISA analysis of IL-1 cytokines in BMDM supernatant after PAO1 or PAO1 plus caspase-1/11 inhibitor. N= 2 biological replicates/genotype/condition. **(e)** ELISA analysis of proinflammatory cytokines in BMDM supernatant after PAO1 or PAO1 plus caspase-1/11 inhibitor. N= 2 biological replicates/genotype/condition. **(f)** Percent BMDMs dying after PAO1 and PAO1 plus anti-IFNAR blockade from. N=7 IF images/genotype/condition. **(g)** ELISA analysis of IL-1 cytokines in BMDM supernatant after PAO1 or PAO1 plus anti-IFNAR blockade. N= 2 biological replicates/genotype/condition. **(h)** ELISA analysis of pro-inflammatory cytokines in BMDM supernatant after PAO1 or PAO1 plus anti-IFNAR blockade. N= 2 biological replicates/genotype/condition. **(i)** Quantification of caspase-11 in BMDMs after knock down by siRNA and PAO1 challenge. N=5 IF images/genotype/condition. **(k)** Percent BMDMs dying after PAO1 and PAO1 plus caspase-11 knock down by siRNA from **(j)** representative IF. N=8 IF images/genotype/condition. **(l)** ELISA analysis of IL-1 cytokines in BMDM supernatant after PAO1 or PAO1 caspase-11 knock down by siRNA. N= 4 biological replicates/genotype/condition. **(m)** ELISA analysis of pro-inflammatory cytokines in BMDM supernatant after PAO1 or PAO1 caspase-11 knock down by siRNA. N= 2 biological replicates/genotype/condition. Statistics: (a) Students t-test. (b) Two-Way ANOVA. (c, d, e, f, g, h, i, k, l, & m) One-Way ANOVA. *P<0.05, **P<0.01, ***P<0.001, ****P<0.0001 Error bars represent SEM.

### Intratracheal instillation of a non-lethal dose of PAO1 elicits greater lung cytokine production and alterations in lung immune cell populations in *Polg^D257A^* mice

Patients with primary MtD often experience recurrent viral and bacterial infections of the respiratory tract, including infections by PA^5^. These events can trigger metabolic decompensation and can progress to systemic inflammatory endpoints and/or sepsis. However, few studies have examined infection kinetics and lung immune responses in Polg-related mitochondrial disorders. We therefore employed intratracheal instillation protocols to administer LPS or PAO1 into the lungs of *Polg^D257A^* mutator mice. Histological analysis of resting *Polg^D257A^* lungs did not reveal obvious differences in morphology or cellular composition (Supplementary Fig. 13a-c), although consistent with our earlier findings^20^, basal expression of ISGs were elevated compared to WT littermates (Supplementary Fig. 13d). After LPS was instilled into lungs through cannulation of the trachea, *Polg^D257A^* mice secreted more TNFα and IL-6 at all timepoints and increased IL-1β at the 12-hour timepoint in the bronchial alveolar lavage fluid (BAL) compared to WT mice (Supplementary Fig. 13e).

To next characterize lung innate immune responses to PAO1 infection in the *Polg^D257A^* mice, we challenged mice with a non-lethal dose of PAO1^58^. Both WT and *Polg^D257A^*mice infected with 1×10^4^ colony forming units (CFU) of PAO1 via intratracheal instillation lost weight initially but returned to normal weight (Supplementary Fig. 14a) and exhibited no culturable bacteria (data not shown) 3 days post infection. Flow cytometric analysis of lung cells in BAL fluid revealed an overall decrease in the percentage of CD45^+^ immune cells at rest and after infection, consistent with general leukopenia noted in *Polg^D257A^* mice^20,25^ (Supplementary Fig. 14b-c). Analysis of specific immune cell markers on lung CD45^+^ cells revealed few changes in BAL populations (Supplementary Fig. 14d-l) between uninfected WT and *Polg^D257A^*mice, although there were more SiglecF^+^ and F4/80^+^ cells at rest perhaps indicating elevated numbers of lung macrophages (Supplementary Fig. 14e,g). Six hours post infection, there was a marked reduction in SiglecF^+^ cells in *Polg^D257A^* mice relative to WTs and an increase in CD3^+^ T cells (Supplementary Fig. 14e,i). Cells labeling with the neutrophil marker, Ly6G, were dramatically increased at 6 hours, although there were no differences between WT and *Polg^D257A^*mice (Supplementary Fig. 14k). At 48 hours post infection, neutrophils remained the most predominant CD45^+^ population in both genotypes, but we observed a reversion toward increased numbers of cells staining positive for macrophage markers SiglecF^+^ and F4/80^+^ in *Polg^D257A^* mice (Supplementary Fig. 14e,g,k). Despite modest changes in cellular composition of the BAL fluid, *Polg^D257A^*mice had elevated numbers of CD45^+^ cells expressing TNFα both at rest and six hours post infection (Supplementary Fig. 14m), consistent with increased TNFα production after LPS challenge.

Multianalyte analysis of the BAL fluid using Legendplex assays revealed 2-4 fold increases in lung pro-inflammatory cytokines in *Polg^D257A^*mice 6 hours post PAO1 infection (Fig. 7a, Supplementary Fig. 15a). Consistent with resolution of infection in both genotypes, cytokine differences in the BAL of WT and *Polg^D257A^* mice were less pronounced at 48 hours post infection (Supplementary Fig. 15b). However, IFNβ and inflammatory cytokines associated with the skewing and differentiation of T cell Th1 (IL-27) and Th17 subsets (IL-23) remained elevated in *Polg^D257A^*mice in an IFNAR-dependent fashion (Supplementary Fig. 15b). Analysis of lung protein extracts revealed elevated caspase-1 cleavage, as well as increased caspase-11, NLRC4, and NLRP3 expression in *Polg^D257A^* mice 48 hours post intratracheal instillation of 1×10^4^ CFU of PAO1. GSDMD, GBP5, and other ISGs were also elevated in *Polg^D257A^* lungs, consistent with sustained IFNβ expression observed by Legendplex (Supplementary Fig. 15c).

**Fig. 7:**
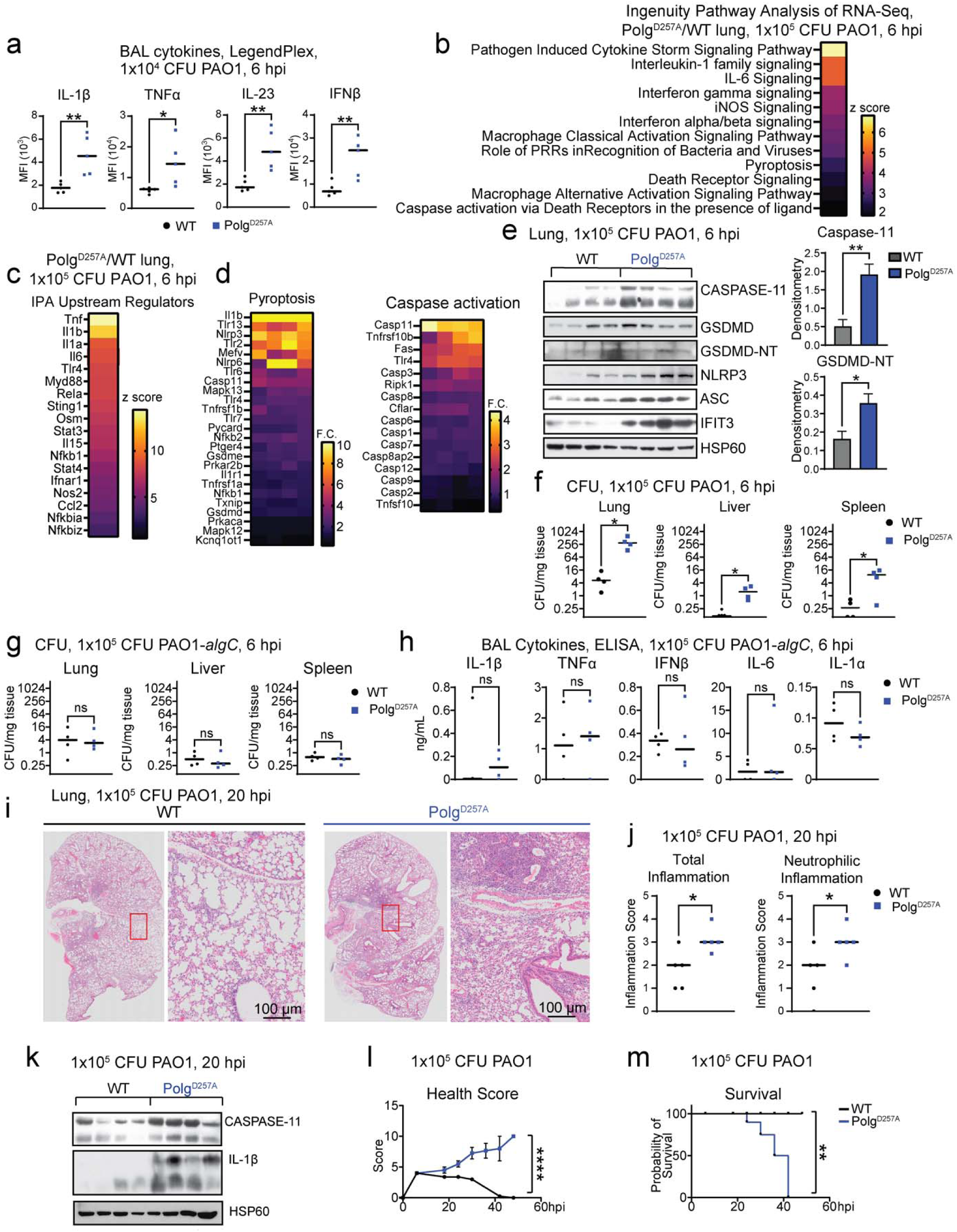
Intratracheal instillation of PAO1 into lungs elicits greater inflammation, caspase-11 expression, and death of *Polg^D257A^* mutant mice. **(a)** LegendPlex pro-inflammatory cytokine analysis of bronchial alveolar lavage (BAL) 6 hours post infection (hpi) with 1×10^4^ colony forming units (CFU) of PAO1. N=5 mice/genotype **(b)** Ingenuity Pathway Analysis (IPA) of RNA-seq from lung showing Z-scores of pathways in *Polg^D257A^*homozygous mutant BMDMs compared to wild-type (WT) after PAO challenge. N=3 biological replicates/genotype/condition. **(c)** IPA Upstream Regulator analysis showing Z-scores in *Polg^D257A^* homozygous mutant lungs compared to WT after 6 hour PAO1 challenge. **(d)** Log^2^ fold changes of representative genes in IPA pathways from (**a**) in *Polg^D257A^*lungs compared to WT after PAO1. **(e)** Protein expression or WT and *Polg^D257A^* lung homogenates 6 hpi with 1×10^5^ CFU of PAO1 and densitometry. **(f)** Colony counts from homogenized tissue infected with 1×10^5^ PAO1 for 6 hours. N=4 mice/genotype. **(g)** Colony counts from homogenized tissue infected with 1×10^5^ PAO1-*algC* for 6 hours. N=4 mice/genotype. **(h)** ELISA analysis of pro-inflammatory cytokines in BAL of mice infected with 1×10^5^ PAO1-*algC* for 6 hours. N=4 mice/genotype. **(i)** Representative hematoxylin and eosin (H&E) staining of lung sections 20 hpi with 1×10^5^ CFU of PAO1. **(j)** Graphs of pathology scoring from H&E stained lungs in **(i)** histology. N=5 mice/genotype **(k)** Protein expression or WT and *Polg^D257A^* lung homogenates 20 hpi with 1×10^5^ CFU of PAO1. **(l)** Illness scoring of WT and *Polg^D257A^* mice over PAO1 infection time course. N=4-5 mice/genotype **(m)** Kaplan-Meier survival curve after infection with 1×10^5^ CFU of PAO1. N= 4-5 mice/genotype Statistics: (a, e, f, g, h, & j) Students t-test (l) Two-Way ANOVA (m) Kaplan-Meijer Survival Curve. *P<0.05, **P<0.01, ***P<0.001, ****P<0.0001 Error bars represent SEM.

Given that recurrent respiratory tract infections are often observed in patients with MtD, we next infected mice with 1×10^4^ CFU PAO1, followed by a 7-day rest period, then re-infected with 1×10^4^ CFU PAO1. Histological analysis of lungs 24 hours following re-infection revealed monocyte and granulocyte infiltration into lungs, areas of vascular hemorrhage, and more eosin positive cells in *Polg^D257A^* lungs relative to WT (Supplementary Fig. 16a-b). Lung IL-12 family cytokines and IFNβ were significantly higher in *Polg^D257A^* mice at the 24-hour harvest point following re-infection, while other cytokines were trending higher in *Polg^D257A^*mice, although they did not reach statistical significance (Supplementary Fig. 16c). We also observed higher bacterial burdens in the lungs and spleens of *Polg^D257A^* mice compared to WT controls (Supplementary Fig. 16d). Together, these data indicate that *Polg^D257A^* mice exhibit hyperinflammation and impaired antibacterial immunity upon re-infection with PAO1.

### PAO1 elicits greater caspase-11 expression, increases inflammation, and reduces survival of *Polg* mutant mice

We next utilized a higher dose of PAO1 to further assess innate immune responses *Polg* mutant mice. RNAseq of lung tissue collected six hours after intratracheal instillation of 1×10^5^ CFU PAO1 revealed a striking upregulation of many inflammatory-, IFN-, and programmed cell death-related pathways in *Polg^D257A^* mice relative to WT littermates (Fig. 7b-c, Supplementary Fig. 17a). In agreement with macrophage data, transcriptional signatures of pyroptosis were apparent, with *Casp11*, *Nlrp3*, *Il1a*, and *Il1b* some of the most abundantly enriched transcripts in *Polg^D257A^* versus WT lung lysates (Fig. 7d). *Polg^R292C^*lung homogenates also displayed upregulation in transcriptional pathways indicative of hyperinflammation, elevated IFN signaling, and potentiated pyroptotic cell death 6 hours post PAO1 infection (Supplementary Fig. 17b-d). Western blotting of *Polg^D257A^*lung homogenates reinforced RNAseq data, showing increased protein expression of pyroptosis factors including caspase-11 and cleaved GSDMD (Fig. 7e), as well as a depletion in OXPHOS Complex I and IV subunits 6 hours post infection (Supplementary Fig. 17e). Moreover, *Polg^D257A^* mice exhibited elevated bacterial burdens in the lung, liver, and spleen (Fig. 7f). Increased bacterial dissemination was accompanied by elevated plasma proinflammatory cytokines, as well as elevated ISG expression with reduced OXPHOS in *Polg^D257A^* liver homogenates (Supplementary Fig. 17f-g). We next utilized the PAO1-*algC* mutant to further define whether reduced LPS sensing via caspase-11 would alter response to infection in *Polg^D257A^* mice. Compared to WT PAO1, PAO1-*algC* did not disseminate (Fig. 7g), or trigger increased expression of pyroptotic proteins and cytokines in *Polg^D257A^*lungs relative to WTs (Fig. 7h, Supplementary Fig. 17h). Overall, these data suggest *Polg* mutant mice display increased caspase-11-medated sensing of PAO1 LPS, elevated bacterial burdens, and hyperinflammatory responses during acute PAO1 infection

At later timepoints, *Polg^D257A^* lungs displayed elevated neutrophilic inflammation and overall inflammatory scores via histological analysis (Fig. 7i-j, Supplementary Fig. 17i), despite harboring similar bacterial burdens to WT mice (Supplementary Fig. 17j). However, caspase-11 and IL-1β remained elevated in lung lysates of *Polg^D257A^*mice compared to WT littermates twenty hours post infection (Fig. 7k). Using a comprehensive health scoring system to track breathing, hydration, body appearance, and activity, we noted that *Polg^D257A^*mice appeared sicker and continued to decline in health, whereas WT animals recovered from infection (Fig. 8l). Even though we observed a steady decline in the health of *Polg^D257A^* mice at 48 hours after infection, PAO1 were largely cleared in both *Polg^D257A^*and WT animals (Supplementary Fig. 17k). However, *Polg^D257A^* mice sustained elevated ISG expression in peripheral tissues, suggesting sustained IFN-I signaling (Supplementary Fig. 17l-m). Finally, we observed that a greater number of *Polg^D257A^* mice reached a moribund state after infection with 1×10^5^ CFU of PAO1 relative to WT littermates (Fig. 8m). Overall, these in vivo data suggest that *Polg^D257A^* mice are more prone to excessive inflammation and acute lung injury following infection with PAO1.

## Discussion

Patients with MtD experience a greater frequency of bacterial and viral infections that can rapidly progress to sepsis or systemic inflammatory response syndrome (SIRS), leading to premature death^5^. Recurrent infections can also accelerate MtD progression by promoting metabolic decompensation, a life-threatening condition that hastens deterioration of mitochondrial function^59–61^. Polg is the sole mitochondrial DNA polymerase, and variants in the *POLG* gene are a prevalent cause of MtD^62^. There are several clinical reports of bacterial or viral infections unmasking Polg-related MtD symptoms and hastening disease progression^63,64^. Moreover, one study revealed the presence of inflammatory cytokines in the cerebrospinal fluid of a patient with Polg-related MtD^12,19^. As cytokine-mediated toxicity is an important driver of metabolic decompensation^7^, there is a critical need to understand immune responses in patients with Polg-related disorders and other MtD to manage infections and inflammation-related complications. This study builds upon our prior work to characterize innate immune responses in mouse models of MtD^20,65^ and leverages bacterial infection, cytokine profiling, and in vitro and ex vivo cellular assays to define molecular and cellular nodes governing hyperinflammatory immune phenotypes in two models of Polg-related MtD (Supplementary Fig. 18).

Our prior work revealed that elevated IFN-I responses in *Polg^D257A^*mutator mice are mediated by chronic mtDNA release and cGAS-STING signaling^20^. Using our new, disease relevant *Polg^R292C^*mouse model, we observed increased IFN-I signatures basally and after challenge with PAO1, similar to *Polg^D257A^*. Analogous variants in human POLG at R309 affect proofreading activity^66^, and therefore we hypothesize that the *Polg^R292C^*exonuclease mutation also impairs replication fidelity, leading to mtDNA release and chronic IFN-I signaling. A recent study reported that fibroblasts from patients carrying the *POLG1* c.2243G>C, p.W748S mitochondrial recessive ataxia syndrome (MIRAS) allele display reduced IFN-I signaling in response to viral infection and nucleic acid stimuli^67^. The MIRAS variant affects POLG holoenzyme processivity but not the exonuclease domain, and thus impaired exonuclease activity leading to mtDNA mutations and deletions may selectively trigger IFN-I in the *Polg^D257A^*and *Polg^R292C^* backgrounds. Despite reduced IFN-I, mice homozygous for the MIRAS variant do exhibit increased necroptosis, elevated liver immune cell infiltration, and potentiated inflammatory cytokine secretion during viral infection^67^. Thus, inflammatory cell death and hyperinflammation are common features in both patient cells and mouse models of Polg-related MtD.

Our data reveal that IFN-I signaling is a key driver of caspase-11-dependent pyroptosis and hyperinflammation in *Polg* mutant macrophages. Pre-treatment with an anti-IFNAR neutralizing antibody was sufficient to reduce both augmented PI uptake after LPS transfection and cytokine hypersecretion after challenge with PAO1. IFN-I is critical to license caspase-11 expression^41^, but also to enhance expression of interferon-inducible GTPases like GBPs, which help to expose cytosolic LPS for detection^31,68–71^. We have found that macrophages from both the *Polg^D257A^* and our new MtD-relevant *Polg^R292C^* model exhibit enhanced expression of multiple GBPs at rest and after challenge with PAO1. Moreover, we observed that GBP5 strongly co-localizes with intracellular PAO1 and caspase-11 in *Polg^D257A^* BMDMs. GBPs are important in orchestrating caspase-11 activation to bacterial outer membrane vesicles (OMVs)^50,51^ and can localize to damaged cell membranes^45^. Although the mechanisms by which GBPs sense intracellular pathogens and facilitate caspase-11 activation are not fully understood, silencing of caspase-11 or GBP2 and GBP5 markedly reduced pyroptosis and the secretion of IL-1 and other cytokines from both *Polg* mutant BMDMs. TEM imaging also revealed altered PAO1-containing phagolysosomal membranes and increased bacteria-associated vesicles in *Polg^D257A^* BMDMs. Thus, it is likely that GBPs and caspase-11 more readily sense PAO1 LPS due to phagolysosmal leakiness, or elevated OMV generation, yet more work is required to fully define the underlying mechanisms.

Our data reveal that *Polg* mutant macrophages undergo rapid pyroptosis upon PAO1 challenge in a NLRP3- and caspase-1-dependent fashion. Although caspase-1 cleavage is elevated in *Polg^D257A^* compared to WT macrophages during PAO1 challenge and its inhibition is sufficient to block augmented PA-induced cell death and IL-1β secretion, we found that *Polg^D257A^* BMDMs are less responsive to direct activation of the NLRP3 and NLRC4 inflammasomes by their respective ligands. In contrast, activation of the caspase-11 inflammasome via LPS priming followed by transfection of LPS into the cytosol drove more PI uptake and IL-1β expression in *Polg^D257A^* BMDMs. Therefore, sensing of cytosolic PA LPS by the non-canonical inflammasome leads to more robust activation of the NLRP3 for caspase-1 dependent cell death and cytokine secretion. Notably, a PAO1-*algc* mutant lacking robust LPS synthesis did not hyper-stimulate pyroptosis and cytokine secretion, providing further support for caspase-11 as the predominant driver of inflammatory cell death in *Polg* mutant mice.

It is interesting to note that treatment of *Polg* mutant macrophages with caspase-1 inhibitor dramatically reduced nearly all cytokine levels in culture supernatants after challenge with PAO1. These include inflammasome-independent cytokines such as TNFα, IL-6, IFNγ, and IL-12 family members as well as the DAMP IL-1α, which does not require caspase-1 cleavage for secretion but is downstream of caspase-11 and GSDMD^37,72^. This is in contrast with cytokine secretion in WT BMDMs, where AcYVAD treatment only reduced IL-1β levels during challenge with PAO1. Similarly, anti-IFNAR blockade or caspase-11 and GBP knockdown reduced hyper-secretion of IL-1β levels and reduced IL-6 and TNFα in *Polg* macrophages, while having marginal effects on WT BMDM cytokine profiles. While blocking the IL-1R did not reduce early cell death in *Polg* mutant macrophages, it was sufficient to blunt elevated cytokines a day after PAO1 challenge. This suggests that early IL-1α release downstream of capsase-11 and GSDMD synergizes with IFN-I signaling to sustain hyperinflammation in PAO1-challenged *Polg* mutant BMDMs. Our data do not definitively rule out contributions for canonical inflammasome-mediated IL-1β in potentiating NF-κB-mediated cytokine release. However, the link between caspase-11 activation and IL-1α suggest that the early release of this alarmin may prime neighboring cells for activation of canonical inflammasomes, further promoting increased cell death and inflammation in *Polg* mutant macrophages.

Although it is appreciated that bacterial and viral infections are problematic in MtD, few studies have examined innate immune responses in relevant mouse models of disease. Using an intratracheal instillation model, we found that *Polg^D257A^* mice exhibit hyperactive innate immune responses resulting in elevated cytokine production, changes in lung immune cell dynamics recruitment, and increased morbidity following PAO1 infection. Notably, we observed elevated expression of inflammasome proteins including caspase-11, ISGs including GBPs, and increased caspase-1 cleavage in *Polg^D257A^* lung homogenates after infection. Remarkably, RNAseq analysis of PA infected lungs from 3-month-old *Polg^R292C^* mice also revealed increased pyroptosis, interferon, and hyperinflammatory signaling. These data agree with our in vitro findings implicating IFN-I as an overarching regulator of caspase-1/11 activation and hyperinflammation in our models. Additional studies are warranted to define the cellular populations that govern lung hyperinflammation and to determine the degree to which caspase-11 shapes inflammatory signaling and morbidity in *Polg* mutant mice in vivo.

While *Polg^D257A^* mice had higher bacterial burdens in the lung and peripheral tissues at acute timepoints, they ultimately managed to control the infection by 20 hours post infection and cleared bacteria by 48 hours. Despite this, lungs from *Polg^D257A^* mice were more inflamed at 20 hours post infection and the mice had more severe sickness scores, resulting in higher rates of morbidity. This suggests that the adverse outcomes observed in *Polg* mutant animals may be due to chronic and damaging hyperinflammation as opposed to impaired bacterial clearance. Therefore, therapeutic targeting of IFN-I and caspase-11 may represent promising avenues to restore proper immune balance patients with Polg-related MtD. Although our work does not include clinical data, two recent publications revealed elevated IFN-I signatures in the whole blood and peripheral blood mononuclear cells (PBMCs) from patients with diverse MtD disease variants^11,73^. Moreover, gene ontology analysis revealed heightened IL-1β signatures in MtD PBMCs^11^. Our study provides important new insight into the innate immune mechanisms governing infection-related hyperinflammation in mouse models of MtD. The consistency of our discoveries across two different *Polg* mutant models strengthens the relevance of our findings to human disease and suggest that IFN-I and inflammasome hyperactivity may be critical determinants of increased rates of sepsis and SIRS in patients with MtD.

---

**Supplementary Fig. 1:**
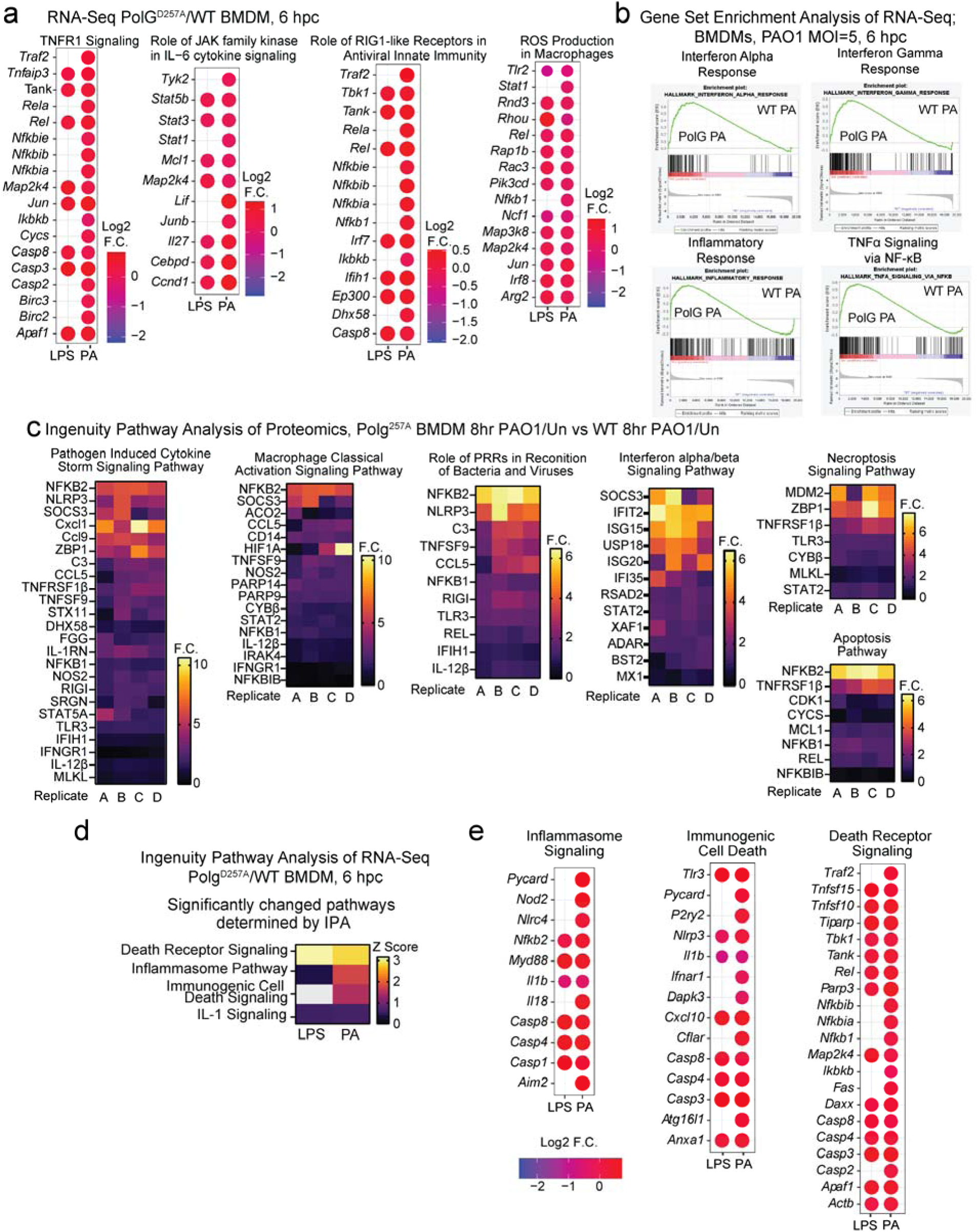
*Polg^D257A^* macrophages exhibit elevated IFN-I and pro-inflammatory gene expression when challenged with LPS or PAO1. **(a)** Log^2^ fold changes of representative genes in IPA pathways from (Fig. 1b) in *Polg^D257A^* BMDMs compared to WT after challenge with LPS or PAO1. N=3 biological replicates/genotype/condition. **(b)** Gene Set Enrichment (GSEA) pathways of RNA-seq data in WT ang *Polg ^D257A^* BMDMs after challenge with PAO1. Heatmaps represent fold changes of each group relative to WT PAO1 challenged BMDMs. N=3 biological replicates/genotype/condition. **(c)** Ingenuity pathway analysis of proteomics from *Polg^D257A^* (PAO1, 8hpc/Un)/WT (PAO1, 8 hpc/Un) BMDM proteins in upregulated pathways from Fig 1g. N=4 biological replicates **(d)** Ingenuity Pathway Analysis (IPA) of RNA-seq from bone marrow-derived macrophages (BMDMs) showing Z-scores of pathways related to cell death signaling in *Polg^D257A^*mutant BMDMs compared to wild-type (WT) after LPS or PAO1 challenge. N=3 biological replicates/genotype/condition. **(e)** Log^2^ fold changes of representative genes in IPA pathways from **(d)** in *Polg^D257A^*BMDMs compared to WT after LPS or PAO1.

**Supplementary Fig. 2:**
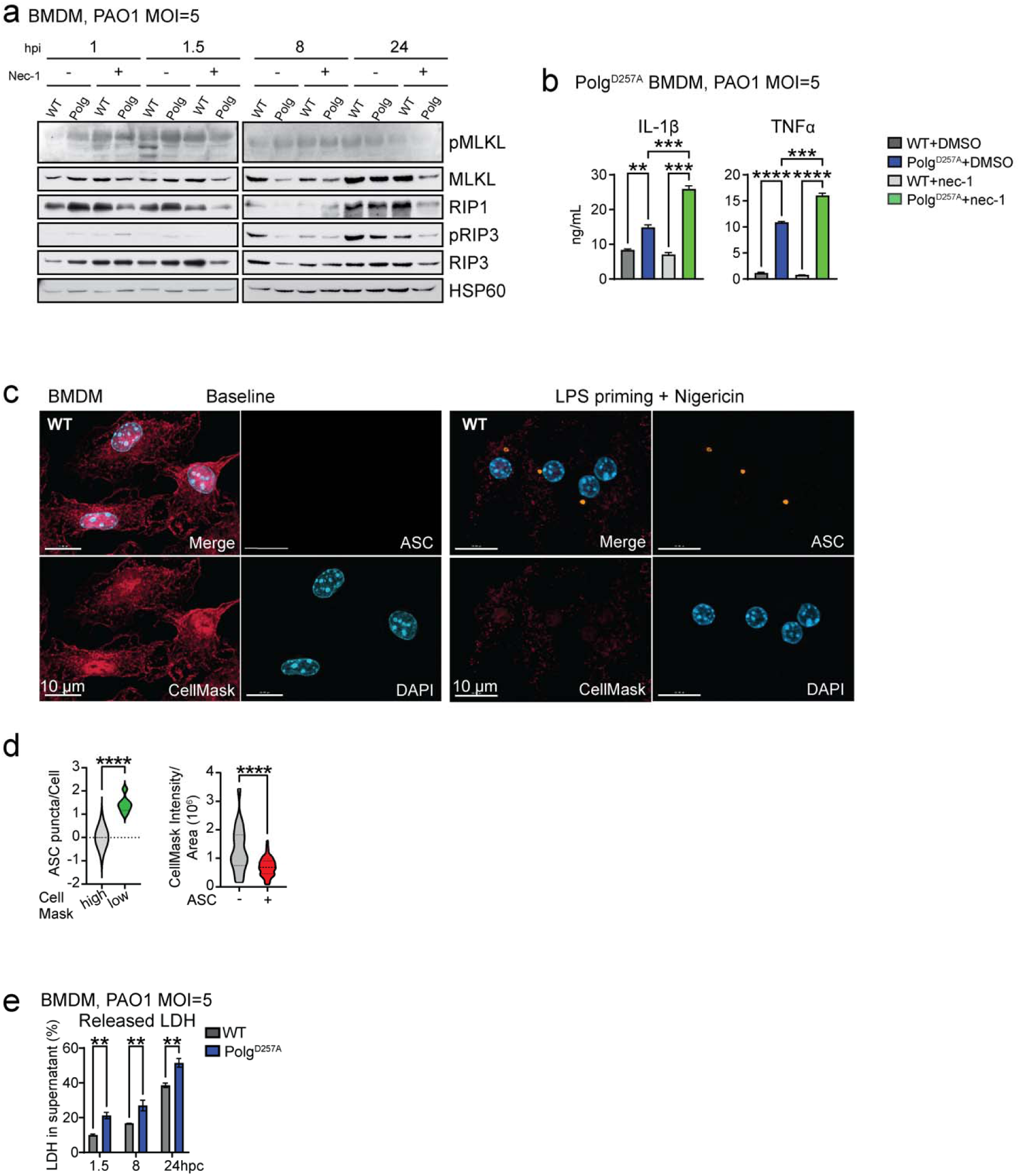
*Polg^D257A^*macrophages exhibit increased inflammatory cell death when exposed to bacteria. **(a)** Western blot of BMDM protein lysates after nec-1 priming plus PAO1 for protein involved in necroptosis. **(b)** ELISA analysis of inflammatory cytokines in BMDM supernatant after PAO1 or PAO1 plus necroptosis inhibitors. N= 2 biological replicates/genotype/condition/timepoint. **(c)** Representative IF of BMDMs unchallenged or primed with LPS for 4 hours and stimulated with nigericin for 1 hour to activate the NLRP3 inflammasome. Cells are stained with DAPI to mark the nucleus, CellMask to mark total cell volume, and anti-ASC antibody to mark puncta. **(d)** Number of ASC puncta per cell in cells with high and low CellMask staining in **(a)** IF. N= 7 images/condition. **(e)** Percent LDH released from BMDMs during PAO1 challenge. N=3 biological replicates/genotype/timepoint. Statistics: (d) Students t-test (b & e) Two-way ANOVA. *P<0.05, **P<0.01, ***P<0.001, ****P<0.0001 Error bars represent SEM.

**Supplementary Fig. 3:**
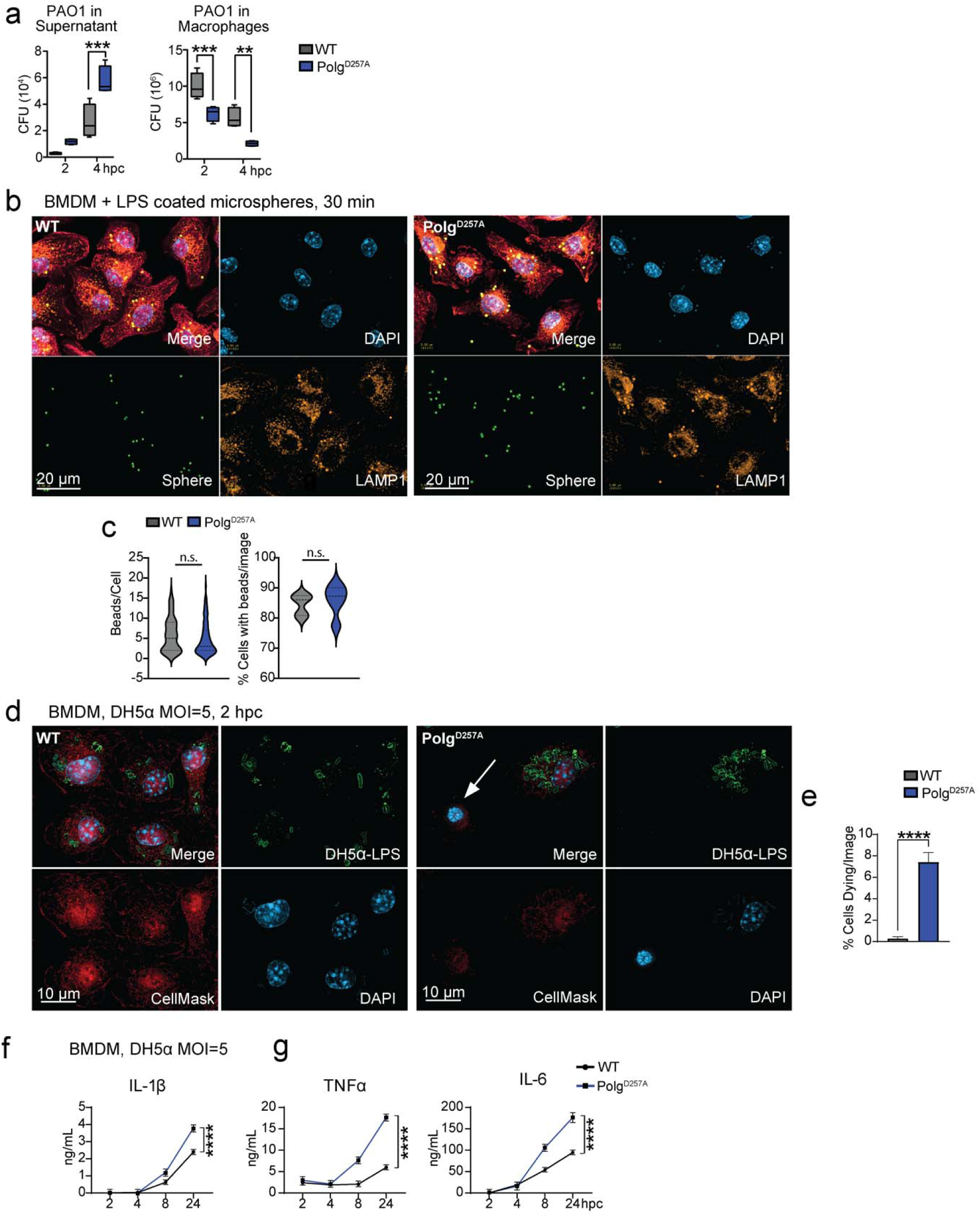
*Polg^D257A^*macrophages have a greater release of PAO1 that is not due to a deficiency in phagocytosis or PAO1 virulence factors. **(a)** PAO1 colony forming units obtained from BMDM supernatant or detergent lysates 2 and 4 hours post challenge (hpc). N=4 biological replicates/genotype/timepoint. **(b)** Representative IF of BMDMs phagocytosing LPS coated, fluorescent microspheres. Cells were stained with DAPI to mark the nucleus, CellMask to label the cell volume, and anti-LAMP1 to mark phagolysosomes. **(c)** Beads per cell quantified from **(e)** IF. N= 100 cells/genotype. **(d)** Percent BMDMs dying after challenge with DH5α **(e)** IF. N=7 IF images/genotype. **(e)** Representative IF of BMDMs stained with DAPI to mark the nucleus, CellMask to mark the total cell volume, and anti-LPS to mark DH5α bacteria. White arrows highlight dying cells characterized by condensed nuclei and loss of CellMask staining. **(f)** ELISA analysis of IL-1β in BMDM supernatants after challenge with *Escherichia coli* strain DH5α. N= 4 biological replicates/genotype/timepoint. **(g)** ELISA analysis of inflammatory cytokines in BMDM supernatants after challenge with *Escherichia coli* strain DH5α. N= 4 biological replicates/genotype/timepoint. Statistics: (c & d) Students t-test (a, f & g) Two-way ANOVA. *P<0.05, **P<0.01, ***P<0.001, ****P<0.0001 Error bars represent SEM.

**Supplementary Fig. 4:**
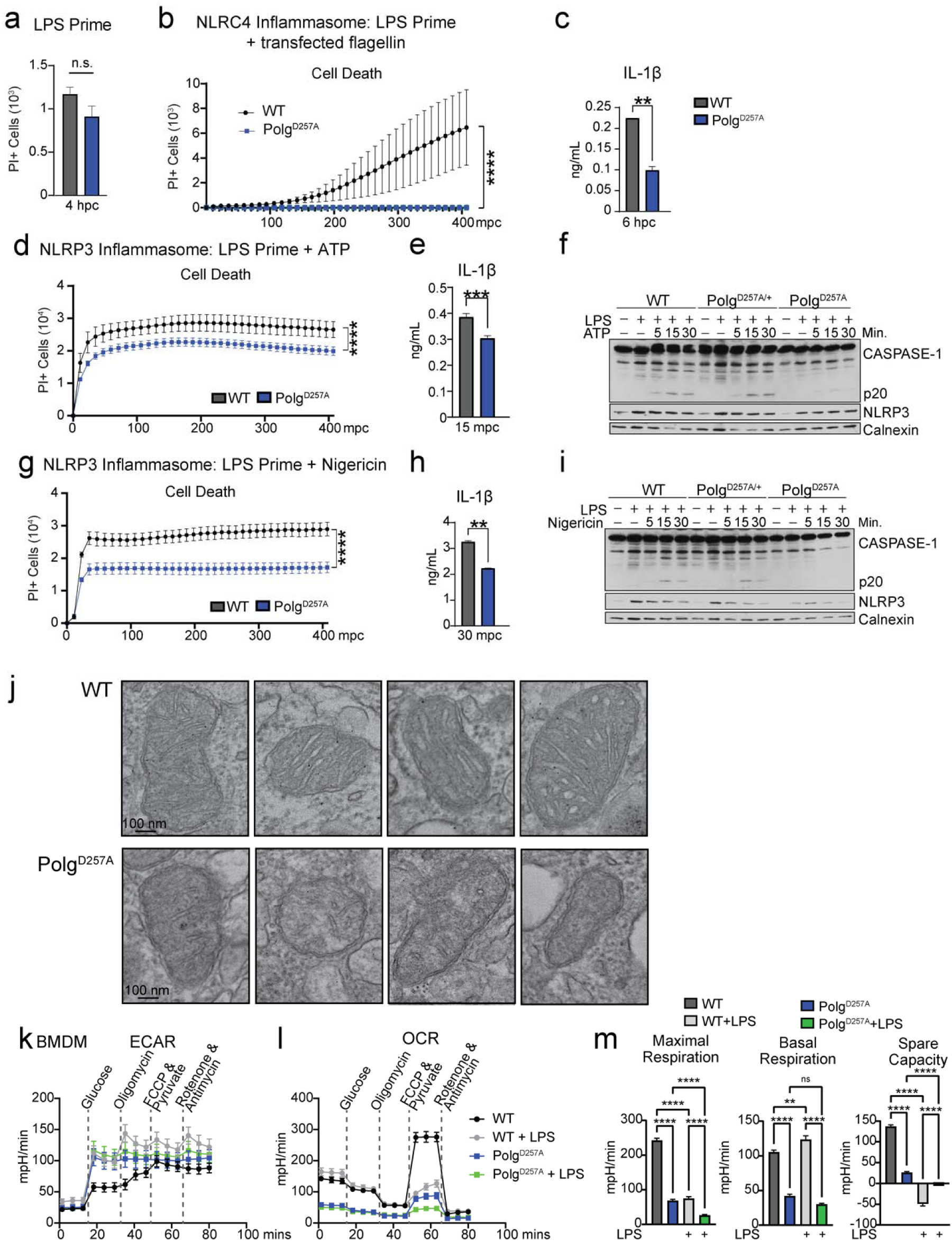
Assessment of roles for canonical inflammasome activity and metabolism in driving increased cell death and pro-inflammatory cytokine secretion in *Polg^D257A^*macrophages. **(a)** Cytation5 imaging quantification of PI positive cells in BMDMs after LPS priming for 4 hours. N= 3 biological replicates, 5 technical replicates/genotype. **(b)** Cytation5 imaging quantification of PI positive cells in BMDMs after LPS priming plus PA-flagellin transfection. N= 5 biological replicates/genotype. **(c)** ELISA analysis of IL-1β in BMDM supernatant after LPS priming plus PA-flagellin transfection. N= 2 biological replicates. **(d)** Cytation5 imaging quantification of PI positive cells in BMDMs after LPS priming plus ATP. N= 6 biological replicates/genotype. **(e)** ELISA analysis of IL-1β in BMDM supernatant after LPS priming plus ATP. N= 2 biological replicates. **(f)** Western blot of BMDM protein lysates after LPS priming plus ATP. **(g)** Cytation5 imaging quantification of PI positive cells in BMDMs after LPS priming plus nigericin. N= 6 biological replicates/genotype. **(h)** ELISA analysis of IL-1β in BMDM supernatant after LPS priming plus nigericin. N= 2 biological replicates. **(i)** Western blot of BMDM protein lysates after LPS priming plus nigericin. **(j)** Representative TEM images of mitochondria from BMDMs. **(k)** & **(l)** Seahorse analysis of BMDMs unchallenged or treated with overnight LPS. Extracellular acidification rate (ECAR) **(k)** and oxygen consumption rate (OCR) **(l)** are plotted. N=3 biological replicates and 3-4 technical replicates. **(m)** Quantification of Seahorse OCR measurements. N=3 biological replicates and 3-4 technical replicates. Statistics: (b, e, & h) Two-Way ANOVA (m) One-Way ANOVA (a, c, f, & i) Students t-test. *P<0.05, **P<0.01, ***P<0.001, ****P<0.0001 Error bars represent SEM.

**Supplementary Fig. 5:**
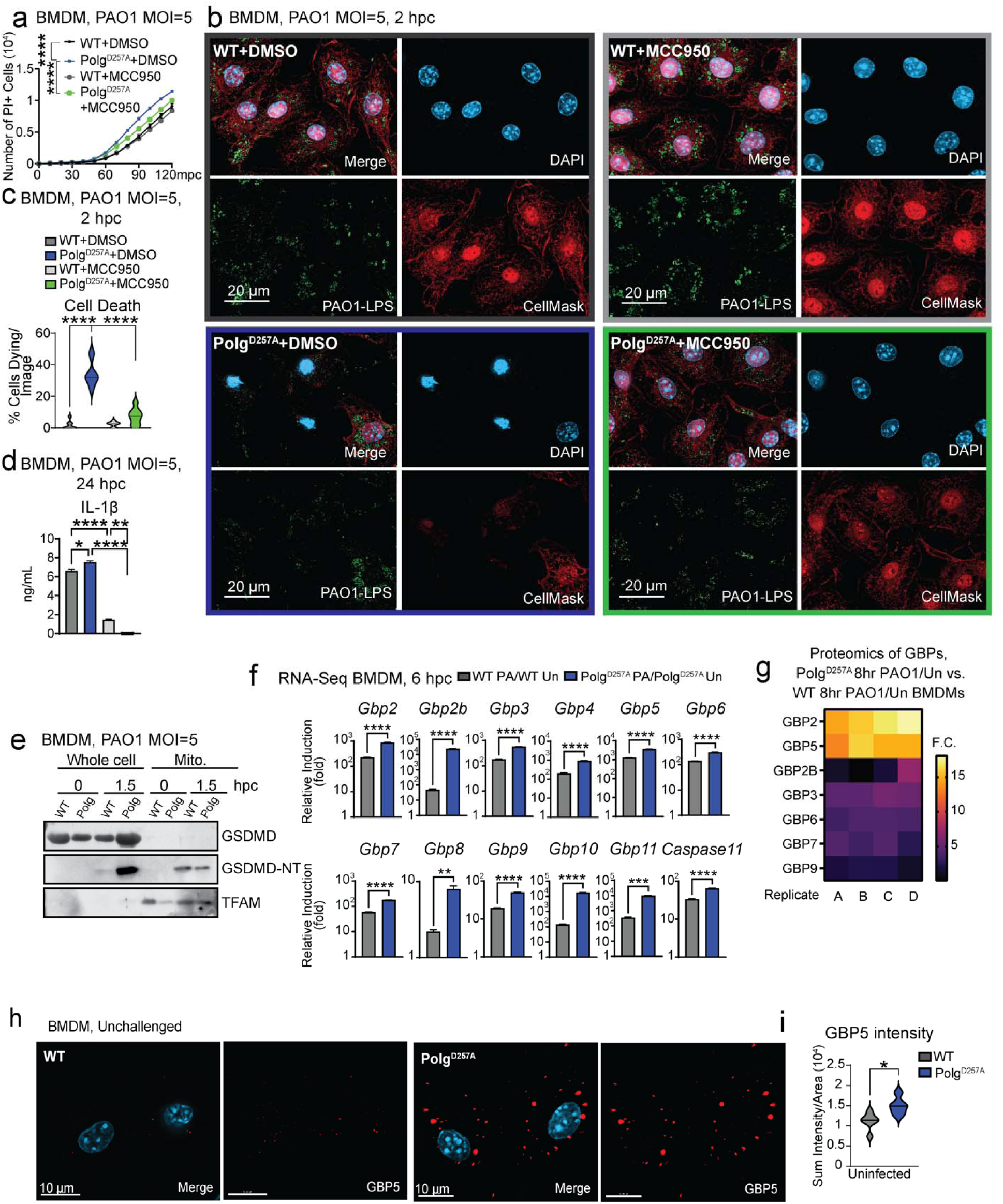
The NLRP3 inflammasome is downstream of caspase-11 activation in *Polg^D257A^*macrophages. **(a)** Cytation5 imaging quantification of PI positive cells in BMDMs with PAO1 or PAO1 plus MCC950. N= 6 biological replicates/genotype/timepoint. **(c)** Percent BMDMs dying after PAO1 and PAO1 plus NLRP3 inhibitor, MCC950, from **(b)** representative IF. N=8-9 IF images/genotype/condition. **(d)** ELISA analysis of pyroptotic cytokines in BMDM supernatant after PAO1 or PAO1 plus NLRP3 inhibitor, MCC950. N= 2 biological replicates/genotype/condition. **(e)** Western blot of GSDMD localization during PAO1 challenge. **(f)** RNA-seq from BMDMs showing induction of Gbps and Caspase-11 RNA 6 hpc with PAO1. N=3 biological replicates/genotype/condition. **(g)** GBP expression in *Polg^D257A^* (8hr PAO1/Un)/WT (8hr PAO1/Un) BMDMs by proteomics. N=4 biological replicates/genotype/condition. **(h)** Representative IF of unchallenged BMDMs stained with DAPI to mark the nucleus and anti-GBP5 antibody. **(i)** Sum intensity of GBP5 per cell area in resting BMDMs in (**h**) IF. N=5-6 images/genotype Statistics: (f, & i) Student’s t-test (c, & d) One-way ANOVA (a) Two-way ANOVA. *P<0.05, **P<0.01, ***P<0.001, ****P<0.0001 Error bars represent SEM.

**Supplemental Fig. 6:**
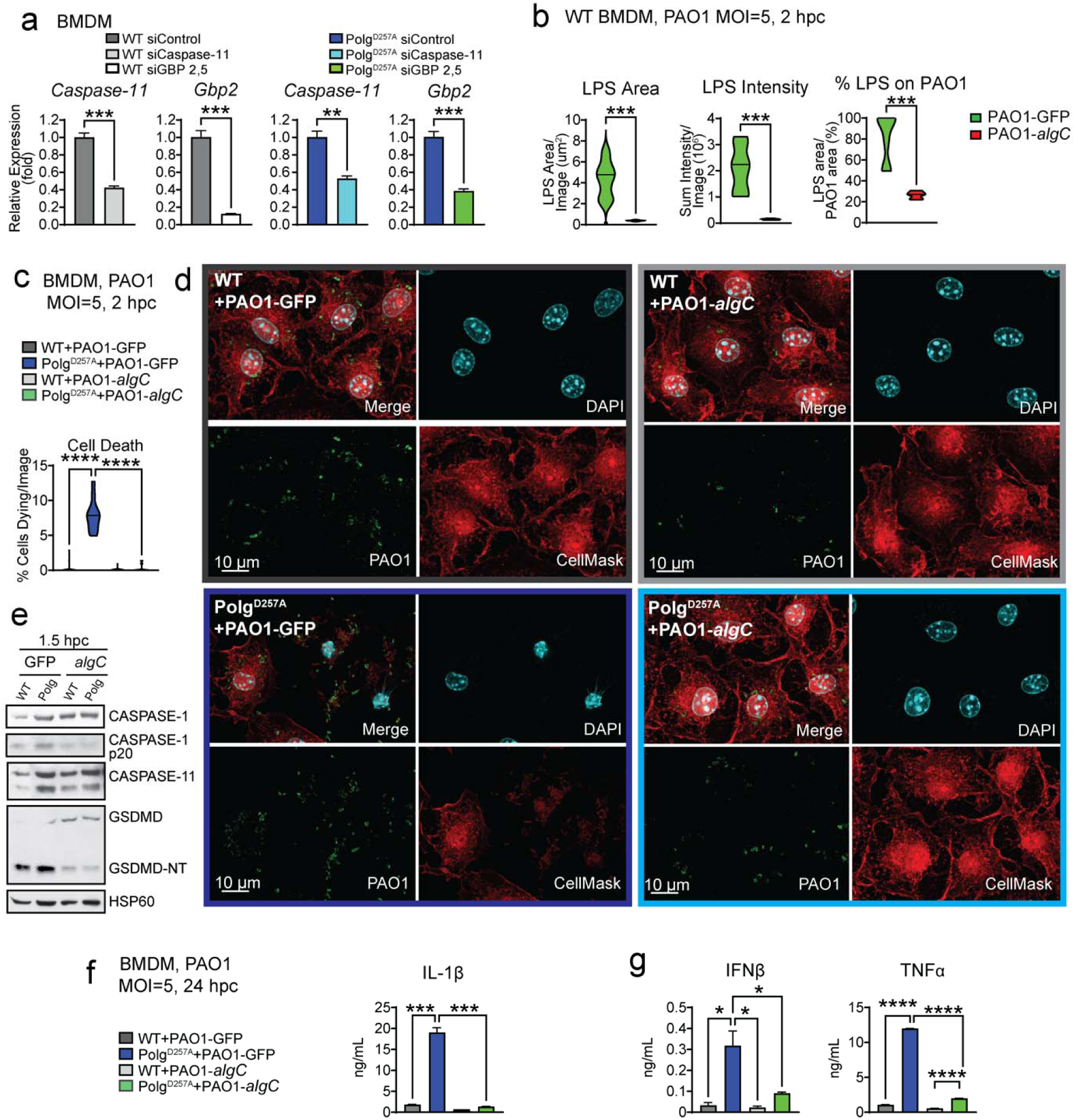
The PAO1 *algC* mutant displays decreased levels of LPS and does not robustly engage the caspase-11 inflammasome or induce potentiated cell death and hyperinflammation in *Polg^D257A^* macrophages. **(a)** qRT-PCR quantification of siRNA knock down efficiency in BMDMs. N=3 technical replicates/genotype. **(b)** Quantification of LPS associated with PAO1 and total LPS staining in BMDMs after PAO1 or PAO1-*algC* challenge by IF. N=6-7 IF images/genotype/condition. **(c)** Percent BMDMs dying after PAO1 and PAO1-*algC* from **(d)** representative IF. N=10 IF images/genotype/condition. **(e)** Western blotting of proteins involved in pyroptosis in BMDMs after PAO1 or PAO1-*algC* challenge. **(f)** ELISA analysis of IL-1 cytokines in BMDM supernatant after PAO1 or PAO1-*algC*. N= 2 biological replicates/genotype/condition. **(g)** ELISA analysis of proinflammatory cytokines in BMDM supernatant after PAO1 or PAO1-*algC*. N= 2 biological replicates/genotype/condition. Statistics: (a & b) Student’s t-test (c, f, & g) One-way ANOVA. *P<0.05, **P<0.01, ***P<0.001, ****P<0.0001 Error bars represent SEM.

**Supplemental Fig. 7:**
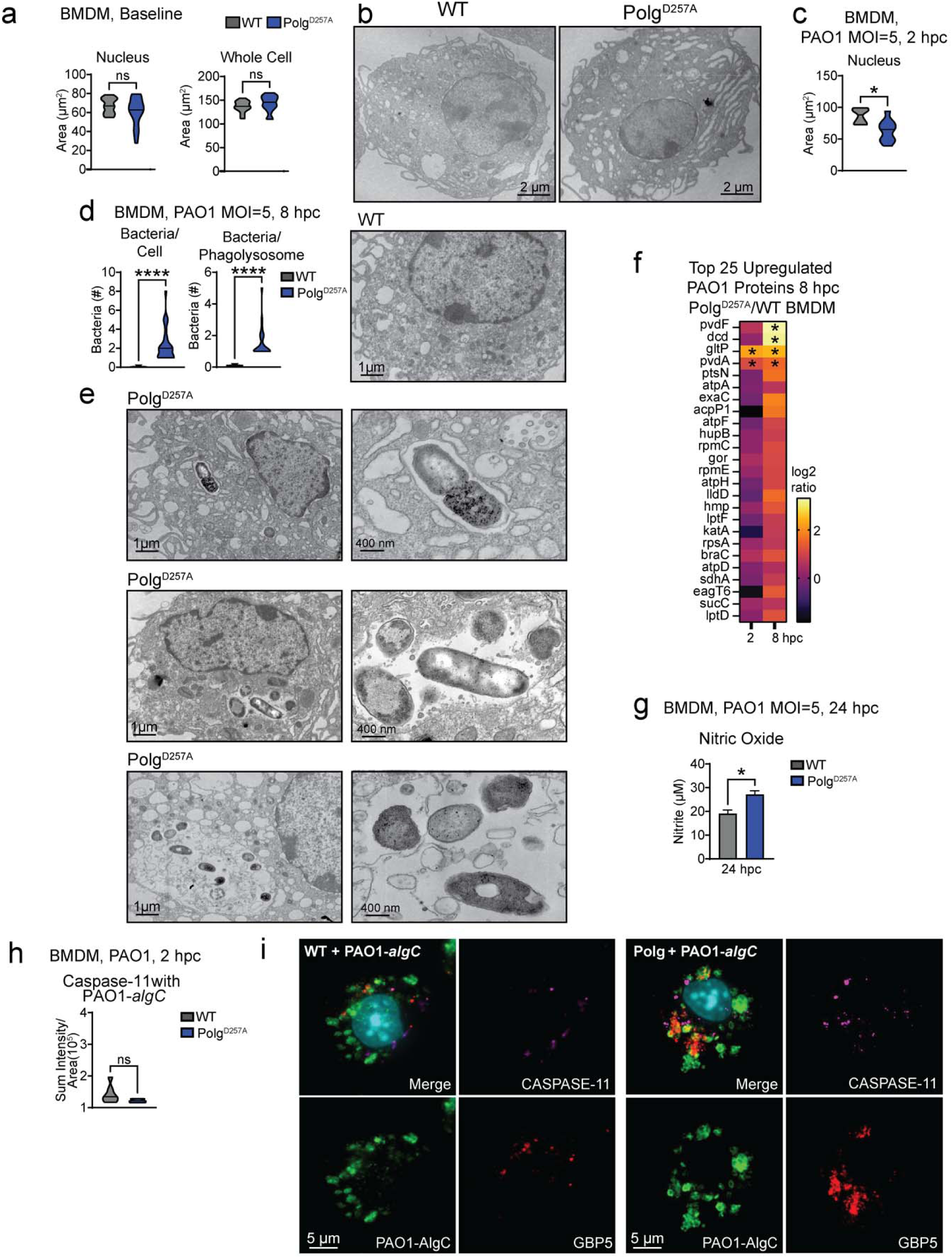
*Polg^D257A^*macrophages have more PAO1 and PAO1 proteins present at late timepoints. **(a)** Quantitation of nucleus and cellular area of BMDMs from **(b)** TEM images of BMDMs at rest. N= 14-15 images/condition. **(c)** Quantitation of nuclear area of BMDMs from (Fig. 6b) TEM images. N= 9 images/condition. **(d)** Quantification of bacteria in BMDMs and their lysosomes 8 hours after PAO1 by **(e)** TEM. N=15-20 cells/genotype. **(f)** Proteomic analysis of PAO1 proteins in *Polg^D257A^* homozygous mutant BMDMs compared to WT 8 hours after PAO1 challenge. N=4 biological replicates/genotype/condition. **(g)** Quantification of nitric oxide in BMDMs 24 hpc with PAO1. N=3 biological replicates/genotype. **(h)** Quantification of capase-11 associated with PAO1-GFP or PAO1-*algC* mutant from representative IF images **(i)**. N=6 IF images/genotype/condition. Statistics: (a, c, d, g, & h) Students t-test. *P<0.05, **P<0.01, ***P<0.001, ****P<0.0001 Error bars represent SEM.

**Supplemental Fig. 8:**
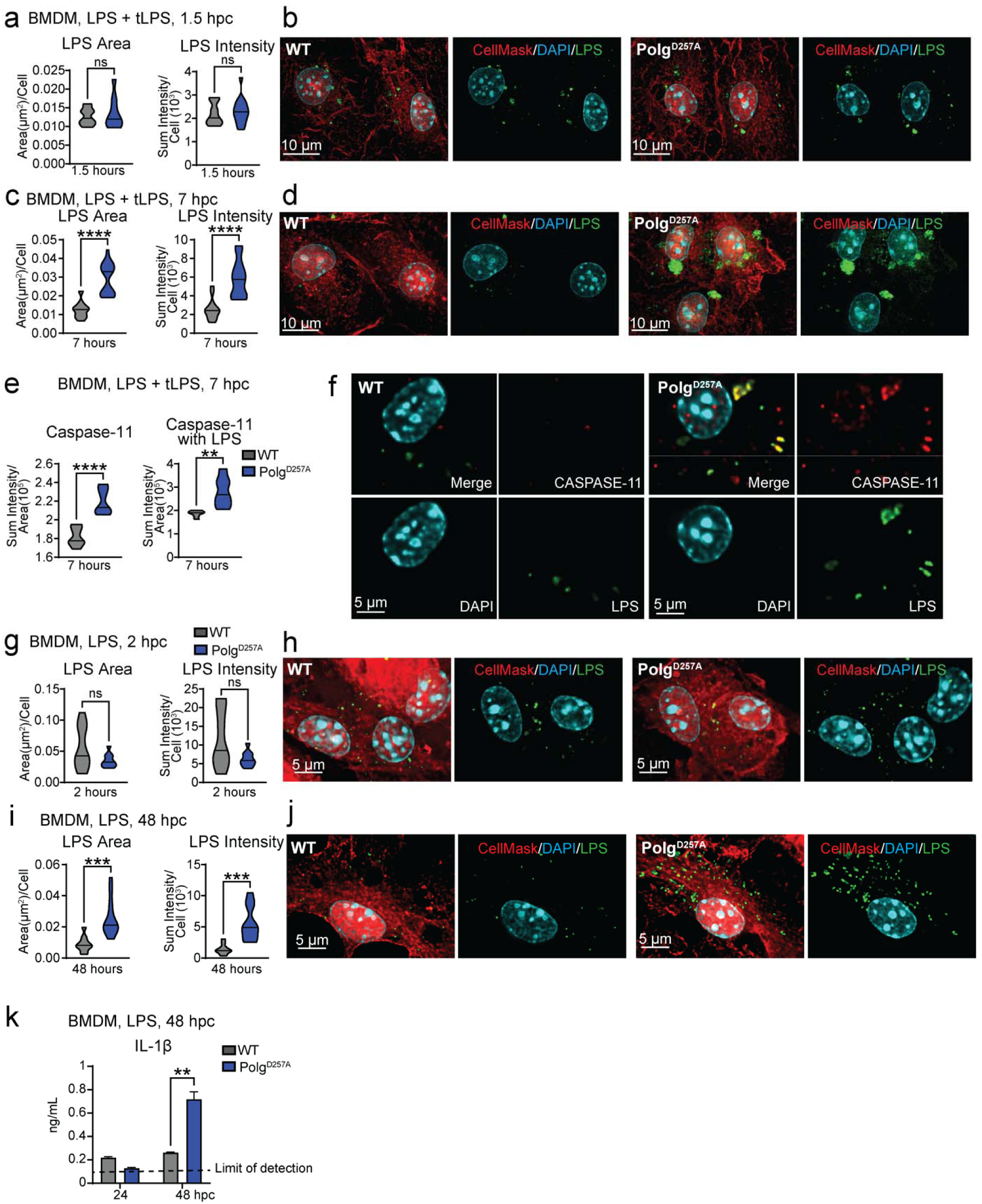
*Polg^D257A^*macrophages uptake similar levels of LPS but maintain higher levels of LPS and more caspase-11 association with LPS at later challenge timepoints. **(a)** Quantification of LPS per cell in BMDMs after 4 hours LPS priming then 1.5 hours LPS transfection from **(b)** IF. N=10-11 IF images/genotype/condition. **(c)** Quantification of LPS per cell in BMDMs after 4 hours LPS priming then 7 hours LPS transfection from **(d)** IF. N=10-12 IF images/genotype/condition. **(e)** Quantification of caspase-11 intensity and caspase-11 associated with LPS in BMDMs after 4 hours LPS priming then 7 hours LPS transfection from **(f)** IF. N=7 IF images/genotype/condition. **(g)** Quantification of LPS per cell in BMDMs 2 hours post LPS challenge from **(h)** IF. N=8-9 IF images/genotype/condition. **(i)** Quantification of LPS per cell in BMDMs 48 hours post LPS challenge from **(j)** IF. N=10 IF images/genotype/condition. **(k)** ELISA analysis of IL-1β in BMDM supernatant after LPS. N= 2 biological replicates/genotype/timepoint. Statistics: (a, c, e, & g) Students t-test (k) Two-way ANOVA. *P<0.05, **P<0.01, ***P<0.001, ****P<0.0001 Error bars represent SEM.

**Supplementary Fig. 9:**
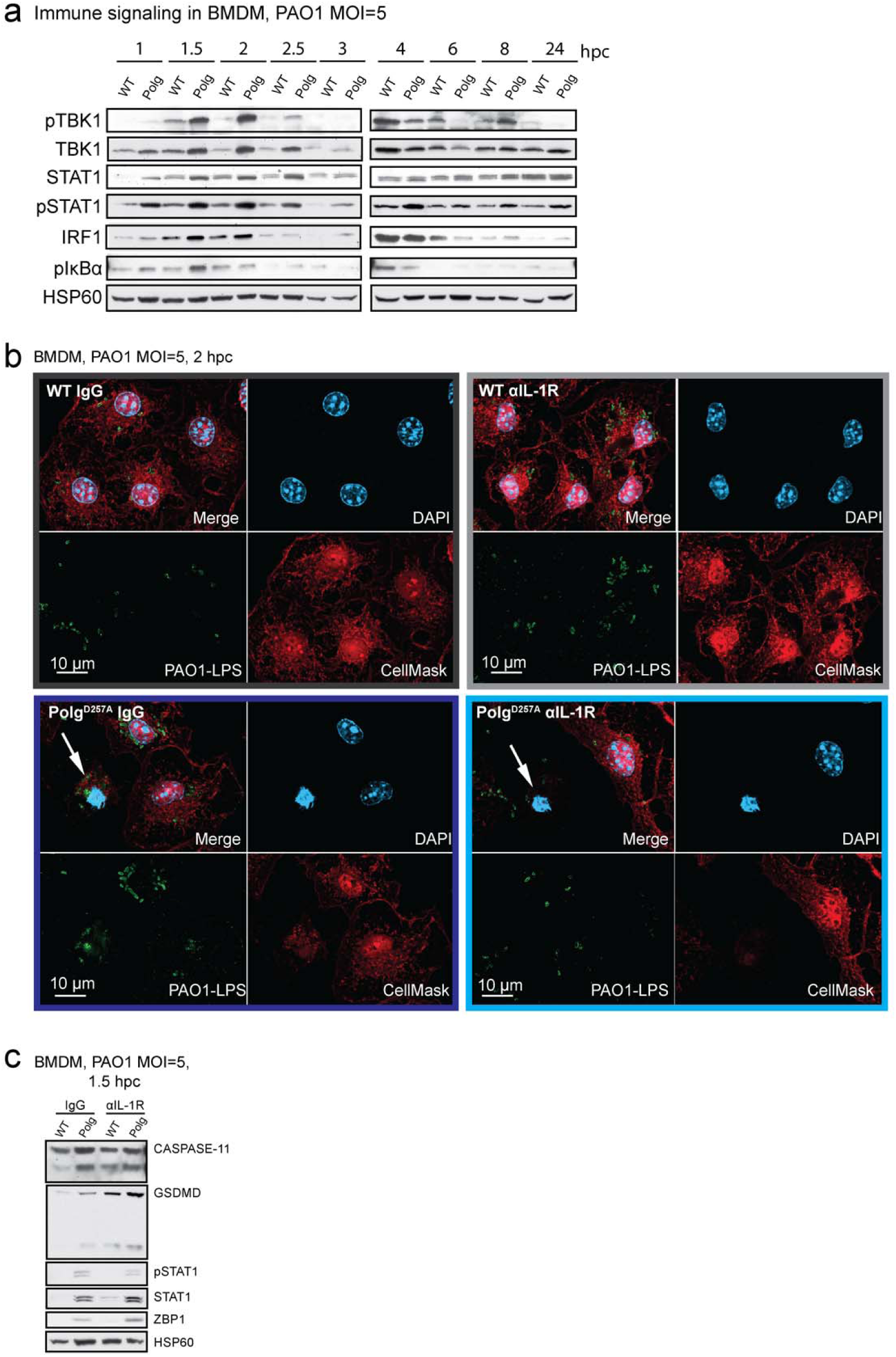
*Polg^D257A^* BMDMs exhibit enhanced NF-κB signaling that is downstream of the IL-1R when challenged with PAO1. **(a)** Western blot of signaling in BMDMs during a PAO1 time course. **(b)** Representative images of cell death quantitation in Fig. 5f. **(c)** Western blot analysis of lysates from BMDMs challenged with PAO1 and PAO1 plus anti-IL-1R blockade 1.5 hpc.

**Supplemental Fig. 10:**
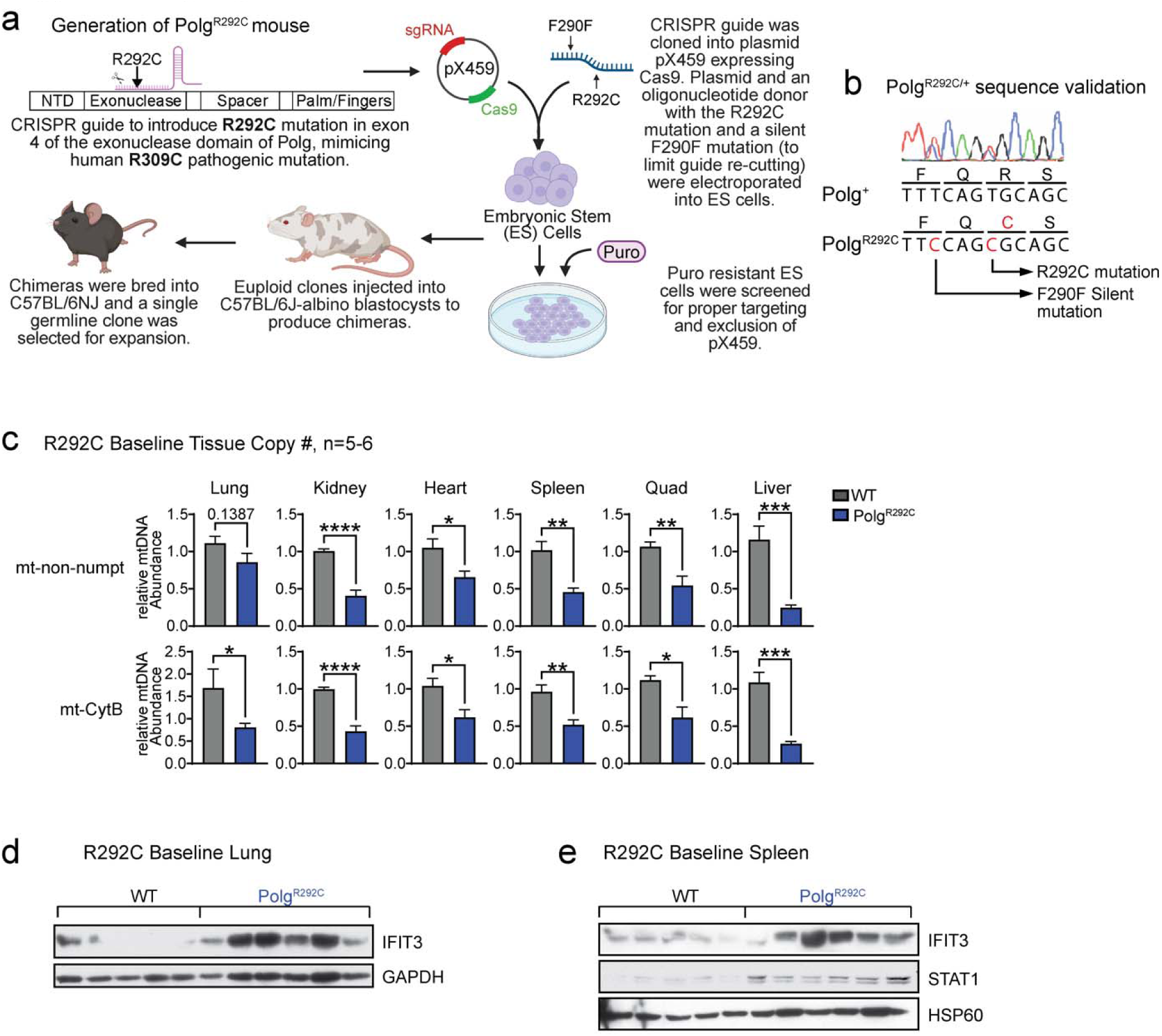
Mice incorporating a deleterious patient mutation (*Polg^R292C^*) exhibit mtDNA depletion and elevated IFN-I responses. **(a)** Generation of the *Polg^R292C^* mouse. **(b)** Chromatograph from Sanger sequencing of DNA from heterozygous *Polg^R292C^* mouse verifying germline incorporation of mutation. **(c)** qPCR analysis of mtDNA abundance in tissues. N= 4-6 mice/genotype. **(d)** Western blot analysis of ISG, IFIT3, in lung lysates from *Polg^R292C^* mice. **(e)** Western blot analysis of ISGs in spleen lysates from *Polg^R292C^* mice. Statistics: (c) Student’s t-test. *P<0.05, **P<0.01, ***P<0.001, ****P<0.0001 Error bars represent SEM.

**Supplemental Fig. 11:**
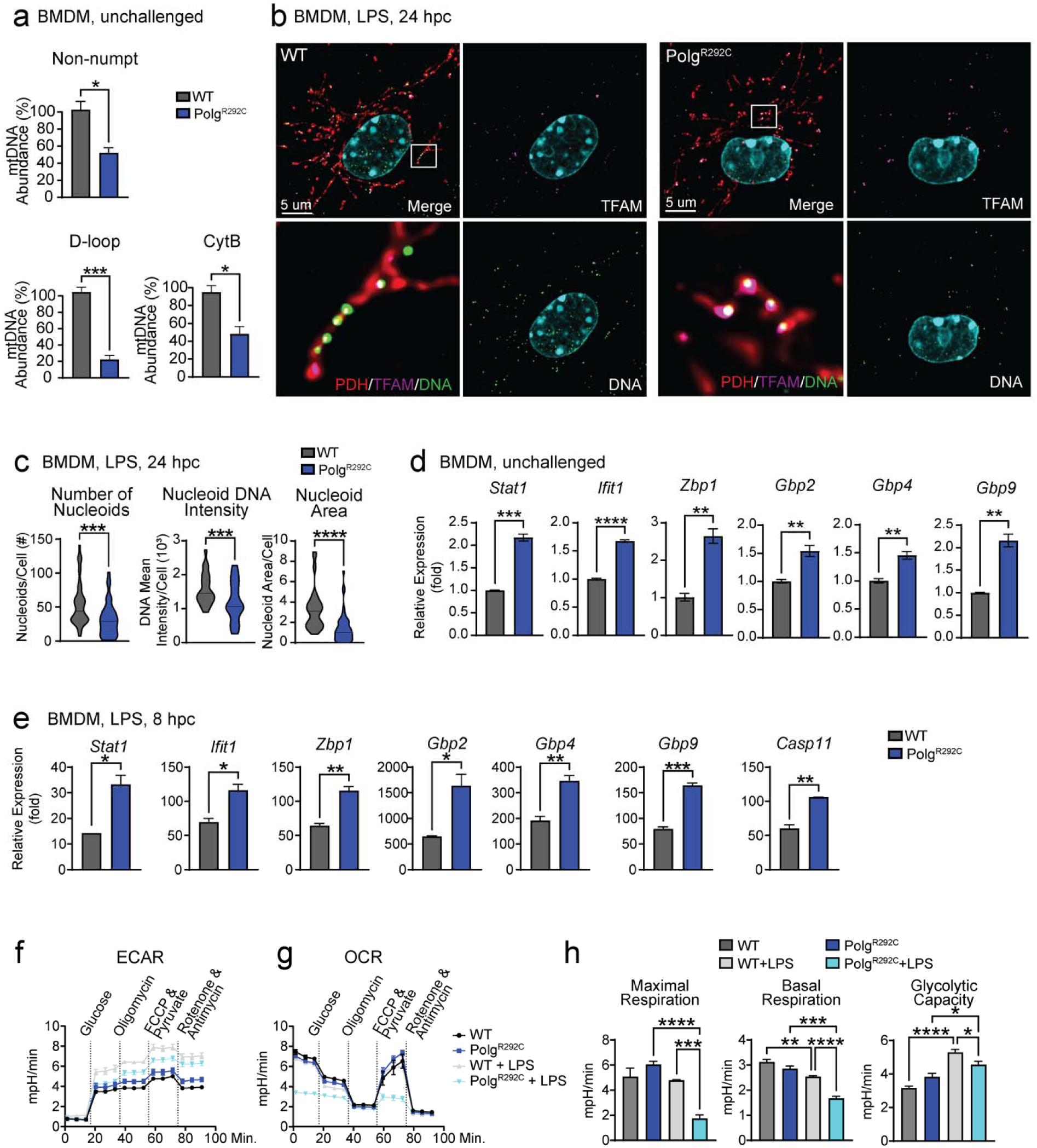
Macrophages incorporating a deleterious patient mutation (*Polg^R292C^*) exhibit mtDNA depletion, elevated IFN-I responses, and altered metabolism. **(a)** qPCR analysis of mtDNA abundance in BMDMs. N= 3 technical triplicates/genotype. **(b)** Representative IF of nucleoids in BMDMs stained with anti-PDH antibody to label mitochondria, anti-DNA antibody to label nucleoids, and anti-TFAM antibody. **(c)** Quantification of nucleoids per cell, mean intensity of DNA signal per nucleoid, and nucleoid area per cell from BMDMs in **(b)** IF. N=40-50 cells/genotype **(d)** qRT-PCR analysis of ISGs in unchallenged *Polg^R292C^* BMDMs. N=3 biological replicates, 3 technical replicates/genotype. **(e)** qRT-PCR analysis of ISGs in *Polg^R292C^* BMDMs 8 hours post challenge (hpc) with LPS. N= 3 technical replicates/genotype. **(f)** & **(g)** Seahorse analysis of BMDMs unchallenged or treated with overnight LPS. Extracellular acidification rate (ECAR) **(f)** and oxygen consumption rate (OCR) **(g)** are plotted. N=2 biological replicates and 5-6 technical replicates. **(h)** Quantification of Seahorse OCR measurements. N=2 biological replicates and 4-5 technical replicates. Statistics: (a, c, d, & e) Students t-test (h) One-Way ANOVA. *P<0.05, **P<0.01, ***P<0.001, ****P<0.0001 Error bars represent SEM.

**Supplemental Fig. 12:**
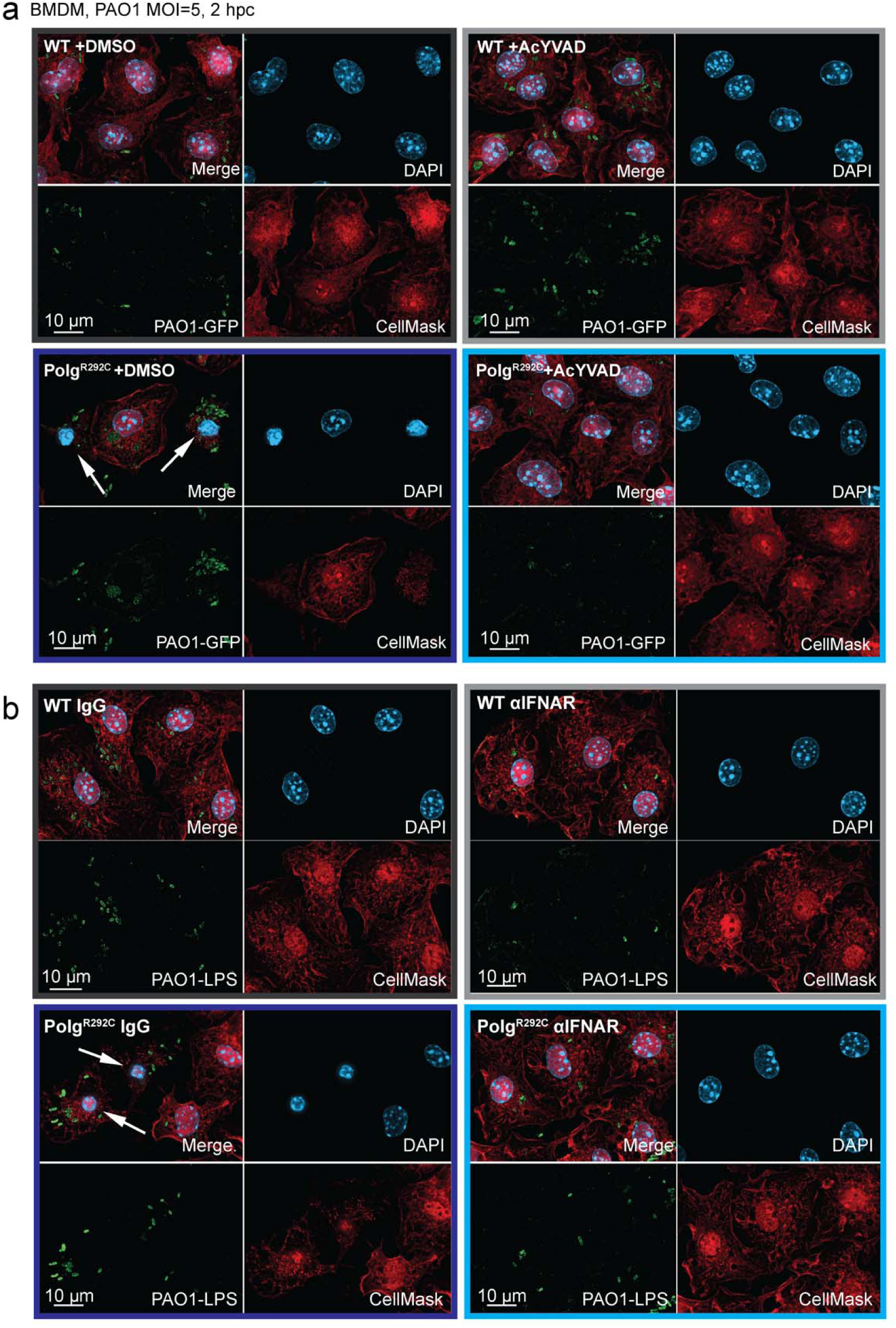
AcYVAD and anti-IFNAR blockade reduces cell death in *Polg^R292C^* macrophages. **(a)** Representative IF of dying cells characterized by condensed nuclei and loss of cell mask staining (white arrows) after PAO1 and PAO1 plus caspase-1/11 inhibitor, AcYVAD, quantified in Fig. 6c. **(b)** Representative IF of dying cells characterized by condensed nuclei and loss of cell mask staining (white arrows) after PAO1 and PAO1 plus anti-IFNAR blockade quantified in Fig. 6f.

**Supplementary Fig. 13:**
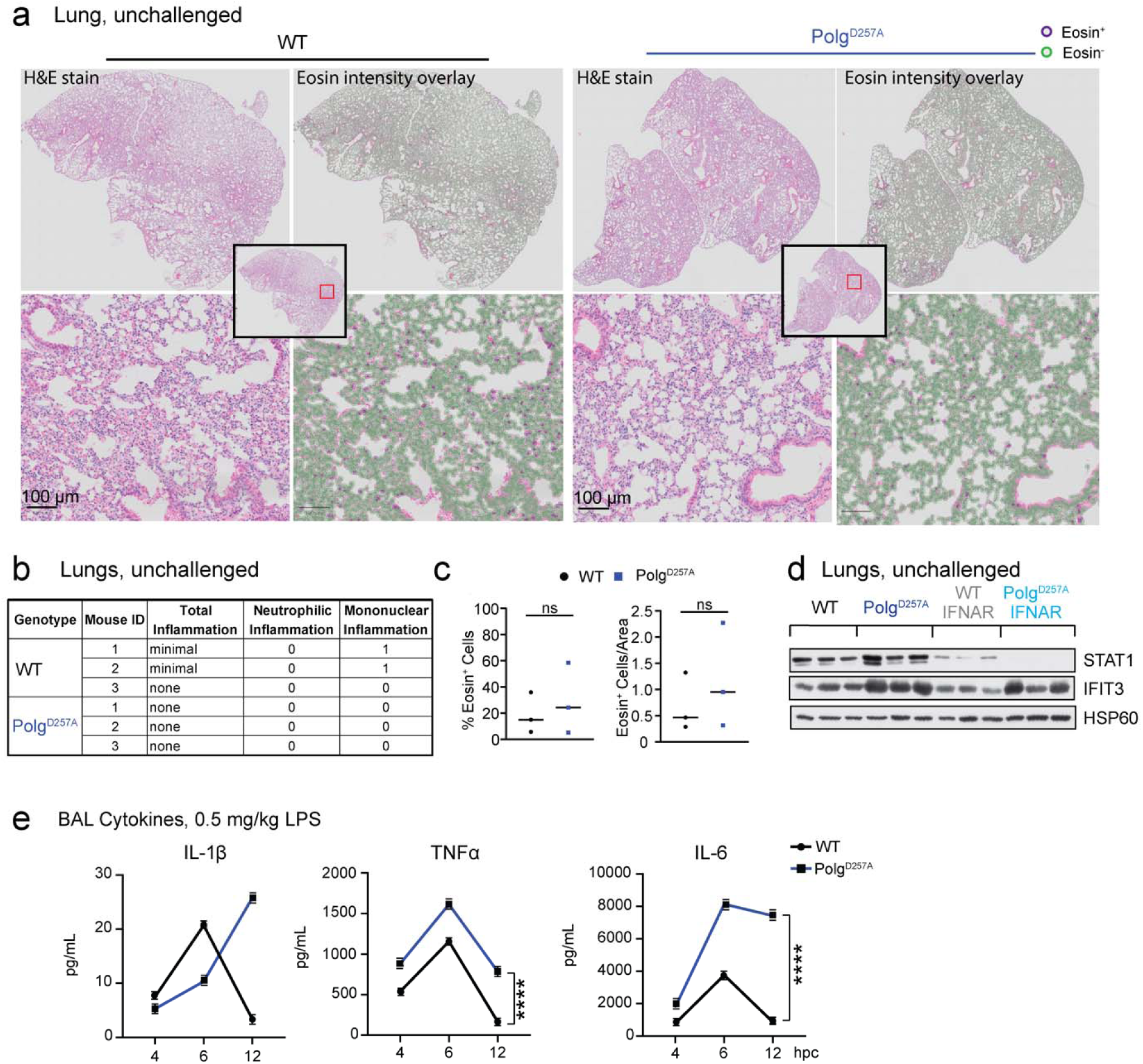
*Polg* mutant mice exhibit elevated IFN-I and inflammatory responses to LPS, but do not exhibit basal differences in lung morphology or immune cell abundance. **(a)** Representative histology images of hematoxylin and eosin (H&E) staining of WT and *Polg^D257A^* lungs, with an overlay of eosin positive cells using QuPath software. **(b)** Pathology assessment of unchallenged lungs. **(c)** QuPath analysis of eosin^+^ cells in the lungs from **(c)**. N=3 mice/genotype **(d)** Protein expression of ISGs in unchallenged WT and *Polg^D257A^* lungs homogenates. **(e)** ELISA analysis of inflammatory cytokines in bronchioalveolar lavage of LPS challenged WT and *Polg^D257A^*lungs. N=2 mice/genotype/timepoint. Statistics: (c) Students t-test (e) Two-Way ANOVA. *P<0.05, **P<0.01, ***P<0.001, ****P<0.0001 Error bars represent SEM.

**Supplementary Fig. 14:**
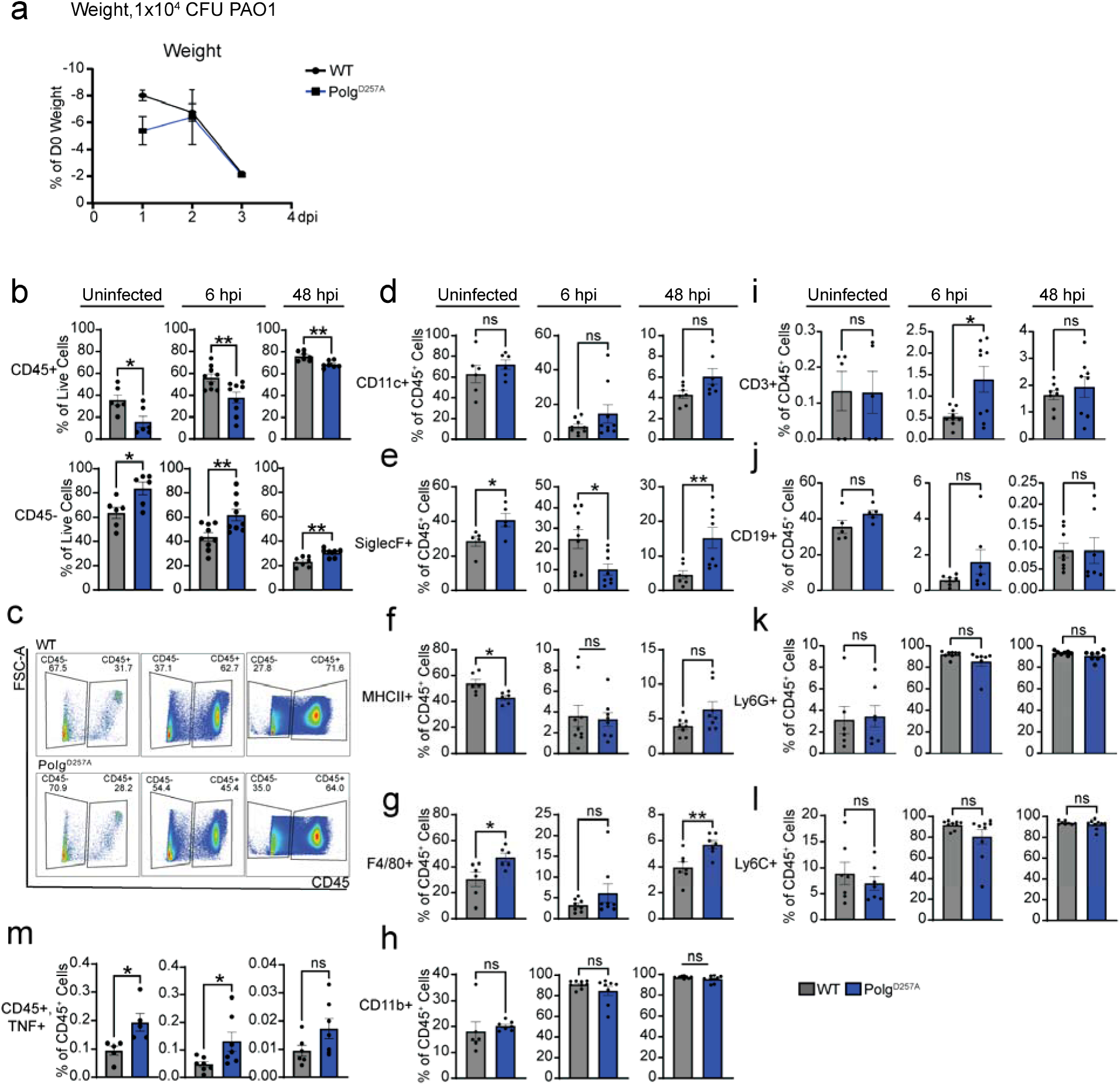
Intratracheal instillation of PAO1 alters immune cell composition in *Polg* mutant mouse lungs. **(a)** Weight change in WT and *Polg^D257A^* mice after infection with 1×10^4^ PAO1. N=4 mice/genotype. **(b)** CD45 expression on cells from bronchioalveolar lavage (BAL) from uninfected and PAO1 infected WT and *Polg^D257A^* mice. N=5-9 mice/genotype/timepoint. **(c)** Gates used to quantify CD45 expression in **(b)**. **(d)** Percent of CD45^+^ cells expressing CD11c from the BAL of uninfected and PAO1 infected lungs. N=5-9 mice/genotype/timepoint. **(e)** Percent of CD45^+^ cells expressing SiglecF from the BAL of uninfected and PAO1 infected lungs. N=5-9 mice/genotype/timepoint. **(f)** Percent of CD45^+^ cells expressing MHCII from the BAL of uninfected and PAO1 infected lungs. N=5-9 mice/genotype/timepoint. **(g)** Percent of CD45^+^ cells expressing F4/80 from the BAL of uninfected and PAO1 infected lungs. N=5-9 mice/genotype/timepoint. **(h)** Percent of CD45^+^ cells expressing CD11b from the BAL of uninfected and PAO1 infected lungs. N=5-9 mice/genotype/timepoint. **(i)** Percent of CD45^+^, CD11b^-^ cells expressing CD3 from the BAL of uninfected and PAO1 infected lungs. N=5-9 mice/genotype/timepoint. **(j)** Percent of CD45^+^, CD11b^-^ cells expressing CD19 from the BAL of uninfected and PAO1 infected lungs. N=5-9 mice/genotype/timepoint **(k)** Percent of CD45^+^ expressing Ly6G from the BAL of uninfected and PAO1 infected lungs. N=5-9 mice/genotype/timepoint. **(l)** Percent of CD45^+^ cells expressing Ly6C from the BAL of uninfected and PAO1 infected lungs. N=5-9 mice/genotype/timepoint. **(m)** CD45^+^ cells with cell associated TNFα expression from BAL of uninfected and PAO1 infected lungs. N=5-9 mice/genotype/timepoint. Statistics: (a) Two-way ANOVA (b, & d-m) Students t-test. *P<0.05, **P<0.01, ***P<0.001, ****P<0.0001 Error bars represent SEM.

**Supplementary Fig. 15:**
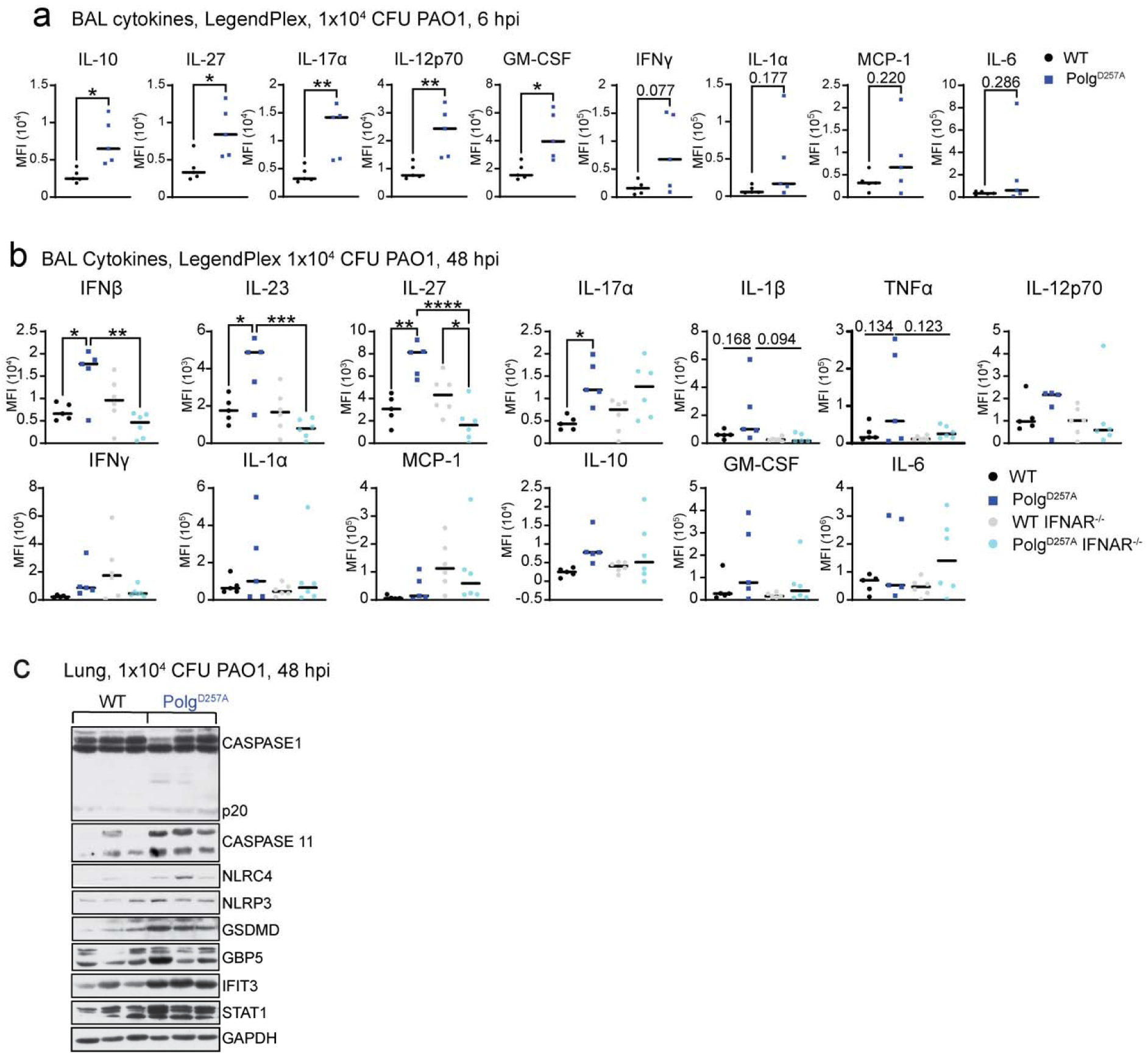
A non-lethal dose of PAO1 induces hyperinflammation and activation of pyroptosis in the lungs of *Polg^D257A^* mice. **(a)** LegendPlex pro-inflammatory cytokine analysis of bronchial alveolar lavage (BAL) 6 hours post infection (hpi) with 1×10^4^ colony forming units (CFU) of PAO1. N=5 mice/genotype **(b)** LegendPlex pro-inflammatory cytokine analysis of bronchioalveolar lavage (BAL) from WT, *Polg^D257A^*, WT IFNAR^-/-^, and *Polg^D257A^* IFNAR^-/-^ lungs infected with 1×10^4^ PAO1 for 48 hours. All undefined comparisons are not significant. N=5-6 mice/genotype. **(c)** Western blot of ISGs and proteins involved in pyroptosis from homogenized lung tissue from mice infected with 1×10^4^ PAO1 for 48 hours. Statistics: (a) Students t-test (b) One-way ANOVA. *P<0.05, **P<0.01, ***P<0.001, ****P<0.0001 Error bars represent SEM.

**Supplementary Fig. 16:**
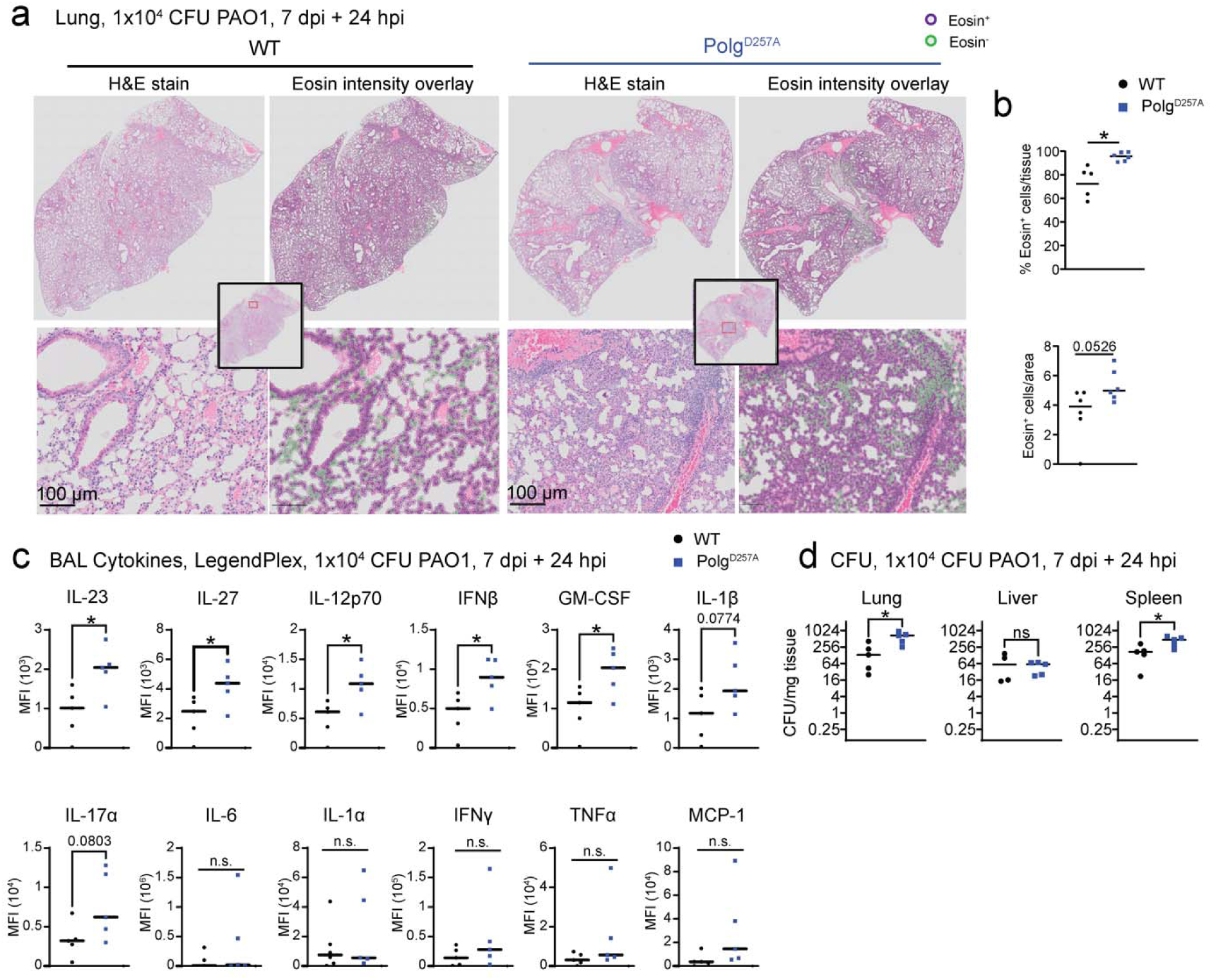
*Polg^D257A^*mice exhibit hyperinflammation in response to infection and reinfection with PAO1. **(a)** Representative histology images of hematoxylin and eosin (H&E) staining of WT and *Polg^D257A^*lungs after initial infection with 1×10^4^ PAO1, followed by 7 days rest, then reinfection with 1×10^4^ PAO1. Overlay of eosin positive cells was done using QuPath software. **(b)** QuPath analysis of eosin^+^ cells in the lung from **(a)** histology. N=6 mice/genotype. **(c)** LegendPlex pro-inflammatory cytokine analysis of BAL from WT and *Polg^D257A^* lungs after initial infection with 1×10^4^ PAO1, followed by 7 days rest, then reinfection with 1×10^4^ PAO1. N=5 mice/genotype. **(d)** Colony counts from homogenized tissue after initial infection with 1×10^4^ PAO1, followed by 7 days rest, then reinfection with 1×10^4^ PAO1. N=5 mice/genotype. Statistics: (b, c, & d) Students t-test. *P<0.05, **P<0.01, ***P<0.001, ****P<0.0001. Error bars represent SEM.

**Supplementary Fig. 17:**
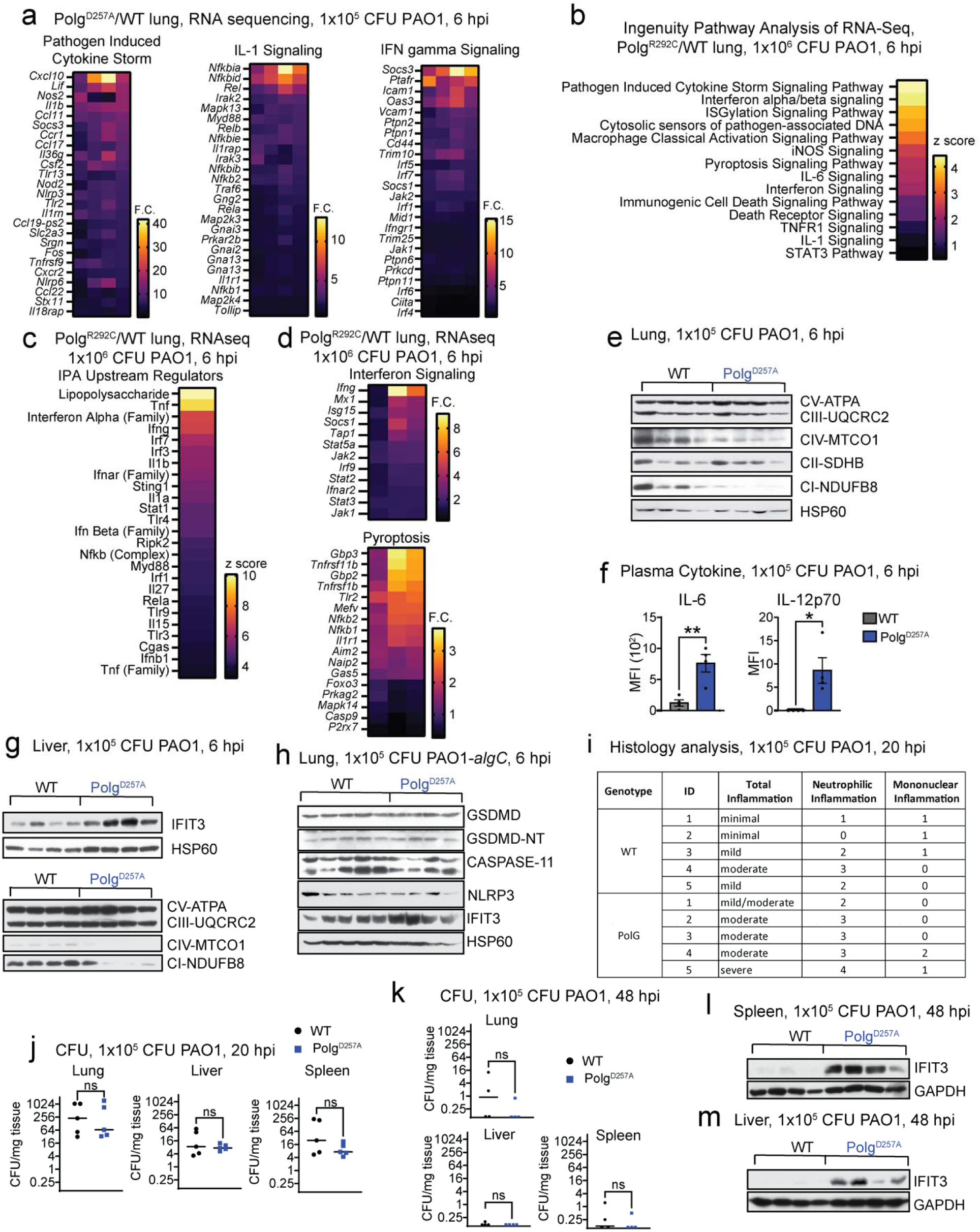
*Polg* mutant mice exhibit increased IFN-I, pyroptotic cell death, and pro-inflammatory responses after challenge with PAO1. **(a)** Fold changes of representative genes in IPA pathways from Fig. 7b in *Polg^D257A^* lungs compared to WT after LPS or PAO1. **(b)** Ingenuity Pathway Analysis (IPA) of RNAseq from lungs showing Z-scores of pathways in *Polg^R292C^*lungs compared to WT after PAO1 challenge. N=3 biological replicates/genotype/condition. **(c)** IPA Upstream Regulator analysis showing Z-scores in *Polg^R292C^* lung compared to WT after PAO1 challenge. **(d)** Log^2^ fold changes of representative genes in IPA pathways from (**b**) in *Polg^R292C^*lungs compared to WT after PAO1. **(e)** Western blot of proteins involved in oxidative phosphorylation in the lung 6 hours post infection (hpi) with 1×10^5^ colony forming units (CFU) of PAO1. **(f)** LegendPlex of circulating pro-inflammatory cytokine in the plasma 6 hpi with 1×10^5^ CFU of PAO1. N=4 mice/genotype **(g)** Western blot of ISG and proteins involved in oxidative phosphorylation in the liver 6 hpi with 1×10^5^ CFU of PAO1. **(h)** Western blot of ISG and proteins involved in pyroptosis in the lung 6 hours post infection with 1×10^5^ CFU of PAO1-*algC*. **(i)** Summary of pathologist scoring of H&E stained lungs infected with 1×10^5^ CFU PAO1 at 20 hpi. N=5 mice/genotype. **(j)** Colony counts from homogenized tissue infected with 1×10^5^ CFU PAO1 at 20 hpi. N=5 mice/genotype. **(k)** Colony counts from homogenized tissue infected with 1×10^5^ CFU PAO1 at 48 hpi. N=4 mice/genotype. **(l)** Western blot of ISGs from homogenized liver and **(m)** spleen from mice infected with 1×10^5^ CFU PAO1 at 48 hpi. Statistics: (f, j, & k) Students t-test. *P<0.05, **P<0.01, ***P<0.001, ****P<0.0001. Error bars represent SEM.

**Supplementary Fig. 18:**
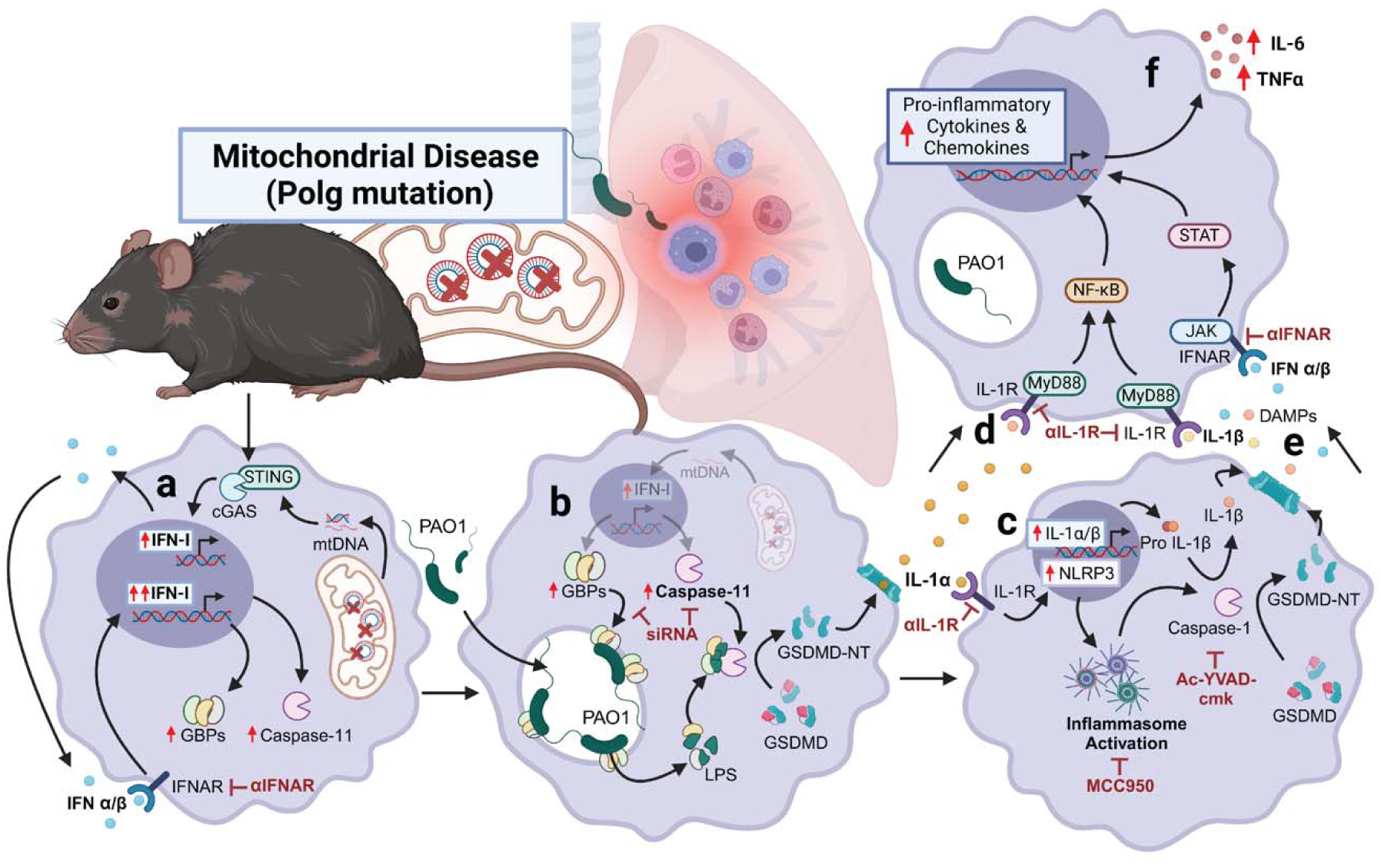
Elevated IFN-I primes macrophages to undergo caspase-11 non-canonical inflammasome activation and drives hyperinflammatory signaling, resulting in lung injury in mice with *Polg*-related mitochondrial dysfunction. Mitochondrial DNA instability in exonuclease-deficient *Polg* mutant mice triggers chronic IFN-I signaling that enhances expression of caspase-11, GBPs, and other ISGs **(a)**. Inhibitors and genetic approaches (maroon arrows) revealed that GBPs accumulate on bacteria and augment activation of the caspase-11 non-canonical inflammasome, resulting in pyroptosis and the GSDMD-dependent release of IL-1α **(b)**. IL-1α signals through the IL-1R to further upregulate the expression of inflammasome mediators including NLRP3 and pro-IL-1β **(c)**, leading to the release of mature IL-1β **(d)** and DAMPs **(e)** that feed forward to sustain and enhance IFN-I and IL-1R signaling, elevating release of other pro-inflammatory cytokines such as TNFα and IL-6 **(f)**. Elevated rates of pyroptotic cell death and hyperinflammation in the lung of *Polg* mutant mice contribute to acute lung injury and systemic inflammation resulting in decreased survival after infection.

## Acknowledgements

We thank Robbie Moore at the Texas A&M College of Medicine Analytical Cytometry Core and Will Schott and Danielle Littlefield in the JAX Flow Cytometry Service for technical assistance. We thank Pete Finger in the JAX Histopathology and Microscopy Sciences Service for assistance preparing macrophage samples for TEM imaging. We acknowledge the JAX Center for Precision Genetics (JCPG) for generating the *Polg^R292C/R292C^* mouse line. We would like to acknowledge the Mass Spectrometry and Protein Chemistry Service within Protein Sciences at The Jackson Laboratory for their guidance and support with the mass spectrometry analyses performed in this study. Finally, we thank Vijay Rathinam for protocols and helpful discussions on non-canonical inflammasome biology.

## Author contributions

A.P.W and J.J.V. designed experiments, analyzed data, and wrote the manuscript. Y.L. assisted with animal experiments and sample processing. M.R. assisted with sample processing. C.G.M. performed inflammasome challenge experiments. J.W. assisted with pathology scoring of histology samples. T.J.S and B.R.H. performed proteomics and analysis. C.P. and R.S. maintained mouse lines and generated BMDMs. S.C.K. and S.A.M. generated the *Polg^R292C^*mouse model and provided samples. P.J.M. and C.L.C. assisted with data analysis and interpretation. A.P.W. conceived the project and provided overall direction.

## Data and materials availability

All data needed to evaluate the conclusions drawn herein are present in the paper and Supplementary Materials. Raw data will be provided upon request. The RNA sequencing and proteomic datasets generated through this work will be deposited in the Gene Expression Omnibus or PRIDE database, respectively, after acceptance of the paper for publication.

## Methods

### Mouse Husbandry and Strains

C57BL/6J (JAX Stock # 000664), *Polg^D257A/D257A^* (JAX Stock # 017341), and *Ifnar1^-/-^* (JAX Stock # 028256) mice were obtained from The Jackson Laboratory and bred and maintained in multiple vivaria at Texas A&M University and The Jackson Laboratory. Experimental *Polg^+/+^* (WT) and *Polg^D257A/D257A^* littermates were obtained from *Polg^D257A/+^*maintenance colony crosses of male *Polg^D257A/+^* to female *Polg^+/+^*. This breeding strategy was adopted to limit the accumulation and maternal transmission of mutant mtDNA molecules to heterozygous breeders. *Polg^D257A/D257A^*mice and littermate controls used for macrophage experiments were aged for 12-16 months. *Polg^D257A/D257A^* mice and littermate controls used for in vivo infections were aged from 7-12 months.

The *Polg^R292C/R292C^* mouse line (B6N(Cg)-*Polg^em5(F290F,R292C)Lutzy/Mmjax^*) was generated using CRISPR/Cas9 endonuclease-mediated genome editing in mouse embryonic stem cells (mESCs). A single guide RNA (CTCCCATCCACAGGCTGCGC) targeting exon 4 of *Polg* was cloned into pX459, which expresses Cas9 nuclease. This plasmid was co-delivered to JM8A3 ESCs via electroporation with a single-stranded oligonucleotide donor (ssODN) that included the missense R292C (CGC>TGC) and an additional silent mutation (F290F; TTC>TTT) to reduce recutting of the targeted allele^74^. Following puromycin selection, resistant clones were screened for proper targeting and to exclude integration of the pX459 plasmid. Multiple clones were selected for expansion and karyotyping, and euploid clones were injected into C57BL/6J-albino blastocysts (B6(Cg)-Tyrc-2J/J; JAX STOCK# 000058) to produce chimeras. Chimeras were bred to C57BL/6NJ (JAX Stock #005304) mice and a single germline clone was selected for expansion and characterization. The line is available through the MMRRC repository (071656-JAX). All *Polg^R292C/R292C^* mice and littermate controls used for macrophage experiments were aged for 5-8 months. *Polg^R292C/R292C^*mice and littermate controls used for in vivo experiments were aged for 3-4 months. Breeding colonies of *Polg^R292C^* mice were maintained similarly to *Polg^D257A^* models. All animal experiments were approved by the Institutional Animal Care and Use Committees (IACUC) at Texas A&M University and The Jackson Laboratory.

### In vivo PAO1 infection

7-12 month old mice were moved into BSL2 procedure space and allowed to acclimate for 1-2 days. Mice were anesthetized with an intraperitoneal injection of 100-10 mg/kg Ketamine-Xylazine cocktail. Mice were then intubated with a 24G catheter. Sterile PBS (sham), 0.5 mg/kg LPS (E. Coli serotype 0111:B4; Invivogen), or the specified dose of PAO1-GFP was pipetted into the catheter in a 25-30 µL volume. A bolus of air was delivered using a syringe to ensure the dose evenly coated the lungs. Mice were kept warm until fully recovered from anesthesia and monitored daily until experimental timepoints were reached.

### Bronchial alveolar lavage fluid collection

Mice were euthanized using intraperitoneal injection of 200 mg/kg Pentobarbital Sodium to preserve the lung environment and structure of the tracheal. Upon confirmation of death the chest cavity and throat were opened. The trachea was canulated using a 23G needle threaded with tubing and the lung was washed with 3mL cold Hanks’ Balanced Salt Solution (HBSS, Thermo Fisher Scientific, 14175-129) containing 100 µM EDTA (VMR Life Science, E522) and used in subsequent flow cytometry and LegendPlex assays.

### Flow Cytometry

Bronchial alveolar lavage fluid (BAL) was spun at 350 g for 5 minutes to pellet the cells. Cells were transferred to a V bottom plate for staining. Cells were stained with viability dye then Fc receptors were blocked with anti-mouse CD16/CD32 (2.4G2) antibody. Surface proteins were stained with antibodies specified in Table 6. Cells were permeabilized with Foxp3/Transcription Factor Staining Buffer Kit (TNB-0607-KIT, Tonbo) and stained with antibodies against intracellular proteins. Flow cytometry analysis was done using a 5L Cytek Aurora.

### Colony forming unit counts from macrophages and tissues

BMDMs or tissues were homogenized in 0.1% NP40 in sterile PBS using an immersion homogenizer for 15 seconds per tissue. Serial dilutions were plated on soy agar containing 100 μg/mL carbenicillin. Plates were incubated at 37°C overnight and colonies were counted.

### Histology

Tissues were collected in cassettes, then were washed in PBS, incubated for 24 hours in 10% formalin, and transferred to 70% ethanol. Tissue embedding, sectioning, and staining were performed at AML Laboratories in Jacksonville, FL or at the histology core at Texas A&M University. Images of hematoxylin and eosin (H&E) staining were collected using an Olympus slide scanner with a 20X objective. Slides were analyzed and scored by a pathologist, Dr. Jessica Wong, at the Jackson Laboratory in Sacramento, CA. Briefly, semi-quantitative histologic scoring from 0 to 4 severity was performed on neutrophilic and mononuclear cell inflammation. 0 = no inflammation, 1 = minimal inflammation including individual cells or small clusters, 2 = mild inflammation including small clusters and increased perivascular and peribronchiolar cuffing, 3 = moderate inflammation including multifocal to coalescent areas and perivascular cuffing, and 4 = severe inflammation including coalescent to patchy-diffuse inflammation.

### Body Condition Scoring

Mice were monitored three times daily and were assigned a body condition score based on an additive score of appearance, behavior, hydration, and respiration. A score of 0 corresponds to a healthy mouse and a score of 10 corresponds to a moribund animal requiring euthanasia. Animals were given an appearance score of 0 if normal, 1 if they lacked grooming or ruffled fur, 2 if they displayed piloerection, and 3 if they displayed hunched posture. Animals were given a behavior score of 0 if they were socially active, 1 if they had decreased mobility, 2 if they showed increased isolation and only moved when prodded, and 10 if they did not ambulate with prodding. Animals were given a hydration score of 0 if the skin tent immediately returned to normal, 1 if the tent took 1-2 seconds to return to normal, 2 if the tent took greater than 2 seconds to return to normal, and 3 if this occurred accompanied by sunken and dry eyes. Animals were given a respiration score of 0 if they displayed a normal breathing pattern, 1 if they had an increased breathing rate with an abdominal component, 3 if their breathing appeared labored, and 10 if their breathing was agonal.

### Cell Culture

L929 cells were obtained from ATCC and maintained in DMEM (Sigma-Aldrich, D5796) and 10% fetal bovine serum (FBS; VWR, 97068-085). Primary bone marrow-derived macrophages (BMDMs) were isolated from femurs of 6-12 month old WT, *Polg^D257A^*and 2-6 month old *Polg^R292C^* mice as described^20^. Briefly, whole femurs were crushed in media and passed through a 70 µm filter. Red blood cells were lysed through osmosis by resuspending in ACK lysis buffer (Quality Biological, 118-156-101) for 2 minutes. Remaining cells were passed through a 40 µm filter, counted, and plated on non-tissue culture treated dishes. Macrophages were differentiated over for 7 days using 30% L929 conditioned medium. Peritoneal macrophages (PerMacs) were collected from the peritoneal cavity via gavage 4 days after intraperitoneal injection of 3% brewer thioglycolate medium (Sigma-Aldrich, B2551) and plated in in medium with 10% FBS. Unless stated, 6 x 10^5^ BMDMs and 1.2×10^6^ PerMacs per millimeter were used in in vitro experiments. Cells were plated and left to adhere overnight prior to challenge.

### *Pseudomonas aeruginosa* (PAO1) in vitro infection

GFP-expressing PAO1 was provided by Dr. Carolyn Cannon and was grown on Soy agar plates (BD, 211043) with 100 μg/mL carbenicillin (Dot Scientific, 46000-5) overnight from a glycerol stock. A single colony was incubated in 5 mL Mueller Hinton broth (BD, 275730) with carbenicillin at 37□C spinning overnight. Overnight culture was back diluted 1:5 and grown for 2 hours until bacteria reached log-phase. Bacteria were washed with sterile phosphate-buffered saline (PBS) and opsonized for 20 minutes in mouse serum (Fisher, PI31880). After two more washed with PBS macrophages were infected at the corresponding MOI and spun at 250 g for 5 minutes to synchronize infection. For inhibitor experiments, cells were primed with 10 µM Ac-YVAD-cmk (Invivogen, inh-yvad), 10 µM Q-VD-OPh (Millipore Sigma, 551476), or 10 µM MCC950 (InvivoGen inh-mcc) for 3 hours prior to challenge. For antibody blockade experiments, cells were primed with 20 µg/mL anti-IFNAR antibody (Bio X Cell, BE0241), anti-IL-1R (Bio X Cell, BE0256), or anti-IgG control (Bio X Cell, BE0083 or BE0091) antibodies for 4 hours prior to challenge. One hour after infection, wells were washed and treated with 300 μg/mL gentamicin (VWR, 0304) for 30 minutes to inactivate extracellular bacteria. The remainder of the experiment was completed with media supplemented with 50 μg/mL gentamicin and inhibitor or blocking antibody when described.

### *Escherichia coli* (E. coli) in vitro infection

E. Coli containing empty pcDNA3.1 vector for antibiotic resistance was grown on LB agar plates (Fisher, DF0401-17) with 100 μg/mL carbenicillin (Dot Scientific, 46000-5) overnight from a glycerol stock. A single colony was incubated in 5 mL LB broth (Fisher, BP1427-500) with carbenicillin at 37□C spinning overnight. Overnight culture was back diluted 1:5 and grown for 2 hours until bacteria reached log-phase. Bacteria were washed with sterile phosphate-buffered saline (PBS) and opsonized for 20 minutes in mouse serum. After two more washed with PBS macrophages were infected at the corresponding MOI and spun at 250 g for 5 minutes to synchronize infection. One hour after infection, wells were washed and treated with 300 μg/mL gentamicin (VWR, 0304) for 30 minutes to kill extracellular bacteria. The rest of the experiment was completed in DMEM + 10% FBS supplemented with 50 μg/mL gentamicin.

### Inflammasome challenge

BMDMs were primed for 4 hours with 20 ng/mL LPS-B5 Ultrapure (InvivoGen, tlrl-pb5lps). When used, caspase inhibitor Ac-YVAD-cmk (InvivoGen, inh-yvad) was added an hour after LPS, to allow for 3 hour priming, at 10 μM concentration. To trigger the non-canonical and NLRC4 inflammasomes respectively, polyethyleneimine (Alfa Aesar, 43896) was used to transfect 2 μg/mL LPS^75^ and 10 ng/mL PA flagella^76^ into the cytosol. To trigger the NLRP3 inflammasome, 5 μM ATP (Sigma, A6419) or 40 μM Nigericin (Cayman Chemical Company, 11437) was added to the media. When used, 20 µg/mL anti-IFNAR antibody (Bio X Cell, BE0241) or anti-IgG control (Bio X Cell, BE0083) were added to cells 5 hours prior to LPS priming and added again upon LPS transfection.

### Immunoblotting

Cells and tissues were lysed in 1% NP-40 lysis buffer supplemented with protease inhibitor and then centrifuged at 15,000 rpm for 10 min at 4 C and the supernatant was collected as the cellular lysate. A bicinchoninic acid assay (BCA) protein assay (Thermo Fisher Scientific, PI23235), was used to quantify protein and 10 to 40 μg total protein were loaded into 10% SDS-polyacrylamide gels and transferred onto 0.22 μM polyvinylidene difluoride membranes. Membranes were dried to return them to a hydrophobic state and then incubated in primary antibodies diluted in 1x PBS containing 1% casein overnight at 4 C rolling. Membranes were washed and incubated with horseradish peroxidase (HRP)-conjugated secondary antibodies for 1 hour at room temperature, shaking. Membranes were developed with Luminata Crescendo Western HRP Substrate (Millipore, WBWR0500). Primary antibody information can be found in Table 1.

**Table 1:** Western blot antibodies.

| Protein | Company | Catalog |
| --- | --- | --- |
| ASC | Cell Signaling Technology | 67824 |
| CASPASE-1 | Adipogen | AG-20B0042-C100 |
| CASPASE-11 | Cell Signaling Technology | 14340 |
| GAPDH | ProteinTech | 60004-1-Ig |
| GBP5 | Cell Signaling Technology | 67798 |
| GSDMD | Abcam | ab209845 |
| GSDME | Cell Signaling Technology | 88874S |
| HSP60 | Santa Cruz Biotechnology | Sc-1052 |
| IFIT3 | Ganes Sen, Cleveland Clinic | N/A |
| IL-1β | Cell Signaling Technology | 12242 |
| IRF1 | Cell Signaling Technology | 8478 |
| IRG1 | Cell Signaling Technology | 17805 |
| NLRC4 | Abcam | Ab201792 |
| NLRP3 | Cell Signaling Technology | 15101 |
| OXPHOS | Abcam | ab110413 |
| pSTAT1 | Cell Signaling Technology | 7649 |
| pTBK1 | Cell Signaling Technology | 5483 |
| pIkBα | Cell Signaling Technology | 2859 |
| TBK1 | Cell Signaling Technology | 3504 |
| STAT1 | Cell Signaling Technology | 9172 |
| ZBP1 | Adipogen | AG-20B-0010 |

**Table 2:** Immunofluorescence microscopy antibodies.

| Protein | Company | Catalog |
| --- | --- | --- |
| CASPASE-11 | Cell Signaling Technology | 14340 |
| GFP | Thermo Fisher Scientific | NC9777966 |
| LPS | Thermo Fisher Scientific | PA173178 |
| PAO1 | Thermo Fisher Scientific | PA173116 |
| GBP5 | Cell Signaling Technology | 67798 |
| TFAM | Proteintech | 23996-1-AP |
| PDH | Abcam | Ab110333 |
| DNA | Sigma Millipore | CBL186 |
| ASC | Cell Signaling Technology | 67824 |

### LegendPlex multianalyte cytokine analysis

Supernatants from macrophages were subjected to centrifugation prior to the assay to remove detached cells and debris from the sample. The assay was completed using the LegendPlex Mouse Inflammation Panel (BioLegend, 740446) according to the included protocol. Samples were analyzed by flow cytometry using a 5L Cytek Aurora and the BioLegend analysis software. Samples were normalized to total protein content of each sample determined by a bicinchoninic acid assay (BCA) protein assay and MFI from the blank control beads was subtracted to exclude background.

### Enzyme-linked immunosorbent assay (ELISA)

Supernatants were collected from macrophages plated on glass coverslips. Macrophages were stained for IF analysis while supernatants were subjected to centrifugation to remove detached cells and debris from the sample. Samples were stored at -80C until use in this assay and were thawed on ice upon use. The assay was completed in 96-well half area clear, flat bottom plates (USA Scientific, 5667-5061) and wells were incubated overnight at 4 C in capture antibodies. Blocking was completed for one hour with PBS containing 10% FBS at room temperature. Standards and samples were added to wells for two hours. All sample dilutions were made in blocking buffer. Next, wells were incubated in detection antibody for one hour. Avidin-HRP and trimethyl boron (TMB) were used for quick color development. Plates were washed with PBS containing 0.05% Tween 20 between each step. Capture and detection antibodies were used in the following pairs from BioLegend (Table 3). Samples were normalized to corresponding nuclei counts from 20X IF images taken on an EDLIPSE Ti2 (Nikon). Nuclei counts were determined using the automated bright spot counting function in NIS-Elements on DAPI and an average of 3 images was used to normalize ELISA. Normalization of BAL from PAO1-*algC* infected mice was normalized to protein concentration. ELISA concentrations were determined using an epoch plate reader (BioTek) and BioTek Gen5 software. Normalized results were graphed in GraphPad.

**Table 3:** ELISA antibodies (Biolegend)

| Protein | Catalog (Binding) | Catalog (Detection) |
| --- | --- | --- |
| TNFα | 506302 | 506312 |
| IL-1β | 503514 | 515801 |
| IL-6 | 504502 | 504602 |
| IL-1α | 503208 | 512503 |
| IFNβ | 519202 | 508105 |

### Quantitative polymerase chain reaction

Relative gene expression with measured by qRT-PCR in cells. Total cellular RNA was isolated using Quick-RNA microprep kit (Zymo Research, R1051). Approximately 0.5 µg of RNA was used to make cDNA with XLT qScript cDNA Synthesis Kit (Quantabio, 95047). cDNA was then used in a qPCR assay using PerfeCTa Green FastMix (Quantabio, 95071). Two to three biological replicates and two to three technical replicates were normalized to *Rpl37* cDNA using the 2^-ΔΔCT^ method. All primer sequences can be found in Table 4.

**Table 4:**
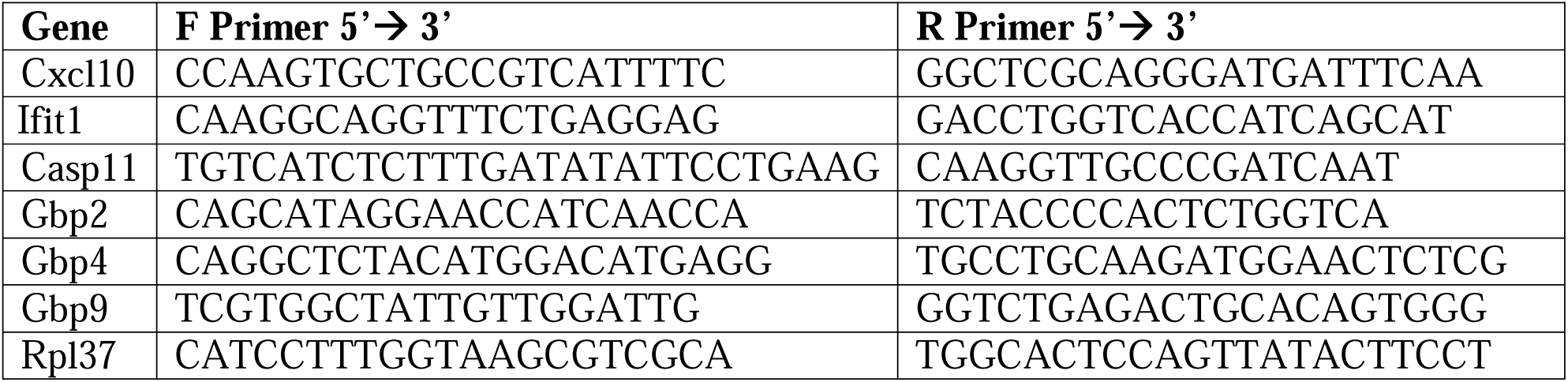
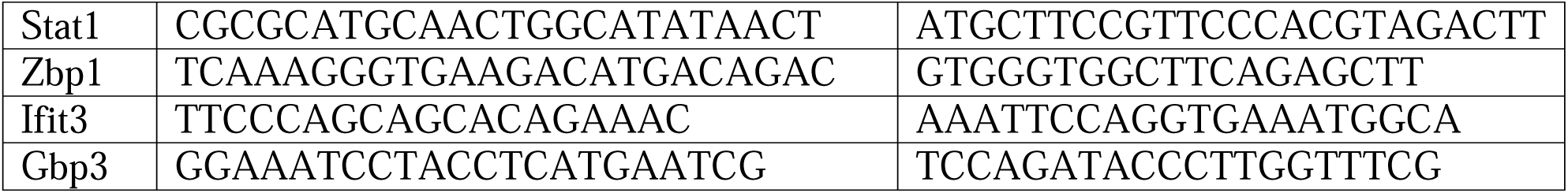
Primer sequences for qRT-PCR analysis.

### Immunofluorescence (IF) imaging

Cells were seeded at 2.5×10^5^ cell/well in a 24 well plate and allowed to grow overnight on glass coverslips sterilized with ethanol. After challenge or infection, cells were washed in PBS and fixed with 4% paraformaldehyde for 20 minutes. Next cells were permeabilized with 0.1% Triton X-100 in PBS for 5 minutes, blocked with PBS containing 5% FBS for 30 minutes, stained with primary antibodies (table 2) for 1 hour, stained with secondary antibodies for 1 hour, and stained with 200 ng/mL 4’,6-diamidino-2-phenylindole (DAPI) (ThermoFisher Scientific, 62247) for 2 minutes. When described, coverslips were stained with 2 µg/mL CellMask (ThermoFisher Scientific, C10046) for 10 minutes after secondary antibody staining and washed multiple times in PBS before DAPI staining. Coverslips were mounted with ProLong Diamond Antifade Mountant (Invitrogen, MP33025) and sealed with nail polish. Cells were imaged with a 40x or 60x oil-immersed objective with an ECLIPSE Ti2 widefield microscope (Nikon) and analyzed with the NIS-Elements AR 5.21.02 software.

Quantification was done from multiple intensity projection (MIP) images generated from Z-stacks. MIPs were made and analysis was done using NIS-Elements AR 5.21.02. Specifically, cell death was determined using the automated bright spot counting function in NIS-Elements on DAPI to count the number of nuclei. Dying cells were manually determined by identifying cells with condensed nuclei and low CellMask staining. Percentage of dying cells were determined and graphed in GraphPad.

Protein intensities were quantified by thresholding a mask over each channel and exporting sum intensities and area of masks. The sum intensities were normalized to area or cell numbers where described. When normalized to cell number, cells were counted using the automated bright spot counting function in NIS-Elements on DAPI. To quantify intensities of colocalized proteins, a mask was generated over each protein. Next, a new mask was made for the overlap of these channels and the intensity and area of each channel in this mask was exported and graphed.

To quantify nucleoids, whole cells were outlines and made into a region of interest (ROI) and these ROIs were numbered. A thresholding mask was made on the channels with TFAM and DNA staining and channel intensities were determined for these masks and graphed. A new mask was made based on the overlap of TFAM and DNA. The area of this mask and number of spots per ROI were exported and graphed in GraphPad. Graphs and statistics were generated in GraphPad. Huygens Essential software (v21.4.0) was used to deconvolve representative images from z-stacks taken with a 60X objective in oil immersion.

### Lactate dehydrogenase release

Lactate dehydrogenase (LDH) was measured in the supernatant of PAO1 challenged macrophages using a LDH-Cytox Assay Kit (Biolegend, 426401) according to manufacturer’s recommendations. A set of cells were treated with 1% Triton X-100 and the supernatant was used to determine total cellular LDH.

### PI uptake

BMDMs in 96 well half area glass bottom imaging plates were seeded at 6×10^4^ cells/well in 50 µL of Phenol-Red free DMEM + 10% FBS. The next day, cells were subjected to inflammasome challenge or infection with PAO1. Propidium iodide (PI, Alfa Aesar, J66584) was added to media and plates were immediately imaged every ten minutes using a BioTek Cytation V imager which maintains 37□C and 5% CO_2_. PI positive nuclei were identified and counted using BioTek Gen5 software. Control wells treated with PI and Triton X-100 (to permeabilize cells) to for normalization. The difference between PI positive cells compared to the initial reading were graphed in line graphs using GraphPad Prism.

### Cell fractionation and mitochondrial enrichment

Macrophages were lifted from cell culture dishes in PBS using a cell scraper. Cells were washed with PBS and then resuspended in Resuspension buffer (10 mM NaCl, 1.5 mM CaCl2, 10 mM Tris-Hcl, pH 7.5) with phosphatase and protease inhibitors. A dounce homogenizer was used to break cell membranes and liberate organelles. Mitochondria were bound to magnetic beads coated with TOMM20 antibodies isolated using a mitochondria isolation kit (Miltenyi Biotec, 120-009-492). The remaining cell components and unbound mitochondria were pelleted and run as the cellular fraction. All samples were lysed in 1% SDS lysis buffer and run on a western blot for analysis.

### Seahorse assay

Mitochondrial respiration and glycolysis in BMDMs were measured using a Seahorse XFe96 Analyzer. Macrophages were plates at a denisity of 5×10^5^ cells/well in DMEM supplemented with 10% FBS and 5% L929 in an Agilent Seahorse XF96 Cell Culture Microplate. When described, 20 ng/mL LPS was added 16 hours prior to analysis. Cells were placed in XF assay medium immediately before analysis. ECAR and OCR measurements were recorded through sequential addition of 10 mM glucose, 1.5 µM oligomycin, 1.5 µM carbonyl cyanide p-trifluoromethoxy phenylhydrazone with 1 mM sodium pyruvate, and finally 833 nM antimycin A plus 833 nM rotenone. Data was analyzed and calculations were made according to^77^.

### siRNA knock down in bone marrow derived macrophages

Bone marrow was isolated from mice as described above. On day four, macrophages were moved to 10 cm dishes. On day five, macrophages were transfected with 25 nM of Negative Control (51-01-14-03), mm.Ri.Casp4.13.1, mm.Ri.Gbp2.13.1, or mm.Ri.Gbp5.13.1 DsiRNAs (IDT) using Lipofectamine RNAiMAX (ThermoFisher Scientific, 13778100) in OPTIMEM (Gibco, 51985-034) according to company recommendations. Six hours after transfection, media was removed and replaced with fresh DMEM containing 10% FBS, 5% L929, and 20 µg/mL anti-IFNAR antibody (Bio X Cell, BE0241) to reduce IFN-I-mediated toxicity due to siRNA transfection. A day after transfection, cells were replated in cell culture plates in macrophage media without anti-IFNAR antibody, then used in infection experiments the next day as described above.

### mtDNA copy number analysis

DNA was isolated from cell pellets or crushed tissues by boiling for 30 minutes in 50 mM NaOH lysis buffer and then neutralizing using 1 M Tris, pH8 solution. DNA was diluted to 2 ng/uL and utilized in qRT-PCR reaction (described above) with mtDNA specific primers. Two to three biological replicates and two to three technical replicates were normalized to nuclear encoded *Tert* using the 2^-ΔΔCT^ method. All primer sequences can be found in Table 5.

**Table 5:**
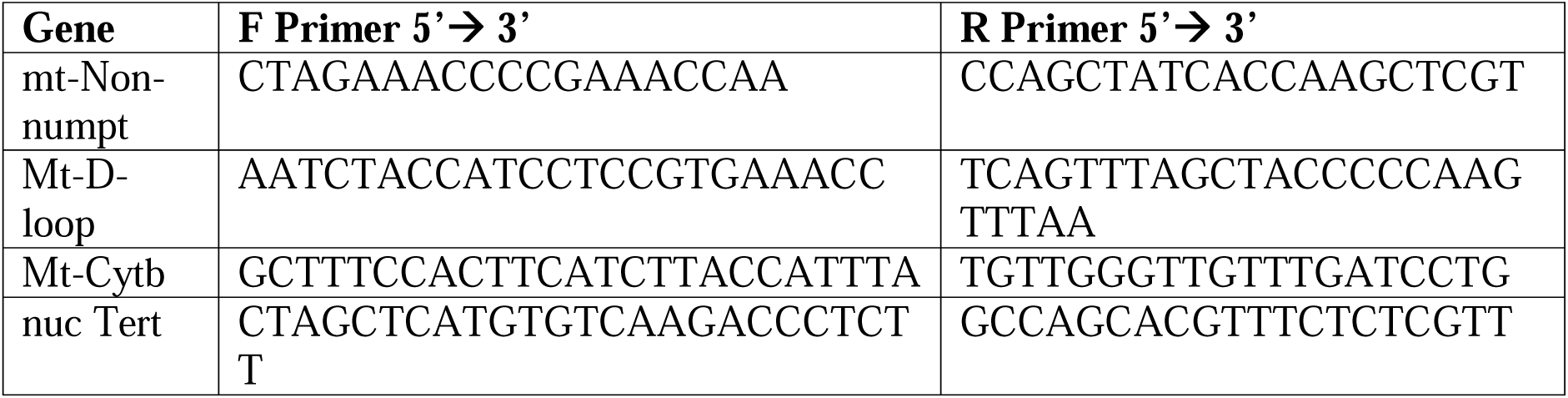
Primers for mtDNA copy number analysis by qPCR.

**Table 6:** Flow cytometry antibody panel.

| Antigen | Fluorophore | Laser | Company | Catalog/ Ref # |
| --- | --- | --- | --- | --- |
| CD45 | BUV395 | UV2 | BD | 564279 |
| Ly6C | BV650 | V11 | BioLegend | 128049 |
| MHCII | BV711 | V13 | BioLegend | 107643 |
| CD19 | BV605 | V10 | BioLegend | 115539 |
| CD3 | APC-fire810 | R8 | BioLegend | 100267 |
| F4/80 | VF450 | V3 | TONBO Biosciences | 75-4801-U025 |
| PA01 | FITC | B2 |  |  |
| CD11c | PE-Cy5 | YG5 | TONBO Biosciences | 55-0114-U100 |
| TNF | PE-Cy7 | YG9 | Biolegend | 506324 |
| Ly6G | PE | YG1 | TONBO Biosciences | 50-1276-U100 |
| Siglec-F | APC | R1 | Biolegend | 155508 |
| CD11b (M1/70) | APC-Cy7 | R7 | TONBO Biosciences | 25-0112-U100 |
| CD326 | PE/Dazzel 594 | YG3 | Biolegend | 118235 |
| Viability | Ghost Dye 710 | R4 | TONBO Biosciences | 13-0871-T100 |

### Transmission electron microscopy

Macrophages were lifted from cell culture dishes and pellets were fixed for 1 hour in 4% PFA. After fixation, cells were rinsed in 0.1M cacodylate buffer and then post-fixed in 1% osmium tetroxide (OsO_4_) in 0.1 M cacodylate buffer for 1 hour at room temperature. Cells were rinsed again in the same buffer then spun down into pellets with 2% liquid agarose in distilled water and placed in a 4 C refrigerator for 30 minutes, allowing the agarose to harden. After trimming the agarose cell pellets, samples were transferred through graded ethanol (40% ETOH to 100% ETOH), propylene oxide, and then infiltrated with 1:1 propylene oxide:Embed 812 resin (Electron Microscopy Sciences, Hatfield, PA) overnight, 1:3 propylene oxide:Embed 812 resin for 4 hours, and finally 100% Embed 812 resin overnight. Tissues were embedded in 100% resin in beam capsules and cured at 65°C for 48 hours prior to sectioning. Thin sections (90 nm) were cut on a Leica EM UC7 ultramicrotome (Leica Microsystems, Buffalo Grove, IL.) with a diamond knife, placed onto 300 mesh copper grids and stained using 2% uranyl acetate and Reynolds lead citrate. Samples were evaluated at 80 kV using a JEOL JM-1230 transmission electron microscope (JEOL, Tokyo, Japan) and images collected with an AMT Nanosprint15 MK-11 sCMOS digital camera (Advanced Microscopy Techniques, Woburn, MA.). Image quantification of cellular and PAO1 areas were completed in using NIS-Elements AR 5.21.02.

### RNA sequencing and bioinformatic analysis

Total cellular RNA was prepared from uninfected BMDMs and infected BMDMs using Quick-RNA microprep kit (Zymo Research, R1051) and used for the next-generation RNA-sequencing procedure at Texas A&M University Bioinformatics Core. BaseSpace Sequence Hub (Illumina) was used for RNA seq analysis. Briefly, Spliced Transcripts Alignment to a Reference (STAR) algorithm of RNA-seq Alignment V2.0.0 software was used to align the results to reference genome Mus musculus/mm10 (RefSeq), and then RNA-seq Differential Expression V1.0.0 software was used to obtain raw gene expression files and determine statistically significant changes in gene expression in *Polg* mutant macrophages relative to WT. Ingenuity Pathway Analysis software (QIAGEN) was used to identify gene families and putative upstream regulators in the datasets. Gene Set Enrichment Analysis (GSEA) Software (Broad Institute) was used to generate enrichment plots and examine pathway enrichment. Heatmaps were generated using GraphPad Prism. Datasets of LPS-treated *Polg^D257A^* BMDMs used in Fig. 1-2 and Supplementary Fig. 1 were previously reported^20^ and are available under Gene Expression Omnibus number GSE171960. Normalized gene counts from PAO1-infected *Polg^D257A^* BMDMs are supplied for review in a supplemental data spreadsheet (see **Dataset 1** Excel file). These data will be deposited into the Gene Expression Omnibus upon acceptance of the paper for publication.

RNA sequencing of lungs was performed by isolating total cellular RNA from infected lungs using a Direct-zol RNA miniprep Plus kit (Zymo Research, R2072). Next-generation RNA-sequencing was performed by The Jackson Laboratory Genomic Technologies Core. Partek Flow (v12.3.0) was used for RNA seq analysis. Briefly, Spliced Transcripts Alignment to a Reference (STAR) algorithm of RNA-seq Alignment was used to align the results to reference genome Mus musculus/mm10 (RefSeq), and then RNA-seq Differential Expression software analysis was used to obtain raw gene expression files and determine statistically significant changes in gene expression in *Polg* mutant macrophages relative to WT. Ingenuity Pathway Analysis software (QIAGEN) was used to identify gene families and putative upstream regulators in the datasets. Heatmaps were generated using GraphPad Prism. Normalized gene counts from PAO1-infected *Polg^D257A^* lungs and PAO1-infected *Polg^R292C^* lungs are supplied for review in a supplemental data spreadsheet (see **Dataset 2** Excel file). These data will be deposited into the Gene Expression Omnibus upon acceptance of the paper for publication.

### In vitro proteomic and bioinformatic analysis

#### Cell sample protein extraction and quantification

The adherent cells were washed on the six-well culture plate with PBS (+calcium and magnesium) three times. After washing the cells, 250 µL of 6M Urea in 50 mM HEPES, pH 8.0 was added to each well. Cells were scraped from the plate and the solution was transferred to a 2 mL microcentrifuge tube. Samples were then snap frozen in liquid nitrogen and transferred to the Mass Spectrometry and Protein Chemistry Service at The Jackson Laboratory for further processing. Each tube had a pre-chilled 5 mm stainless steel bead (QIAGEN) added and were lysed with the Tissue Lyser II (QIAGEN) for 2 minutes at 30 1/s. A magnet was used to remove the stainless steel beads and each sample was further lysed using a 1 mL syringe with a 23-gauge needle for 10 passes. Samples were then ice-waterbath sonicated at 37 Hz (100% power; Firsherbrand FB11207) for 5 minutes (30 seconds on, 30 seconds off for five cycles). The cell lysates were then centrifuged at 21,000 x g at 4°C for 15 minutes and the supernatant was transferred to a new 1.5 mL microcentrifuge tube. A small fraction of the sample was then used to create a 1:50 dilution (sample:50 mM HEPES) and a microBCA assay (Thermo) was run for protein quantification according to the manufacturer protocol.

For each of the quantified cellular protein lysates, 20 µg of protein was diluted in 50 mM HEPES, pH 8.0 to bring the urea concentration less than 2M for the trypsin digest. Each sample was then reduced with 10 mM dithiothreitol (final concentration) for 30 minutes at 37°C for 30 minutes in a ThermoMixer (500 rpm agitation). Samples were then cooled to room temperature and alkylated with 15 mM iodoacetamide for 20 minutes at 21oC in a ThermoMixer (500 rpm agitation). Sequence grade modified trypsin (Promega) was then add at a 1:40 ratio (trypsin:protein) to each sample for digestion at 37°C for 20 hours. Millipore C18 zip-tips (Millipore, ZTC18S096) were then used to desalt and purify the peptide digests according to the manufacturer protocol^78,79^. All purified peptide samples were then dried using a vacuum centrifuge. Dried peptide digests were reconstituted in 20 µL of 98% H2O/2% ACN with 0.1% formic acid via pipetting up and down 10 times, followed by vortexing at max speed for 30 seconds. Reconstituted samples were centrifuged at low speed for 15 seconds on a tabletop minicentrifuge to collect the liquid from the walls in the bottom of the microcentrifuge tube. All liquid was then transferred to a mass spec vial (Thermo) and placed in the Vanquish Neo (Thermo) liquid chromatography autosampler set to 4oC.

#### Liquid Chromatography Mass Spectrometry Analysis

Data independent acquisition mass spectrometry analysis (DIA-MS) was performed on a Thermo Orbitrap Astral mass spectrometer coupled to the Vanquish Neo liquid chromatography system with an EASY-Spray column (PepMap Neo C18, 75 µm x 150 mm, #ES75150PN) in the Mass Spectrometry and Protein Chemistry Service at The Jackson Laboratory. Each sample had two technical replicate runs consisting of an 8 µL injection of reconstituted sample. Before each sample injection the separation column underwent fast equilibration in the method using pressure control at 1500 bar and an equilibration factor of three column volumes. The trapping column underwent fast wash and equilibration using a was factor of 100, an automatic equilibration factor, and flow control at 120 uL/minute. The method gradient was 25 minutes after the column equilibration using Buffer A (100% H2O with 0.1% formic acid) and Buffer B (100% acetonitrile with 0.1% formic acid) with a flow rate of 600 nL/min. The full gradient consisted of 2% B to 4% B from 0-2.5 minutes, 4% B to 6% B from 2.5-4.5 minutes, 6% B to 30% B from 4.5-22.5 minutes, and 30% B to 90% B from 22.5-24 minutes while ramping to 1 µL/minute flow rate in the last step. The column was then washed with 90% B at 1 µL/minute from 24-25 minutes. Orbitrap Astral settings included a static spray voltage using the NSI source at 2000 V in positive mode, an ion transfer tube temperature of 325oC, an expected peak width of 10 seconds, advanced peak determination, a default charge state of 2, EASY-IC lock mass correction, and RunStart mode. Full scan settings in positive mode included an orbitrap resolution of 240,000, scan range of 380-980 m/z, RF lens at 60%, normalized AGC target of 500% with an absolute AGC value of 5e6, and a maximum injection time of 5 ms. DIA method settings included a precursor mass range of 380-980 m/z, an isolation window of 2 m/z with no overlap, normalized HCD collision energy of 25%, detector type set to Astral with a scan range of 150-2000 m/z, RF lens at 60%, normalized AGC target of 500% with an absolute AGC value of 5e4, maximum injection time of 5 ms, and loop control set at 0.6 seconds.

#### DIA-MS data analysis

All Orbitrap Astral RAW data files underwent automatic processing through the Thermo Ardia Server and were directly sent to the Proteome Discoverer 3.1 workstation connected to the system. The RAW data files for each sample were searched against the UniProtKB Mus musculus (sp_tr_incl_isoforms TaxID=10090; v2024-05-01) protein database in Proteome Discoverer (version 3.1) using the CHIMERYS Ardia Server searching algorithm with the inderys 3.0 fragmentation prediction model. Search settings included trypsin digestion, a maximum missed cleavage of one, peptide length of 7-30 amino acids, peptide charge from 1-4, a fragment mass tolerance of 10 ppm, a dynamic oxidation of methionine modification (+15.995 Da), a static carbamidomethlyl modification of cysteine (+57.021 Da), and a false discovery rate of less than 0.05 for all matches; all other settings were set to manufacturer default recommendations. Total peptide amount was then utilized to normalize the ion signal in the samples. Quantification analyses settings included the use of unique peptides for quantification and a protein abundance-based ratio calculation was used for the summed abundances, followed by a background based t-test. Strict parsimony principle was then applied for protein grouping and master proteins were reported. The data was also run with no imputation and low abundance resampling imputation for comparison during the analysis process. The full dataset can be found in the Proteome Exchange PRIDE upload XXXXXXXXXX (number will go here when published, see **Dataset 3** Excel file).

### Statistical Analyses

To ensure rigor and reproducibility, experiments were repeated at least three times with multiple biological replicates. Error bars displayed throughout the manuscript represent the mean +/- s.e.m. Statistical significance between 2 groups was determined via unpaired, two-tailed Student’s t-test or Mann-Whitney test, and differences between 3 or more groups was determined by two-way ANOVA with appropriate post-test. Survival plots were compared by the Log-rank test. Sample sizes were chosen by started methods to ensure adequate power, and no randomization or blinding was used for animal studies. Significance was established as * p<0.05; ** p<0.01; *** p<0.001; **** p<0.0001; or not significant (n.s.).

